# High-resolution, high-throughput screening reveals potent target sites for RNAi and Cas13 systems

**DOI:** 10.1101/2025.11.17.688664

**Authors:** Jonathan Kong, Summerloretta S. K. Leung, Guojia Wu, Chi Fai Lau, Bobo Wing-Yee Mok, Mingxi Weng, Ralf Jauch, Honglin Chen, S. Chul Kwon

**Author notes:** These authors contributed equally to this work (Jonathan Kong, Summerloretta S. K. Leung).

## Abstract

RNA interference (RNAi) and CRISPR-Cas13 are widely used for targeted RNA degradation, yet identifying optimal target sites remains a major bottleneck. Furthermore, it remains unclear whether these mechanistically distinct platforms share target sites because high-resolution comparative RNA knockdown data are lacking. To address this, we developed a high-throughput screening platform utilizing shRNAone, an engineered scaffold that generates highly homogeneous small RNAs. After demonstrating the scaffold’s precision at a large scale, we applied high-resolution tiling screens to identify highly effective target sites in therapeutically relevant TTR and PCSK9 mRNAs, uncovering “clustered potent sites” within their 3′ UTRs. Importantly, the shRNAone platform successfully ranked clinically approved siRNAs—patisiran and inclisiran—among the most potent candidates. Further, by mapping CasRx and PspCas13b silencing landscapes across the TTR mRNA, we demonstrated that the clustered potent sites are largely shared between systems and are associated with local RNA accessibility. Additionally, while shRNAone and CasRx activities were largely restricted to the 3′ UTR, PspCas13b efficiently targeted both the coding sequence and the 3′ UTR. Together, this work reveals the existence of shared potent target sites between RNAi and Cas13 and provides a screening platform that can reduce the time and cost of developing RNA therapeutics.

**SIGNIFICANCE:** siRNAs are approved drugs that degrade disease-associated mRNAs, yet target-site selection relies largely on trial and error. Directly testing candidate sites via siRNA synthesis and knockdown measurement scales poorly. Conventional shRNA libraries are risky to interpret due to imprecise processing, and screening low-abundance secretory proteins has remained difficult to establish. We engineered an shRNA scaffold that produces highly homogeneous small RNAs and paired it with a multi-copy knock-in reporter system capable of screening such low-abundance targets. Comparative screening with two Cas13 nucleases showed that all three systems largely share the potent sites, suggesting these sites are platform-independent. The screen also recovered the target sites of clinically approved siRNA drugs, demonstrating its potential to shorten siRNA drug development.

## INTRODUCTION

RNA-targeting technologies have broadened their utility from widely used laboratory tools to encompass clinical therapies. RNA interference (RNAi) forms the therapeutic basis for approved drugs treating transthyretin amyloidosis^1^ and hypercholesterolaemia^2^. The RNA-targeting CRISPR systems, such as Cas13a^3^, Cas13b^4^, and Cas13d^5^, provide another biochemically distinct method for programmable transcript degradation in mammalian cells. Both platforms rely on a nuclease effector directed by a guide RNA to recognize specific target sequences. Although extensive RNA knockdown data have accumulated for each platform, it remains unclear whether they share fundamental rules for target selection. Because Argonaute 2 (AGO2) and Cas13 employ different RNA cleavage mechanisms and utilize guide RNAs of distinct lengths, it is unknown whether target-site selection rules can be transferred between these divergent systems.

Short hairpin RNAs (shRNAs) have been extensively employed for pooled genetic screening in human cells. The interpretation of these screens relies on the assumption that the measured phenotype corresponds precisely to the designed guide sequence. However, DICER does not always process hairpin substrates into a single uniform product because multiple factors, such as loop position, local sequence context, and duplex structure, dictate the exact cleavage site^6–9^. Consequently, a single designed hairpin can generate a population of small RNA isoforms with guide sequences shifted by one or two nucleotides^10–13^. This heterogeneity can be tolerated in standard low-resolution applications such as gene-level loss-of-function screens because misprocessed species also target the same mRNA. However, this processing variation becomes a critical limitation when the objective shifts from identifying a target gene to selecting a specific target site at nucleotide-level resolution.

Several engineering strategies have sought to generate homogeneous small RNAs from shRNA^6,14–16^. Kay and colleagues adjusted the apical loop position in RNA Pol III-expressed shRNAs to improve cleavage precision^6^, while Bartel and colleagues designed artificial primary microRNAs (pri-miRNAs) based on DROSHA processing rules^14^. However, these alternative shRNA designs lack large-scale validation of guide homogeneity across hundreds of distinct sequences and remain little used relative to miR-30-based backbones. Consequently, the miR-30-based shRNAmir remains the standard scaffold owing to its extensive validation across numerous labs and diverse loss-of-function screening applications^17–20^.

An shRNA system capable of producing homogeneous small RNAs can address additional unresolved challenges in small RNA biology and therapeutic design. First, it enables high-throughput evaluation of how seed and supplementary pairings contribute differently to the phenotypic activity of hundreds of microRNAs (miRNAs). Although seed pairing is firmly established as the primary driver of target recognition, the role of 3′-supplementary pairing remains incompletely resolved. Estimates of its functional contribution vary drastically depending on whether individual or hundreds of miRNAs are analyzed^21–23^, whether the experimental context is cell-free or cell-based^24–28^, and whether the measured output is biochemical binding or functional repression^25,27–29^. Expressing precise miRNA variants from a common shRNA scaffold in a pooled screen can provide a controlled environment to isolate the functional impact of non-seed sequences at an unprecedented scale *in cellulo*. Second, a precision shRNA platform bridges the historical gap between pooled shRNA screening data and synthetic siRNA target selection. Because therapeutic siRNAs require sequence precision at single-nucleotide resolution, processing heterogeneity has previously prevented the direct transfer of potent guide sequences from shRNA screens to siRNA designs. An shRNA scaffold that enables uniform guide biogenesis would therefore directly accelerate the discovery and validation of highly active siRNA target sites.

Even within RNA-targeting CRISPR systems, predicting target-site selection across diverse Cas13 families remains a significant challenge. Representatives from the Cas13b and Cas13d families, notably PspCas13b and CasRx (also known as RfxCas13d), demonstrate robust RNA knockdown in human cells^5,30,31^. While several tiling screens have generated guide design algorithms for individual Cas13 systems, it remains unclear whether an algorithm trained on one system can accurately predict the activity of another. For example, PspCas13b and CasRx utilize distinct structural architectures, including opposite orientations of the direct repeat relative to the spacer sequence, as well as differing guide lengths. This uncertainty persists because single-effector screens cannot distinguish whether identified efficacy determinants are intrinsic to the specific nuclease or fundamentally dictated by the target RNA. Separating these variables requires mapping mechanistically distinct systems across a common transcript at identical resolution.

Here, we therefore aim to address the methodological limitation shared by these RNAi and Cas13 systems. We develop shRNAone, an shRNA backbone engineered for uniform processing to yield homogeneous small RNAs that correspond precisely to the designed sequence. As proof of concept for this small RNA precision, we dissect the phenotypic contributions of seed and non-seed pairings using hundreds of human miRNA sequences. We then demonstrate that highly potent guide RNAs identified through pooled shRNAone screening can be directly translated into synthetic siRNA designs. Finally, we compare the single-nucleotide resolution knockdown profiles of shRNAone, PspCas13b, and CasRx on the same mRNA and reveal “clustered potent sites” shared by these distinct RNA-targeting systems. These findings suggest that exploiting these accessible regions may serve as a highly effective, platform-independent strategy for the rational design of next-generation RNA therapeutics.

## RESULTS

### An optimized shRNA scaffold for precise guide production

To yield a uniform population of small RNAs, an shRNA construct must undergo precise processing and loading. However, our initial evaluation of the two most widely used shRNA backbones, pLKO (The RNAi Consortium, TRC)^32^ and shRNAmir^33^, revealed that neither system satisfies this requirement. Sequencing of small RNAs generated from the pLKO scaffold across three distinct guide sequences demonstrated high frequencies of heterogeneous 5′ ends, possibly due to inconsistent RNA Pol III termination^34–36^ (Supplementary Fig. 1, 2). Similarly, the RNA Pol II-driven shRNAmir produced heterogenous guide RNAs (Supplementary Fig. 1, 2). Because these processing inconsistencies preclude target selection at nucleotide-level resolution, we sought to engineer an optimized scaffold.

We first tested the guide-strand position and length, and the mismatches in the upper stem, in five pri-miRNAs whose miRBase profiles indicate homogeneous production of their mature miRNAs (Supplementary Fig. 1, 2). After identifying the backbone that yielded the most homogeneous small RNAs, we further varied the lower and upper stem structures (Supplementary Fig. 3). We then compared four new designs with three previous designs^15,16,33^, using 144 shRNA sequences per design (Fig. 1a–e, Supplementary Fig. 4, 5). The best design, based on pri-miR-26a-2, produced the intended guide RNA in 90.8% ± 4.7% of all small RNA reads (mean ± s.d.) and was termed shRNAone (Fig. 1e, f). Under the same conditions, the standard shRNAmir design (pGIPZ) produced 74.0% ± 9.9% of all small RNA reads (Fig. 1d, f).

**Fig. 1:**
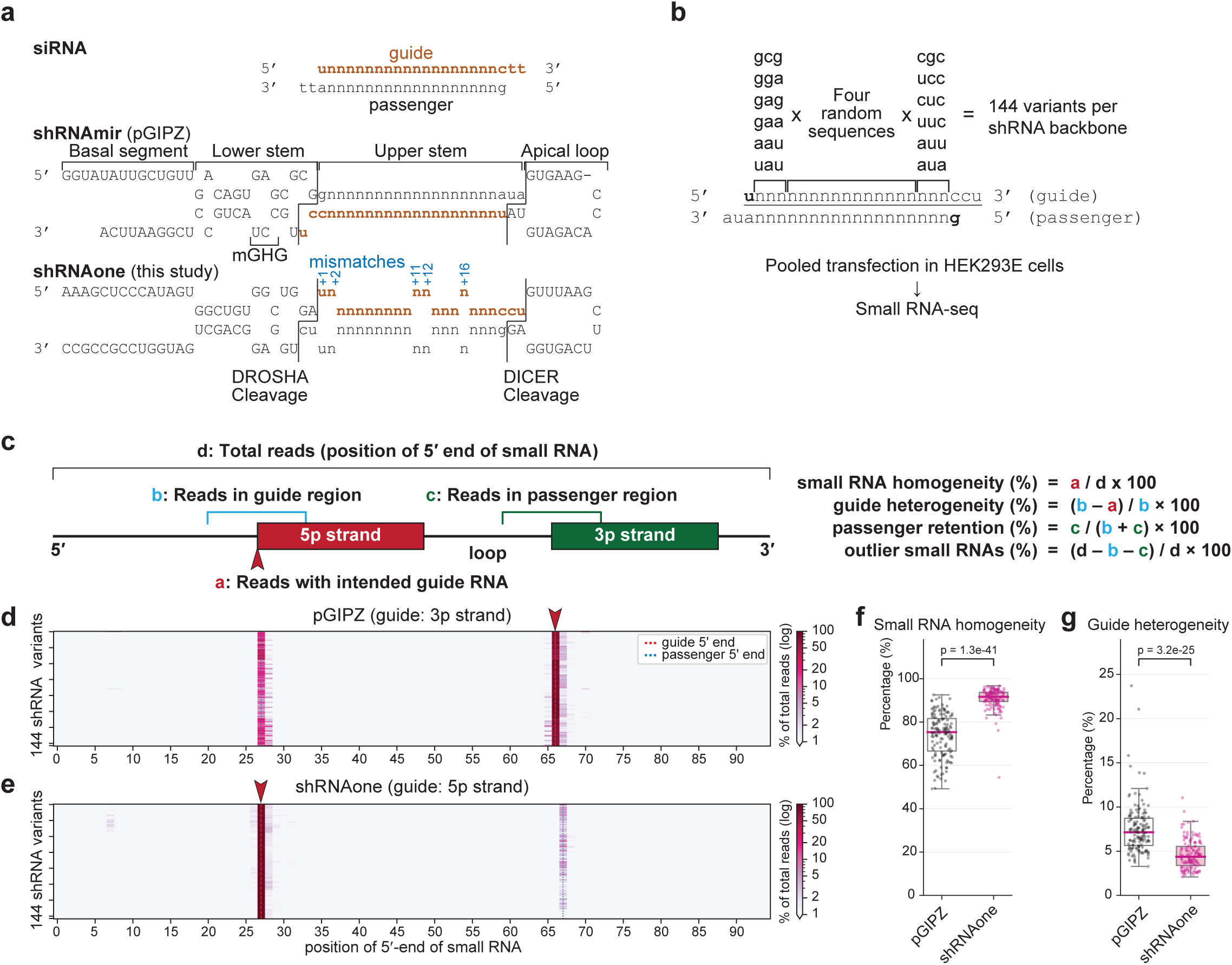
An optimized shRNA scaffold for precise guide production. **a** shRNAone compared with small interfering RNA (siRNA) and shRNAmir (pGIPZ). Red, guide strand; black, passenger strand and invariant backbone; n, user-definable positions; vertical bars, DROSHA and DICER cleavage sites. Guide positions are numbered from the DROSHA site; Mismatched positions in shRNAone (+1, +2, +11, +12, +16) are shown. **b** Benchmarking shRNA library design. A matrix of 144 inserts (4 random middle sequences × 6 guide 5′ triplets [gcg, gga, gag, gaa, aau, uau] × 6 guide 3′ triplets [cgc, ucc, cuc, uuc, auu, aua]) was cloned into each backbones, transfected into HEK293E cells, and profiled by small RNA-seq. **c** Definition of the four metrics. a (red), reads with the intended guide 5′ end; b (blue), all reads in the guide region; c (green), all reads in the passenger region; d, all reads mapping to the construct. **d**, **e** Heatmap of small-RNA 5′ end positions for the 144 shRNA variants expressed from pGIPZ (d; 3p guide) and shRNAone (e; 5p guide). Each row is one variant and each column is a 5′ end position of the mapped reads. Color indicates the percentage of a variant’s total reads starting at that position (each row sums to 100%) on a log scale from 1% to 100%. Dashed red and dotted blue vertical lines mark the intended guide and passenger 5′ ends, respectively. **f**, **g** Small RNA homogeneity (f) and guide heterogeneity (g). Each dot is one sequence variant; horizontal line and whiskers show mean ± sd. The P value was calculated using the Mann-Whitney U test (n = 144). See also Supplementary Fig. 5b.

Next, we analyzed the source of the remaining heterogeneity. All 5p-guide designs showed less guide heterogeneity than 3p-guide designs such as pGIPZ, suggesting that imprecise DICER cleavage increases guide heterogeneity (Fig. 1g, Supplementary Fig. 5). Previous shRNA designs, including pGIPZ, showed many reads from the passenger strand, likely due to inefficient passenger-strand ejection compared to shRNAone (Supplementary Fig. 5). Some designs in which we modified the mGHG motif, which is important for precise DROSHA processing^14,37^, produced undesirable small RNAs from lower stem regions (Supplementary Fig. 5, shRNAmir_GHG^miR-16^). We conclude that shRNAone produces highly homogeneous small RNAs.

We next evaluated its functional efficacy relative to the standard shRNAmir design (pGIPZ). Although both platforms achieved comparable on-target transcript knockdown in wild-type HEK293 cells (Supplementary Fig. 7a, b), they exhibited distinct dependencies on AGO2. In *AGO2*-knockout cell lines, shRNAmir activity was largely abrogated, underscoring its reliance on AGO2-mediated slicing activity for passenger strand removal. Conversely, shRNAone retained silencing activity in all three *AGO2*-knockout clones, significantly outperforming shRNAmir for five of the seven guide sequences tested, while the two backbones performed comparably for the remaining two guides (Supplementary Fig. 7c, d). Taken together, shRNAone not only provides superior processing uniformity but also faithfully engages RNA knockdown comparable to the standard shRNAmir scaffold.

### Precision small RNAs dissect miRNA pairing rules *in cellulo*

Since shRNAone produces highly homogeneous small RNAs, we sought to demonstrate its precision by dissecting miRNA pairing rules, which strictly require single-nucleotide accuracy. In the canonical model, mammalian miRNA target recognition is driven predominantly by the base pairing between the seed region (2–8 nt) and the target RNA (Fig. 2a)^38–40^. The 3′-supplementary pairing (around 13–16 nt) can provide additional binding affinity (Fig. 2a)^38–40^. Structural studies and *in vitro* binding assays showed that the supplementary pairing can increase target affinity by over twenty-fold in some miRNAs^21,41–43^. *In vivo* UV crosslinking studies, such as AGO CLIP, CLASH, and CLEAR-CLIP, also captured the miRNA–target interactions with the supplementary pairings^25,27,44,45^. However, the phenotypic contribution of supplementary pairings *in cellulo* has only been evaluated for a few miRNAs rather than hundreds of them^25,27,38,46^.

**Fig. 2:**
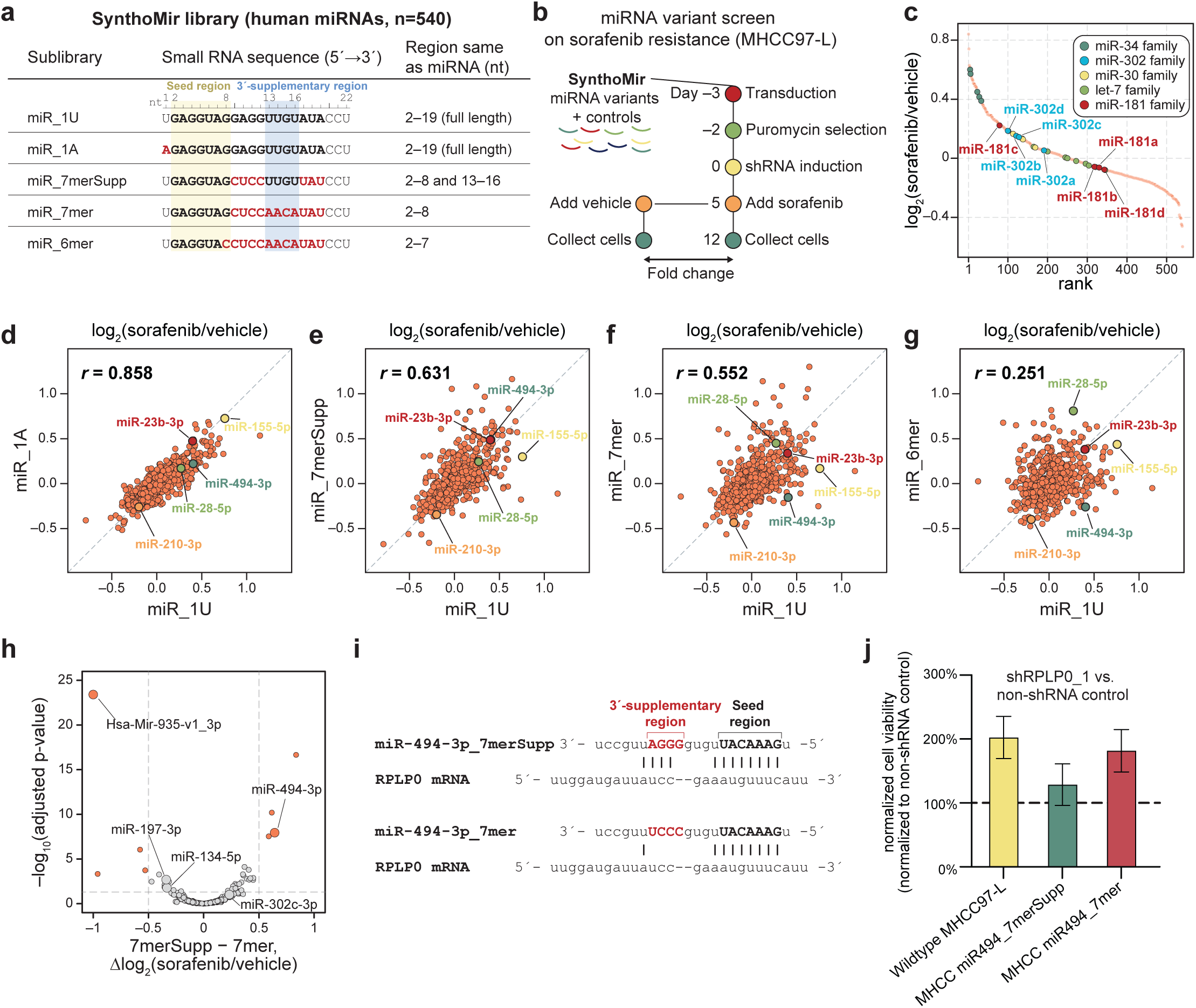
Precision small RNAs dissect miRNA pairing rules *in cellulo*. **a** SynthoMir library composition: five variants for each of 540 human miRNAs (2,700 total), shown for one miRNA. Light type, backbone-fixed 5′ uridine and 3′ CCU; 2–19 nt are customizable. Black, identical to the natural miRNA; red, complementary base (“base-flipped”); yellow and blue shading, seed and 3′-supplementary regions. Final column, miRNA-derived positions retained per sublibrary. **b** Pooled sorafenib-resistance screen in MHCC97-L cells transduced with the SynthoMir library. **c** The 540 miRNAs ranked by the mean log2FC of their miR_1U and miR_1A variants; five families are colored as indicated. **d–g** log2FC of the miR_1U variant against that of the miR_1A (d), miR_7merSupp (e), miR_7mer (f), and miR_6mer variant (g) of the same miRNA. Each orange circle is one miRNA (n = 540). Pearson’s correlation coefficients are shown. **h** Volcano plots comparing the sorafenib response of the 7merSupp design with 7mer. x, difference in log2(sorafenib/vehicle) between the two samples (Δlog2(sorafenib/vehicle)); y, −log10(adjusted P value), Benjamini–Hochberg. Dashed lines, |Δlog2(sorafenib/vehicle)| = 0.5 and P = 0.05; orange, miRNAs passing both cut-offs; grey, all others. Enlarged points denote the labeled miRNAs and carry no quantitative meaning. **i** Predicted duplexes between the miR-494-3p 7merSupp (top) and 7mer (bottom) variants and the RPLP0 mRNA. Red, supplementary region; bold, seed region; vertical bars, Watson–Crick pairs. **j** Normalized cell viability upon RPLP0 knockdown (shRPLP0_1) in WT cells (yellow) or cells expressing miR-494_7merSupp (green) or miR-494_7mer (red). Cell viability is normalized to the corresponding non-shRNA control (dashed line, 100%). Bars, mean ± s.d.; n = 3 biological replicates. See also Supplementary Fig. 14f.

To this end, we designed miRNA variants for all human miRNAs to decouple seed and supplementary contributions *in cellulo* and named the library SynthoMir (Fig. 2a, Supplementary Fig. 8a, Supplementary Table 4a). To build the library, we collected all human miRNAs from MirGeneDB 2.1 (540 unique sequences across 2–19 nt; Supplementary Table 5)^47^ and cloned each as five distinct variants: the native guide sequence starting with U (miR_1U); a 5′ U-to-A substitution (miR_1A); and three variants designed to isolate the contributions of different seed types and 3′-supplementary pairings (miR_7merSupp, miR_7mer, and miR_6mer) (Fig. 2a). The miR_1A control is based on the fact that the AGO2 MID pocket buries guide position 1, preventing it from interacting with the target RNA^48^. Because the AGO2 MID specificity loop readily accommodates both UMP and AMP (Kd = 0.12 and 0.26 mM, respectively) while discriminating against CMP and GMP (3.3–3.6 mM)^49^, the 5′ U and A variants should function similarly. Combined with 100 non-targeting controls, these five sets comprise the SynthoMir library of 2,800 constructs.

To evaluate the biochemical properties of miRNA variants using a phenotype-based approach, we selected a sorafenib resistance model in hepatocellular carcinoma (MHCC97-L) cells^50,51^. We first assessed whether the SynthoMir library faithfully recapitulates established miRNA targeting behaviors in this system (Fig. 2b, Supplementary Fig. 8b, c, Supplementary Table 4b, c). Amplicon sequencing of sorafenib- and vehicle-treated populations revealed highly similar enrichment among the individual members belonging to the miR-30, miR-34, miR-302, and let-7 families (Fig. 2c, Supplementary Fig. 9a–d). This family-level consistency aligns with the expectation that the seed sequence dictates primary target selection^40^. However, the miR-181 family presented an exception to this trend, as miR-181c was enriched by 22% relative to its paralogs, miR-181a, miR-181b, and miR-181d (Fig. 2c). Because miR-181c shares an identical seed with these paralogs but harbors a single-nucleotide deletion in its central region that shifts all downstream nucleotides, its divergent behavior cannot be attributed to seed identity (Fig. 2c, Supplementary Fig. 9e). Instead, this divergence directly implicates non-seed pairing as a determinant of target selection and functional outcome.

To examine this relationship systematically, we compared enrichment across the variant libraries. Phenotypic scores (expressed by log_2_[sorafenib/vehicle]) for miR_1U and miR_1A constructs correlated closely (n = 540, Pearson’s *r* = 0.858, *P* = 1.15 × 10^−1^^17^), confirming that 5′ U and 5′ A mimics function interchangeably (Fig. 2d). Relative to this reference point, correlation with the native miR_1U sequence declined progressively as sequence equivalence was reduced (miR_7merSupp, *r* = 0.631, *P* = 2.75 × 10^−61^; miR_7mer, *r* = 0.552, *P* = 1.88 × 10^−44^; miR_6mer, *r* = 0.251, *P* = 3.58 × 10^−9^) (Fig. 2e–g). Because the miR_7merSupp and miR_7mer designs preserve the identical sequence at positions 2 through 8 relative to miR_1U, they maintain the capacity to form 8mer and 7mer-m8 seed pairings depending on the specific target RNA context. Similarly, the miR_6mer design restricts interactions to 7mer-A1 and 6mer seed pairings. The correlation analysis indicates that while 8mer and 7mer-m8 seed pairings largely recapitulate full-length miRNA function, 7mer-A1 and 6mer seed pairings generally fail to induce meaningful target repression in cells, as reflected in the reduced correlation between miR_1U and miR_6mer. This hierarchical activity aligns closely with established seed-target binding affinities derived from high-throughput *in vitro* studies (8mer > 7mer-m8 > 7mer-A1 > 6mer)^39,52^.

### SynthoMir facilitates the identification of functional miRNA targets that depend on non-seed regions

To identify miRNAs whose activity depends on pairing outside the seed, we compared miR_7merSupp and miR_7mer designs that share identical seed pairing and differ in the 3′ supplementary region (positions 13 to 16) (Fig. 2h, Supplementary Fig. 10a, b, Supplementary Table 4d–g). As a proof of concept, we focused on miR-494-3p, which showed distinct phenotypic effects between these two libraries (Fig. 2h, Supplementary Fig. 10c). We performed RNA-seq (overexpressed miR-494_7merSupp vs. miR-494_7mer), an shRNA library screen (targeting differentially expressed genes from RNA-seq), and *in silico* seed sequence analysis (Supplementary Fig. 10d, 11–13, 14a–c, Supplementary Table 4i, j, 6, 7a–h). As a result, we identified *RPLP0* as the target gene corresponding to miR-494-mediated phenotypic changes (Fig. 2i). We validated the miR-494 target site on *RPLP0* mRNA using qRT-PCR and a luciferase assay (Supplementary Fig. 14d, e). Further, we confirmed that miR-494 and *RPLP0* are in the same regulatory axis by knocking down *RPLP0* in cells expressing either miR-494_7merSupp or miR-494_7mer (Fig. 2j, Supplementary Fig. 14f, g).

Because the RPLP0 transcript harbors a canonical 8-mer seed match for miR-494 (Fig. 2i), these data confirm that seed pairing alone is sometimes insufficient and that 3′-supplementary pairing is required for specific target repression in human cells. Taken together, we demonstrate that the SynthoMir library can facilitate the identification of functional miRNA targets that strictly depend on non-seed regions, and that shRNAone can dissect regulation requiring single-nucleotide precision.

### Enhanced split-GFP screening strategy for low-abundance secretory proteins

After confirming the homogeneity and functionality of the shRNAone scaffold, we utilized this property to test whether tiling shRNAs across endogenous transcripts could identify potent siRNA target sites for therapeutics development. We selected *AGT*, *TTR*, and *PCSK9* as they are targets of approved siRNA therapeutics, alongside *APOA1*. We first prepared parental cells by stably expressing split GFP_1-10_ or mNeonGreen2_1-10_ (mNG2_1-10_) in MIHA cells, an immortalized human hepatocyte cell line (Fig. 3a, Supplementary Fig. 15a)^53^. These large fragments were fused to an N-terminal signal peptide (SigP) and a C-terminal KDEL to ensure proper localization and maximum protein retention within the endoplasmic reticulum (ER)^54^. We then knocked in single or multiple copies of the corresponding small fragments, GFP_11_ or mNG2_11_, along with a C-terminal KDEL, in-frame at the 3′ end of each endogenous coding sequence (CDS). (Supplementary Fig. 15b). As previously reported^53^, multimerized small fragment knock-ins amplify the fluorescence intensity of endogenous proteins (Fig. 3b, Supplementary Fig. 15c–f). This enhanced signal allowed us to successfully separate the knockdown populations from control populations in both *TTR* and *PCSK9* knock-in cells, thereby enabling robust FACS-based screening (Fig. 3c, d, Supplementary Fig. 16a–d).

**Fig. 3:**
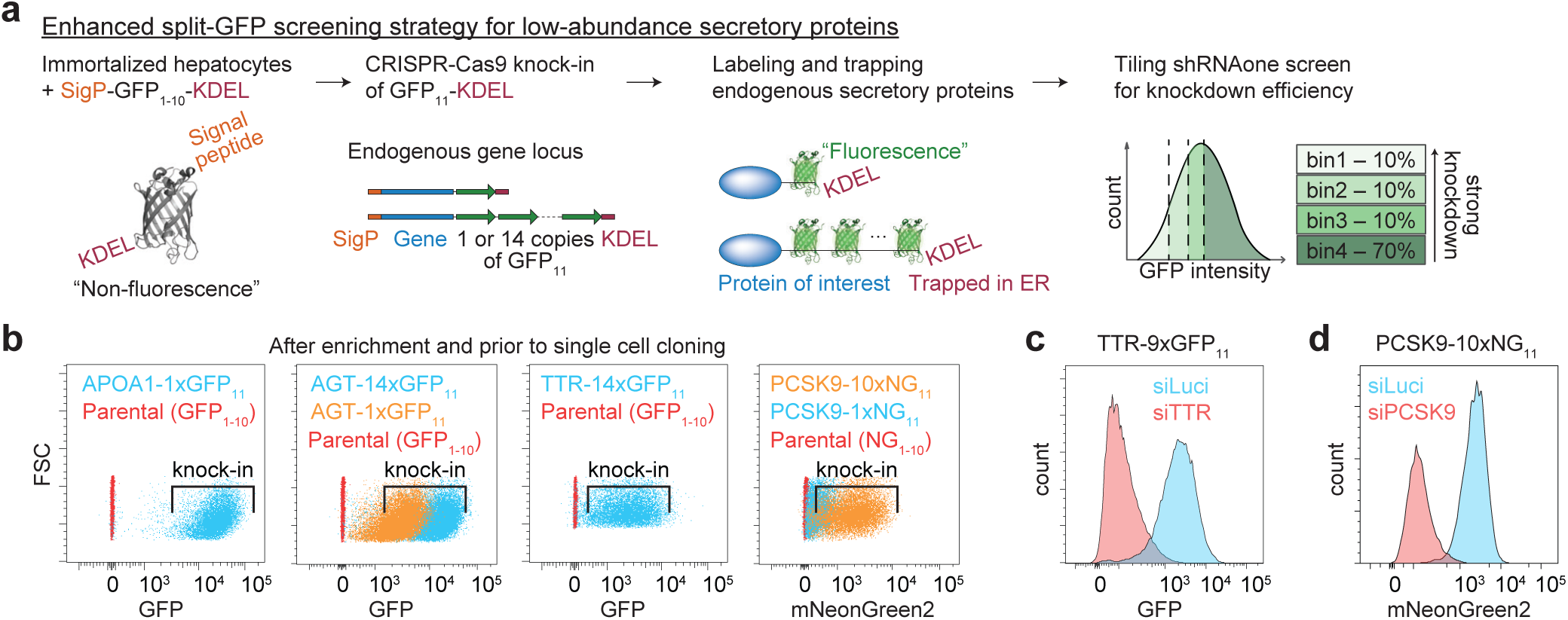
Enhanced split-GFP screening strategy for low-abundance secretory proteins. **a** MIHA cell line expressing ER-retained GFP1–10 (Signal Peptide [SigP]–GFP1–10–KDEL) is non-fluorescent. CRISPR–Cas9 knock-in appends single or tandem GFP11 tag(s) and a KDEL signal to the C terminus of an endogenous secretory gene; complementation in the ER lumen gives fluorescence proportional to protein level and KDEL ensures ER localization. After library transduction, cells are sorted by fluorescent signal into four bins (bin1, lowest 10%; bin2, bin3, next 10% each; bin4, top 70%), giving the knockdown score log2(bin1/bin4). **b** Flow cytometry of polyclonal knock-in populations before single-cell cloning: GFP or mNeonGreen2 (mNG2) fluorescence of the indicated knock-ins, overlaid on parental GFP1–10 or mNG21–10 cells (red). Brackets, knock-in cells. **c, d** Validation of TTR–9×GFP11 (c) and PCSK9–10×mNG211 (d) reporter cell lines using siRNA.

### Single-nucleotide tiling with shRNAone identifies efficient siRNA target sites and reveals “clustered potent sites” on the TTR mRNA

*TTR* is the gene responsible for transthyretin amyloidosis and the clinically validated target for patisiran, the first approved siRNA therapeutic^1^. To test our platform against this established standard, we performed a pilot shRNAone tiling screen for *TTR* knockdown efficiency at 5-nt resolution. The shRNAone library comprised two sublibraries that differed only in the 5′-terminal nucleotide of the guide, utilizing either a starting uridine or adenosine (Supplementary Table 8a–c). The pilot screen identified the patisiran target sequence among the most potent sites, indicating that the shRNAone tiling method can reliably recover a previously validated siRNA target site (Supplementary Fig. 17a–c). Furthermore, the newly identified potent sites demonstrated high reproducibility between U- and A-starting sublibraries and across two biological replicates (Supplementary Fig. 17d, e).

We then tiled the transcript at single-nucleotide resolution against the NM_000371.3 reference, yielding 909 targeting shRNAs and 100 non-targeting controls (Supplementary Table 8d–f). shRNAs whose target sites fell entirely within 1–110 nt or 727–929 nt of that reference were not enriched above non-targeting controls (*P* = 0.804, Mann-Whitney U test). Because these intervals correspond exactly to the 110 nt of 5′ UTR and 212 nt of 3′ UTR removed when the annotation was revised to NM_000371.4 (total length 616 nt; CDS spanning 27–470 nt), this lack of enrichment indicates that these regions are absent from the mature transcript in MIHA cells (Fig. 4a, b, Supplementary Fig. 18a).

**Fig. 4:**
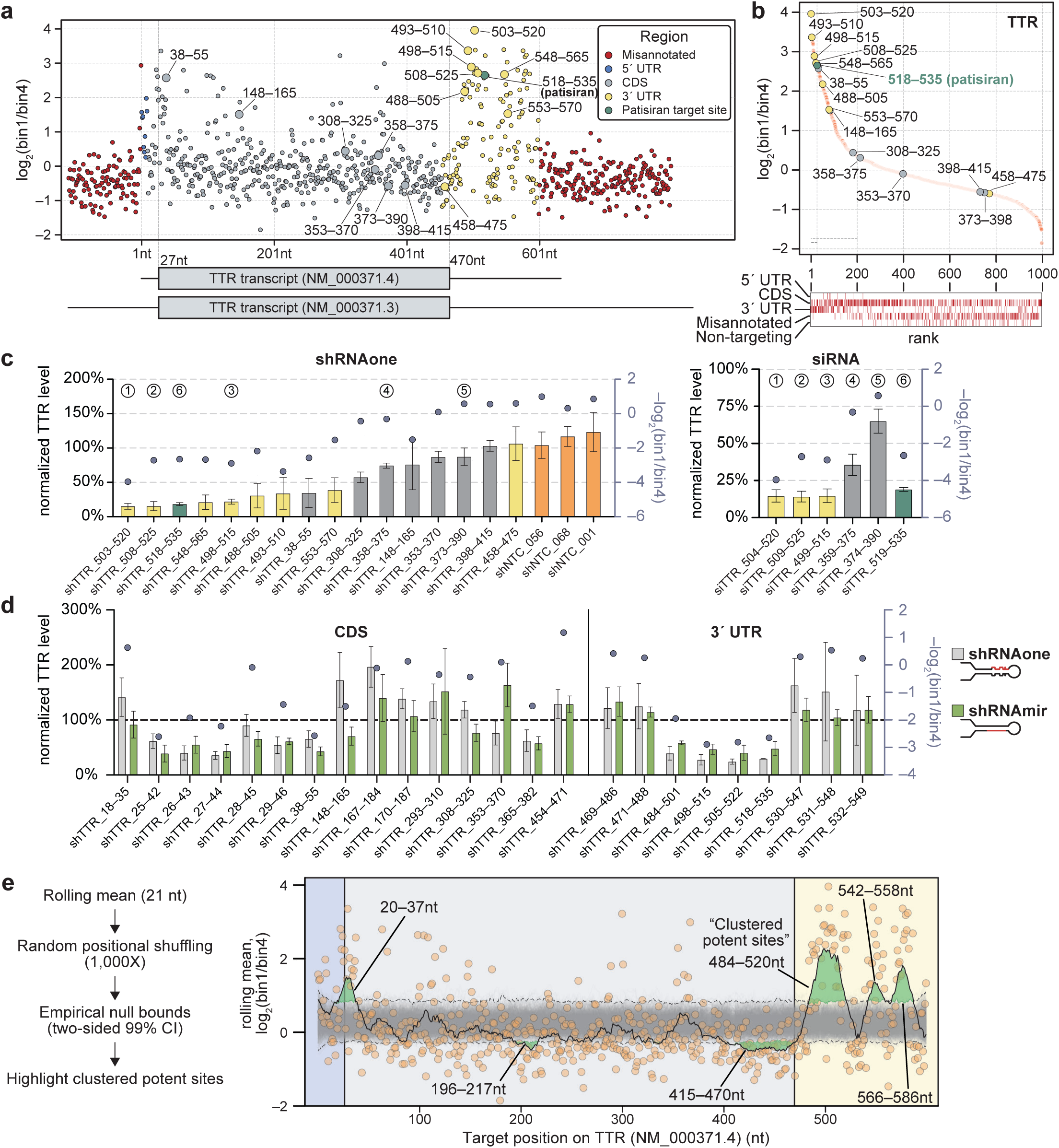
Single-nucleotide tiling with shRNAone identifies efficient siRNA target sites and reveals “clustered potent sites” on the TTR mRNA. **a** shRNAone 1-nt tiling screen on TTR. log2(bin1/bin4) for every library shRNA is plotted versus target-site position and colored by region. Misannotated, shRNAs tiling a sequence unique to the superseded annotation NM_000371.3 and absent from NM_000371.4. Enlarged points, shRNAs validated in (c). Transcript models below (boxes, CDS 27–470 nt; lines, UTRs). **b** Library shRNAs ranked by log2(bin1/bin4), labeled as in (a). **c** Validation of individual shRNAone (left) and siRNA (right) constructs. Bars (left axis), TTR level measured by qRT-PCR, normalized to non-shRNA control (left) or siLuc (right); circles (right axis, inverted), −log2(bin1/bin4) from (a). Colors as in (a); circled numbers, matched shRNA/siRNA sites. **d** shRNAone (grey) versus shRNAmir (green) at matched CDS and 3′ UTR sites, plotted as in (c); schematics at right, guide strand red. Dashed line, 100%. **e** Clustered potent sites defined by random positional shuffling analysis. Left, workflow. Right, shRNAone log2(bin1/bin4) (orange), 21-nt rolling mean (black), null distribution from 1,000 rounds of random positional shuffling (grey) and its 99% confidence interval (dashed). Green, clustered potent sites (rolling mean exceeding the upper null bound). Background, 5′ UTR (blue), CDS (grey), 3′ UTR (yellow). Data in c, d are mean values ± s.d. of n = 3 biologically independent experiments.

To exclude any potential artifacts arising from the split-GFP reporter, such as ER stress, 16 targeting and 3 non-targeting shRNAs were subsequently validated individually in wild-type MIHA cells. Knockdown of endogenous *TTR* measured by qRT-PCR strongly correlated with screen enrichment (Fig. 4c, left, Supplementary Fig. 18b, Supplementary Table 8g; *r* = 0.937, *P* = 3.65 × 10^-9^). This correlation confirms that phenotypic ranking in the knock-in reporter system accurately reflects the silencing of the native mRNA. Finally, synthetic siRNAs targeting six of these validated sites reproduced the same functional relationship (Fig. 4c, right; matching circled numbers denote identical target sites used in the shRNAone experiments). This outcome establishes that single-nucleotide tiling with shRNAone can systematically map efficient siRNA target sites across an entire transcript.

Across the tiled transcript, shRNAs targeting the CDS yielded consistently lower enrichment than those targeting the 3′ UTR (Fig. 4b). Because shRNAone retains activity in the absence of AGO2, likely due to the exploitation of non-slicing AGO proteins, we tested whether this positional bias reflected an intrinsic property of the shRNAone backbone or a general feature of RNAi. We cloned identical guide sequences targeting 15 CDS sites and nine 3′ UTR sites into both shRNAone and the conventional shRNAmir backbone, whose activity is largely dependent on AGO2 (Supplementary Fig. 7c, Supplementary Table 8g). Only two of the 15 coding sequence constructs and none of the nine 3′ UTR constructs exhibited significant differences in knockdown efficiency between the two backbones (Fig. 4d, Supplementary Fig. 18c). The low efficiency of CDS sites is therefore backbone-independent, suggesting that the 3′ UTR is an ideal target region for designing potent siRNA therapeutics.

Interestingly, we observed that many potent sites were clustered along the *TTR* mRNA. To define cluster boundaries, we compared the rolling mean of shRNA enrichment along the transcript with an empirical null distribution obtained from 1,000 rounds of random positional shuffling of the same scores across all positions. Clustered potent sites were defined as contiguous regions in which the observed rolling mean exceeded the 99.5th percentile of this null distribution (Fig. 4e, left). This empirical threshold resolved four distinct clusters of potent sites on the *TTR* transcript (Fig. 4e, right, Supplementary Fig. 19a). One cluster flanked the start codon, while the remaining three localized to the 3′ UTR. Khvorova and colleagues recently reported similar clusters of potent siRNA target sites across several genes, including *APP*, and our findings in *TTR* corroborate and extend this observation^55^.

### Broad utility of shRNAone tiling for identifying potent therapeutic siRNA target sites

*PCSK9* is a key driver of hypercholesterolemia and serves as the clinically validated target for inclisiran^2^. Having established the shRNAone tiling method, we therefore applied this approach to the *PCSK9* transcript at 3-nt resolution (Fig. 5a, Supplementary Fig. 20a, b, Supplementary Table 9a–c). Random positional shuffling analysis resolved 7 clusters of potent sites on the *PCSK9* transcript (Fig. 5b, Supplementary Fig. 19b). All 7 potent clusters mapped to the 3′ UTR, whereas all 11 regions with rolling means below the lower empirical bound (0.5^th^ percentile of the shuffled distribution) localized exclusively to the 5′ UTR or CDS (Fig. 5b). The shRNA matching the inclisiran target sequence ranked 31st out of 1,230 constructs, placing it in the top 3% and demonstrating that the screen successfully recovers a known, highly potent target site (Fig. 5a). For individual validation, we selected 19 targeting shRNAs, including the inclisiran match, alongside three non-targeting controls. We performed these validations in unmodified, wild-type MIHA cells, where qRT-PCR measurements of endogenous *PCSK9* knockdown strongly correlated with the pooled screen scores (Fig. 5c, Supplementary Fig. 20c, Supplementary Table 9d; *r* = 0.819, *P* = 3.18 × 10^−6^).

**Fig. 5:**
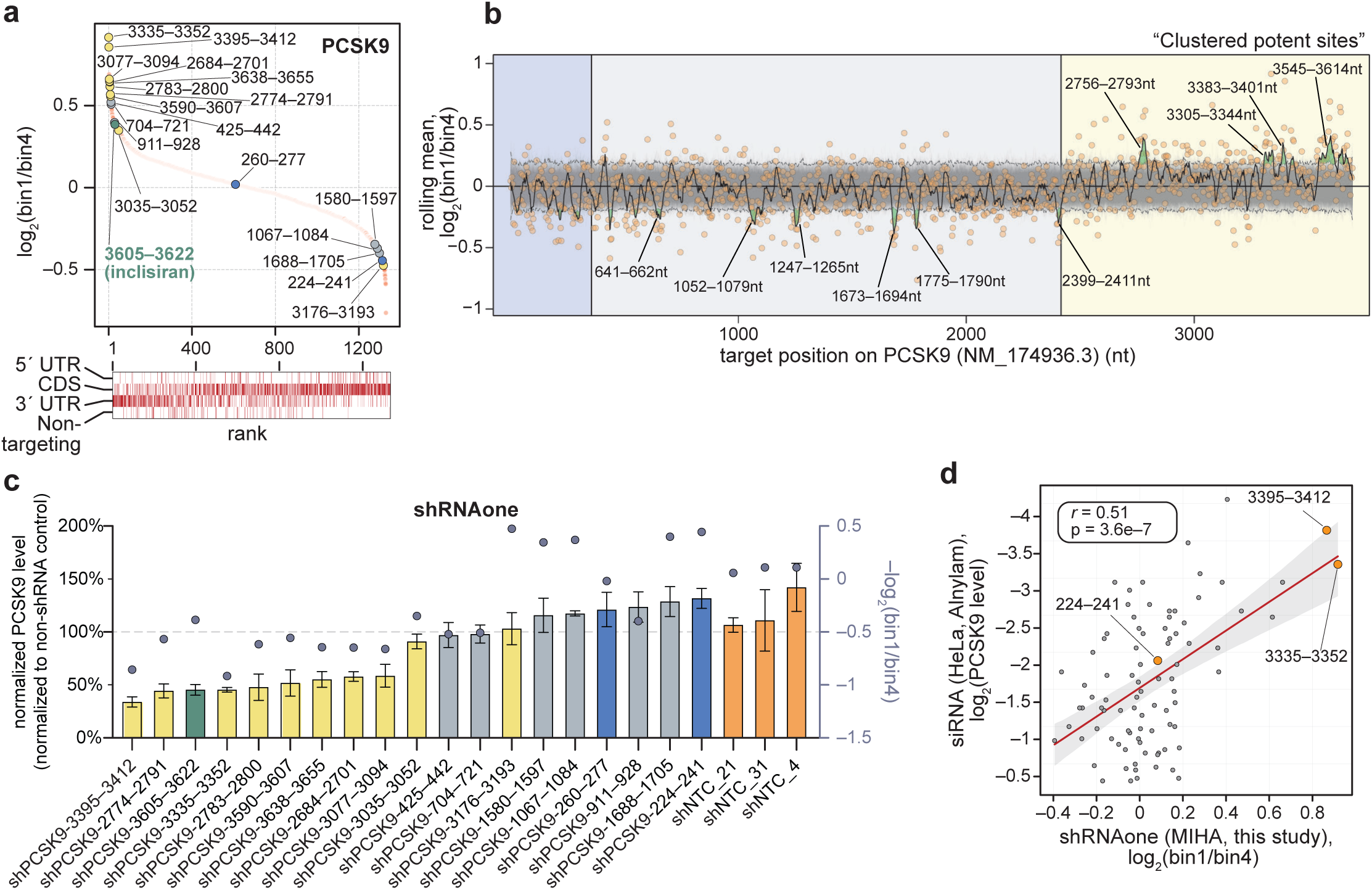
Tiling shRNA screening identifies potent RNAi target sites on the PCSK9 mRNA. **a** Library shRNAs ranked by log2(bin1/bin4), from a tiling shRNAone screen in PCSK9–10×mNG211 MIHA reporter cells. **b** Clustered potent sites defined by random positional shuffling analysis. 21-nt rolling mean of log2(bin1/bin4) (black) with individual shRNAone log2(bin1/bin4) (orange), the null distribution from 1,000 rounds of random positional shuffling (grey) and its 99% confidence interval (dashed). Green, clustered potent sites (rolling mean exceeding the upper null bound). Background, 5′ UTR (blue), CDS (grey), 3′ UTR (yellow). **c** Validation of individual shRNAone constructs in WT MIHA cells. Bars (left axis), PCSK9 mRNA measured by qRT-PCR and normalized to the non-shRNA control (dashed line, 100%); colors as in Supplementary Fig. 20b, with non-targeting controls (shNTC_21, shNTC_31, shNTC_4) in orange. Circles (right axis, inverted), the corresponding −log2(bin1/bin4) from (a). **d** Comparison with an independent siRNA dataset in HeLa cells (Alnylam, U.S. Patent No. 8,809,292 B2, 2014) across 85 shared target positions. x, shRNAone log2(bin1/bin4) in MIHA cells (this study); y, log2 PCSK9 level after siRNA treatment. Red line, linear fit; grey band, 95% confidence interval; orange, labeled sites. Two-sided Pearson correlation. Data in (c) are mean values ± s.d. of n = 3 biologically independent experiments.

Encouraged by this strong correlation, we next investigated whether our shRNAone screen could predict the site-level knockdown activity measured on an entirely independent platform. We compared our shRNAone knockdown data generated in MIHA cells with an Alnylam siRNA dataset generated in HeLa cells (U.S. Patent No. 8,809,292 B2, 2014). Target knockdown correlated significantly across the two datasets (Fig. 5d; n = 85, *r* = 0.51, *P* = 6.2 × 10^−7^), suggesting that site-level targeting rules transfer robustly between the two formats. This cross-platform predictive power further supports the use of shRNAone tiling as a highly scalable pre-screening strategy for prioritizing candidate target sites in siRNA drug development.

### shRNA, CasRx, and PspCas13b share clustered potent sites in the 3′ UTR of TTR mRNA

Building on our initial 1-nt shRNAone tiling screen across the *TTR* transcript, we extended this high-resolution strategy to two representative CRISPR systems, CasRx and PspCas13b. This approach enables a direct comparison of RNAi- and CRISPR-mediated silencing landscapes across the same mRNA. To optimize target repression, we expressed cytosolic crRNAs using a U1 promoter system known to enhance Cas13-mediated RNA degradation (Fig. 6a, Supplementary Table 10, 11)^56^. When evaluating the overall knockdown profiles as Gaussian-smoothed trends, the CasRx data largely reproduced the broad architecture of the shRNAone landscape (Fig. 6a, b, Supplementary Fig. 21). Specifically, CasRx guide enrichment declined across the CDS and resolved into discrete activity peaks within the 3′ UTR (Fig. 6b). In contrast, PspCas13b yielded many potent sites in both the CDS and the 3′ UTR, indicating that its target-site selection is not restricted to the 3′ UTR (Fig. 6b, Supplementary Fig. 22).

**Fig. 6:**
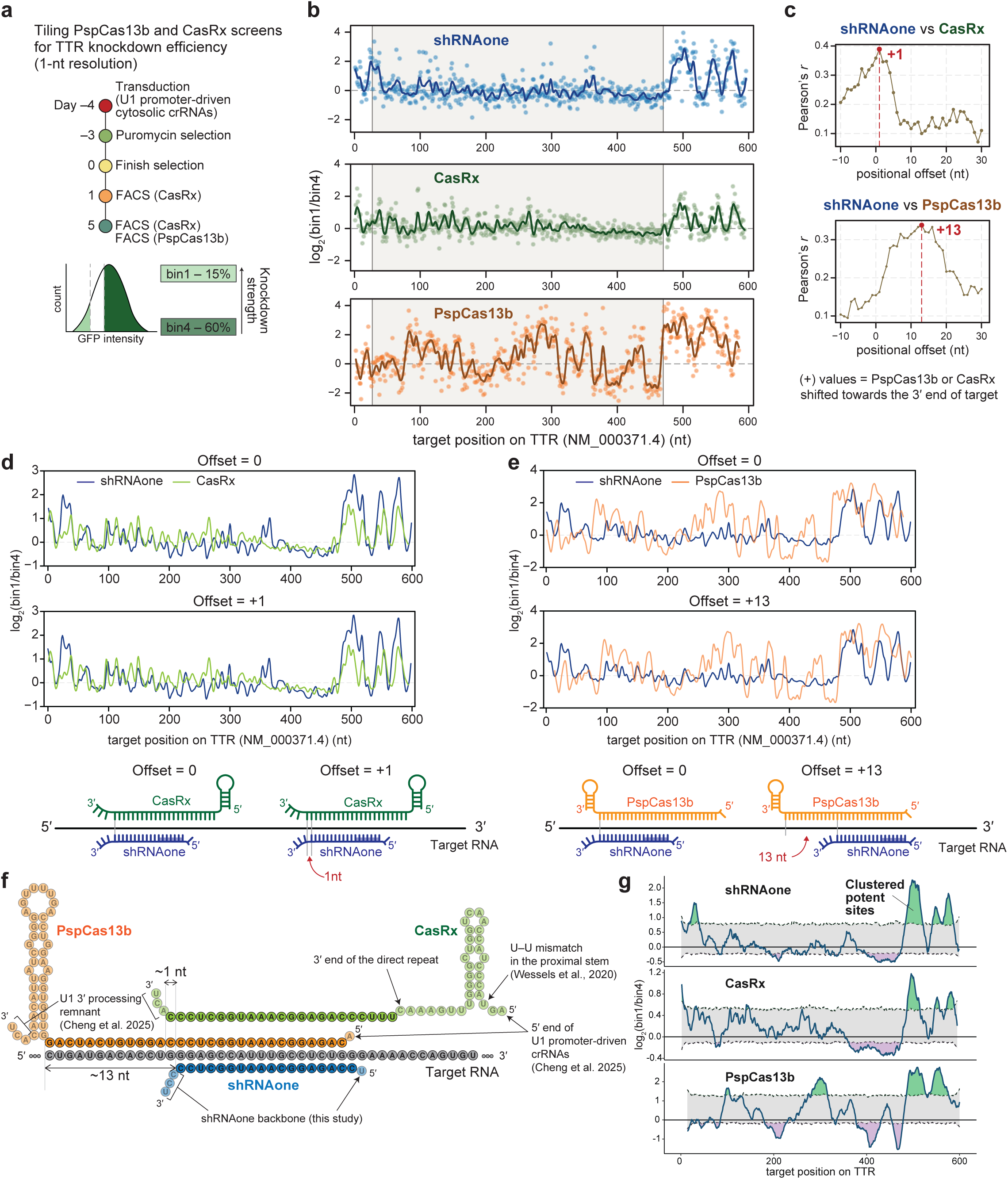
shRNA, CasRx, and PspCas13b share clustered potent sites in the 3′ UTR of TTR mRNA. **a** Design of the 1-nt-resolution tiling PspCas13b and CasRx knockdown screens in reporter cells. Knockdown strength is scored as log2(bin1/bin4). **b** Knockdown profiles across the TTR transcript (NM_000371.4) for shRNAone (blue, mean of biological triplicates), CasRx (green, day-5 data), and PspCas13b (orange, mean of biological triplicates). Points, individual shRNA or crRNA; lines, Gaussian smoothed line. Grey shading, coding sequence (27–470 nt). shRNAone data are re-plotted from Fig. 4. **c** Positional offset Pearson correlation between the shRNAone profile and the CasRx (top) or PspCas13b (bottom) profile. **d, e** Profiles overlaid at zero offset (top) and at the offset when correlation reaches maximum (bottom) for shRNAone versus CasRx (d) and shRNAone versus PspCas13b (e). Schematics below show the corresponding alignment of the crRNA spacer and the shRNAone guide on the target RNA. **f** Nucleotide-level alignment of the PspCas13b crRNA, CasRx crRNA, and shRNAone guide on a common target RNA. **g** Clustered potent sites of shRNAone (top), CasRx (middle), and PspCas13b (bottom) on the TTR mRNA (rolling mean window = 21 nt). The overlap between systems is shown for visual comparison and was not tested statistically.

To compare these knockdown profiles quantitatively, we performed a pairwise cross-correlation analysis using the raw data points rather than the smoothed trend lines. By systematically shifting the positional offset of the Cas13 profile relative to the shRNAone profile along the transcript, we identified the specific alignments that maximized their functional concordance. The highest correlation between shRNAone and CasRx occurred at an offset of +1 nt (maximum Pearson’s *r* = 0.389; Fig. 6b–d). In contrast, the optimal alignment between shRNAone and PspCas13b clustered at offsets of +11 to +14 nt, peaking at +13 nt (maximum *r* = 0.338; Fig. 6b, c, e). Mapping the guides onto a shared target sequence at this +13 nt offset superimposes the shRNA seed region (positions 2 to 8) directly onto the 5′ region of the PspCas13b crRNA, a region previously established as a critical seed-like recognition segment^57^. Similarly, the +1 nt offset between CasRx and shRNAone yields a modest overlap of the two guide sequences, consistent with their comparable lengths (Fig. 6f, Supplementary Fig. 23).

Next, to determine whether the clustered potent sites identified by shRNAone, CasRx, and PspCas13b overlap, we applied random positional shuffling to their respective *TTR* knockdown profiles (Fig. 6g). Consistent with our Gaussian-smoothed trends, these high-activity clusters overlapped substantially within the *TTR* 3′ UTR. This suggests that these silencing hotspots reflect intrinsic properties of the target mRNA, so these regions are promising targets for multiple platforms.

### Target RNA accessibility is associated with guide efficacy across diverse targeting systems

Given the correlated knockdown profiles observed among shRNAone, CasRx, and PspCas13b, a substantial component of targeting efficiency appears to be intrinsically encoded within the target mRNA. To investigate this, we evaluated target-site accessibility (the probability of being unpaired) using RNAplfold (Fig. 7a)^58,59^. The RNAplfold algorithm relies on three parameters to calculate target-site accessibility (Fig. 7a). *W* specifies the size of the sliding window, *L* defines the maximum distance allowed between pairing bases, and *U* dictates the length of the RNA stretch evaluated for single-strandedness (Fig. 7a, Supplementary Fig. 24a)^58^.

**Fig. 7:**
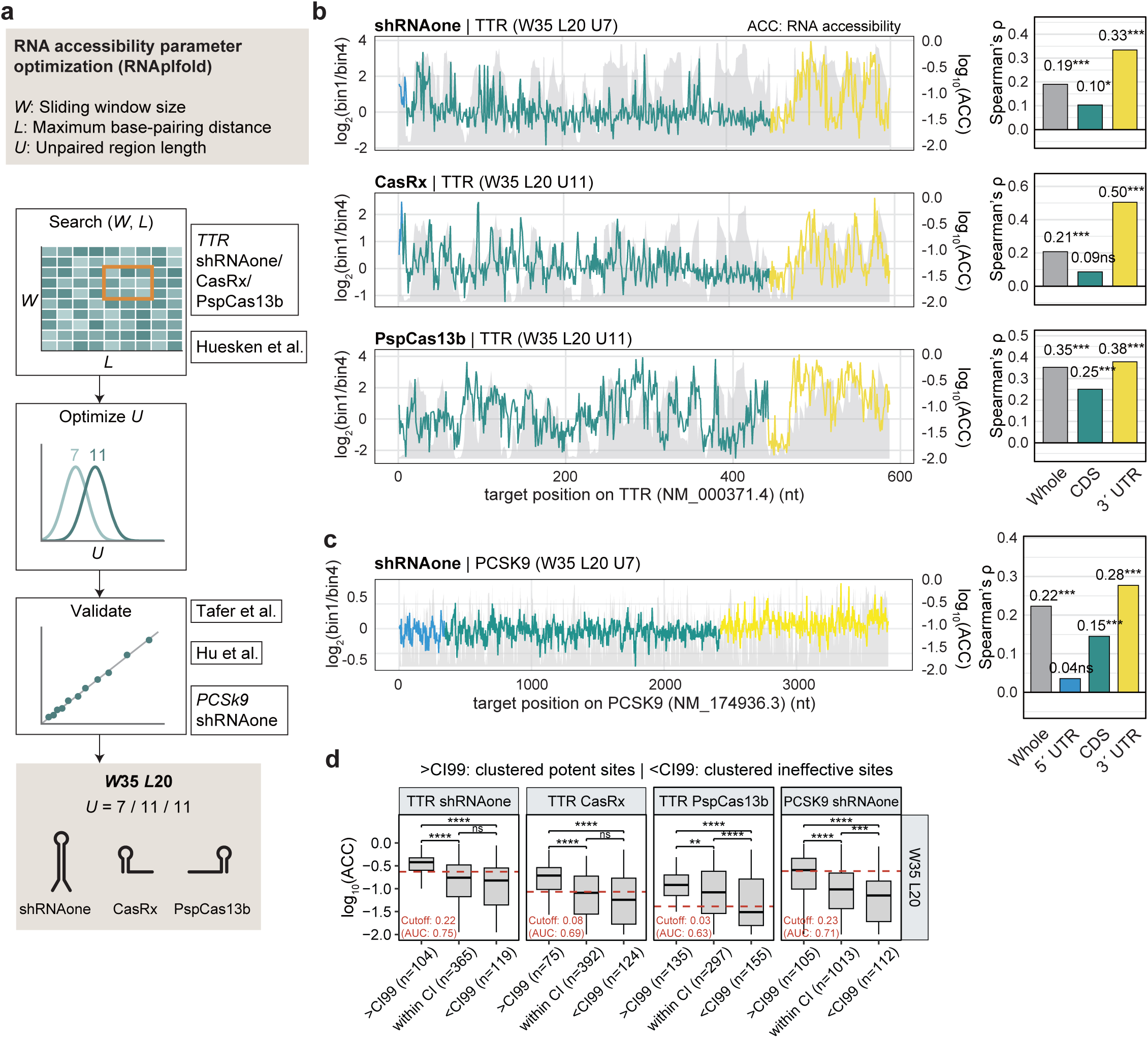
Target RNA accessibility is associated with guide efficacy across diverse targeting systems. **a** Pipeline for accessibility parameter optimization. See also Supplementary Fig. 24a. **b** Dual-track visualization of predicted target accessibility and experimentally measured knockdown activity for three RNA-targeting effectors (shRNAone, CasRx, and PspCas13b) across the TTR transcript (NM_000371.4). For each effector, the grey shaded area depicts the log10-transformed 3′-end unpaired probability computed by RNAplfold under the optimal parameter set (W = 35, L = 20; shRNAone U = 7, CasRx U = 11, PspCas13b U = 11), while the colored line traces the mean log2FC of enrichment, with colors indicating transcript region (5′ UTR, grey; CDS, cyan; 3′ UTR, yellow). Accompanying bar charts (right) report Spearman correlation coefficients between accessibility and log2FC within the Whole, CDS, and 3′ UTR regions. **c** Application of the optimal TTR-derived parameter set (W35_L20_U7) to the PCSK9 transcript. Right panel: Spearman correlation between predicted accessibility and experimentally measured shRNAone log2FC across the Whole, 5′ UTR, CDS, and 3′ UTR regions of PCSK9 (NM_174936.3). **d** Accessibility comparison across random positional shuffling-defined target-site clusters on TTR targeted by shRNAone, CasRx, and PspCas13b (panels 1–3) and PCSK9 targeted by shRNAone (panel 4). Clusters were defined by comparing the rolling mean (window = 21 nt) of log2FC against the distribution obtained from 1,000 rounds of random positional shuffling; sites exceeding the 99.5th percentile of the null distribution were classified as “clustered potent sites” (>CI99), those falling below the 0.5th percentile as “clustered ineffective sites” (<CI99), and the remainder as background (within CI). Box plots show log10-transformed accessibility under W35_L20. Red dashed lines mark the optimal classification threshold determined by Youden’s index, with the corresponding AUC reported. Pairwise Wilcoxon test significance is indicated by asterisks (*P < 0.05, **P < 0.01, ***P < 0.001, ****P < 0.0001).

Previously, Hofacker and colleagues applied this computational method to large-scale siRNA datasets and determined the optimal parameters for siRNAs (*W* = 80, *L* = 40, and *U* = 8 or 16)^60,61^. In our study, however, because different RNA-targeting systems likely have distinct binding requirements, we needed to systematically re-determine the optimal parameters. We first conducted a combinatorial screen of RNAplfold parameters and identified the most informative binding mode at the target site (3′; Supplementary Fig. 24a, b), the dominant non-linear parameter (*U*; Supplementary Fig. 24c), and the optimal range of *U* for predicting *TTR* knockdown (PspCas13b: 9–23 nt, CasRx: 8–14 nt, shRNAone: 6–14 nt, at *W*80 and *L*40; Supplementary Fig. 24d). Notably, the range of optimal *U* values for PspCas13b was generally longer than for shRNAone, suggesting PspCas13b requires a more extended accessible region to initiate binding.

We next searched for the optimal *W* and *L* parameters within these *U* ranges (Fig. 7a, Supplementary Fig. 25a). When evaluated against the large-scale siRNA dataset (n = 2,327)^61^, a new parameter set (*W*35, *L*20) demonstrated predictive performance equal to or slightly exceeding that of the original parameters (*W*80, *L*40)^60^ (Supplementary Fig. 25b). Using this refined model (*W*35, *L*20), we systematically re-evaluated *U* to identify the sequence length yielding the highest Spearman correlation. We observed peak performance at *U* = 7 for shRNAone and *U* = 11 for both PspCas13b and CasRx, and therefore adopted these values for all subsequent analyses (Fig. 7a, Supplementary Fig. 25c).

To assess generalizability, we next evaluated the optimized parameters on two independent datasets excluded from all optimization steps (Supplementary Fig 25d). For shRNAone (W35_L20_U7), we used the siRNA evaluation set (n = 360) from Tafer et al.^60^, which was excluded from the original W80_L40_U8 parameter derivation. For PspCas13b (W35_L20_U11), we used tiling data (n = 117) from Hu et al.^62^. Efficient and precise U6-driven expression requires a purine at the transcription start site^63^. Because the Hu et al. library did not incorporate this requirement, we analyzed the full crRNA set (n=117) and the purine-starting subset (n=54) separately. Predicted accessibility correlated modestly but significantly with knockdown across all three evaluations (shRNAone: Spearman’s *r* = 0.20, *P* = 0.0002; PspCas13b full: *r* = 0.19, *P* = 0.045; PspCas13b purine-only: *r* = 0.28, *P* = 0.047; Supplementary Fig. 25d), indicating that the optimized parameters generalize beyond the data used to select them.

Circular shift analysis showed that the observed correlations under the optimized parameters (W35_L20) significantly exceeded a null distribution designed to preserve spatial autocorrelation for PspCas13b and CasRx, whereas the shRNAone correlation was marginal (empirical *P* = 0.059; Supplementary Fig. 25e). This targeted optimization strategy allowed us to define the maximum achievable correlation when using RNA accessibility as the sole predictor of guide efficacy.

Next, using these optimal parameter sets, we investigated regional differences in predictive power by comparing the Spearman correlations derived from the CDS against those from the 3′ UTR (Fig. 7b, Supplementary Fig. 26a). PspCas13b maintained a robust correlation across both the CDS and the 3′ UTR, suggesting that its knockdown activity is associated with local RNA accessibility regardless of regional context (Fig. 7b, Supplementary Fig. 26a). In contrast, shRNAone and CasRx exhibited strong correlations exclusively within the 3′ UTR, indicating that local accessibility predicts efficacy in the 3′ UTR but provides limited predictive power within the CDS (Fig. 7b, Supplementary Fig. 26a).

We next tested cross-transcript generalizability on the *PCSK9* shRNAone tiling data, which were not used at any stage of parameter optimization. Consistent with the *TTR* data, the *PCSK9* knockdown profile exhibited a significantly higher correlation with predicted accessibility in the 3′ UTR than in the CDS (Fig. 7c). Together with the external evaluations above, this indicates that the optimized parameters generalize across transcripts and effectors.

Finally, we confirmed that the clustered potent sites previously identified in *TTR* and *PCSK9* via shRNAone profiling exhibit significantly higher overall RNA accessibility compared to either background regions or clustered ineffective sites (Fig. 7d). Strikingly, the potent clusters defined independently by CasRx and PspCas13b profiling on *TTR* demonstrated the exact same accessibility enrichment (Fig. 7d). Together, these findings suggest that extended local RNA accessibility is a shared molecular basis for the clustered potent sites across all three systems.

### A transcriptome-wide resource for PspCas13b putative guide design

Recently, several machine learning or deep learning prediction tools for CasRx utilizing comprehensive feature engineering have been published, while the only existing PspCas13b guide design tool (Fareh2024) is provided by Hu et al. and relies on a knowledge-based heuristic scoring algorithm.^62,64–67^. Inspired by this landscape, we compared our thermodynamic accessibility models against these complex or heuristic algorithms. Building on the robust baseline established above, we evaluated our optimized RNAplfold parameters (*W*35_*L*20) head-to-head against these external tools (Supplementary Fig. 27a, b). For CasRx, dedicated tools such as Sanjana2020^64^, TIGER2023^65^, and Hsu2023^66^ generally outperformed simple RNA accessibility (Supplementary Fig. 27a, b). Notably, these CasRx-specific algorithms could effectively predict potent target sites even within the *TTR* CDS, highlighting the capacity of advanced feature engineering to capture complex sequence determinants beyond local RNA folding. By contrast, another CasRx tool, DeepCas13^67^, exhibited weaker predictive power on the *TTR* transcript (Supplementary Fig. 27a, b), and its maximally correlated guide position failed to align with our empirical data (Supplementary Fig. 27a, +2 position).

Because Fareh2024 is currently the sole guide prediction tool for PspCas13b^62^, we evaluated it alongside the CasRx algorithms against our PspCas13b screening data (Supplementary Fig. 27a, b). All tested external models exhibited poor predictive power for PspCas13b-mediated knockdown of *TTR*, even across various offset positions (Supplementary Fig. 27a, b). This lack of generalization demonstrates that currently available tools cannot reliably predict potent PspCas13b guide sequences. Conversely, our simple RNA accessibility models maintained a good predictive advantage for PspCas13b (Supplementary Fig. 27b). On this basis, we applied our optimized parameter sets (*W*35_*L*20_*U*11) across all human protein-coding transcripts, computing RNAplfold accessibility for every 30-nt PspCas13b target window and ranking guides within each transcript by descending accessibility. We provide these rationally designed crRNA sequences for PspCas13b via a publicly available website (https://cas13b.chullab.com; an example gene is shown in Supplementary Fig. 27c).

## DISCUSSION

In this study, we developed shRNAone, an optimized scaffold engineered for highly homogeneous small RNA expression, and utilized it to generate high-resolution knockdown profiles of RNAi and Cas13 systems in human cells. To demonstrate the precision of this scaffold, we systematically dissected the functional dependencies of seed and supplementary pairings, which had not been previously evaluated at a large scale *in cellulo*. We next demonstrated the utility of shRNAone tiling in identifying highly effective siRNA target sites, offering a highly scalable pre-screening strategy for the development of siRNA therapeutics. Most importantly, our comparative single-nucleotide tiling screens revealed the existence of “clustered potent sites” shared by mechanistically distinct RNA-targeting systems. Because these clusters coincide with locally accessible regions shared across platforms, they offer promising target sites for future RNA therapeutics. Collectively, these high-resolution profiles provide a comprehensive guide for designing optimal RNA-targeting tools.

### shRNAone as a precision platform for small RNA biology

Evolutionary analysis of miRNAs reveals that strict sequence conservation frequently extends beyond the seed region^39^. This constraint is traditionally attributed to the necessity of maintaining specific duplex structures for precise DROSHA and DICER processing. However, because single mismatches do not alter overall duplex length, the biogenesis machinery can likely tolerate these variations, suggesting that additional functional pressures drive this conservation. One such pressure is target-directed miRNA degradation (TDMD), a regulatory mechanism requiring extensive complementarity between the miRNA and its target^68–74^. Alternatively, non-seed conservation likely reflects a strict functional requirement for supplementary pairing to repress specific target RNAs in certain cellular or developmental contexts. As demonstrated by our *RPLP0* validation, a canonical 8-mer seed match is sometimes insufficient for target repression, rendering 3′-supplementary pairing indispensable. Furthermore, relative to the baseline correlation between the miR_1A and miR_1U libraries (Pearson’s *r* = 0.858; Fig. 2d), the reduced correlation of the miR_7merSupp library with miR_1U (*r* = 0.631; Fig. 2e) indicates that the central nucleotides (positions 9 to 12) and the tail region (positions 17 to 22) modulate specific miRNA–target interactions. Identifying those non-seed pairings that drive cellular phenotypes remains a critical challenge. The SynthoMir library therefore provides a functional platform to systematically map these complex targeting dependencies.

### Clustered potent sites in 3′ UTRs: implications for therapeutic design

Our high-resolution shRNAone tiling screens revealed that potent target sites are clustered extensively within the 3′ UTR of the *TTR* transcript. Subsequent analysis demonstrated that two distinct Cas13 systems exhibit strong knockdown efficacy at these same loci, and this shared positional preference is associated with local RNA accessibility. Consistent with these observations, Khvorova and colleagues recently reported a similar clustering of highly active siRNA target sites across several transcripts, including *APP*^55^. Both studies highlight strong enrichment of these targetable regions in the 3′ UTR. This spatial preference aligns closely with early mechanistic studies by Bartel and colleagues, which established that effective miRNA target sites preferentially reside near both ends of the 3′ UTR^39^. Because directing siRNAs or other RNA-targeting therapeutics, including antisense oligos or circular RNAs, toward these clustered regions has the potential to be more robust than relying on isolated target sites. Thus, systematically mapping these highly permissive regions across the human transcriptome represents a critical objective for future therapeutic development. Importantly, our study demonstrates that any of the three effectors we tested can be utilized to experimentally identify “clustered potent sites.”

### PspCas13b as an efficient platform for targeting the coding sequence

This study provides the first single-nucleotide resolution comparison of three representative RNA knockdown systems, specifically shRNAone, CasRx, and PspCas13b. Overall, shRNAone and CasRx exhibited highly concordant knockdown profiles, whereas PspCas13b displayed a divergent targeting pattern, particularly within the CDS. Because the RNA accessibility profile of the *TTR* mRNA closely aligned with the PspCas13b knockdown profile (Fig. 7b), it appears that shRNAone and CasRx are inefficient at silencing CDS target sites even when those regions are open. In contrast, PspCas13b efficiently knocks down target mRNAs through accessible regions across both the CDS and the 3′ UTR. These distinct positional biases suggest that PspCas13b effectively competes with ribosomes, whereas shRNAone and CasRx are readily displaced by translation^75^. Supporting this mechanistic hypothesis, we and others have recently demonstrated that catalytically dead PspCas13b (dPspCas13b) can effectively block translation by tightly binding the start codon of a target mRNA^56,76^. These insights provide a rational framework for selecting the optimal Cas13 system for specialized applications, such as RNA base editing or splicing modulation.

### A guide RNA resource for PspCas13b

PspCas13b has not been widely used in mammalian applications compared to CasRx, particularly in lentiviral pooled screens. This is largely because its direct repeat contains an internal termination signal (i.e., a poly-U tract) that truncates crRNAs expressed from RNA Pol III promoters, such as U6 (Fig. 6f). We recently resolved this issue by driving crRNA expression with a Pol II-dependent U1 promoter, which enables robust PspCas13b-mediated RNA knockdown in human cells^56^. A second limitation lies in guide design. Whereas several predictive models exist for CasRx, only one tool is currently available for PspCas13b^62^. However, because their training library did not fix the first nucleotide to a purine which is required for efficient U6 transcription^63^, the resulting algorithm heavily weights a 5′ guanosine in the guide RNA (pairing with a 3′ C on the target site). We propose that this parameter captures a transcriptional requirement rather than an intrinsic targeting rule of the nuclease. Consequently, when an expression vector supplies a constant transcription start nucleotide, such as an A for the U1 promoter, this sequence preference no longer applies. Indeed, the Fareh2024 model exhibited limited correlation with our PspCas13b dataset, whereas established CasRx algorithms strongly correlated with our CasRx screens. To address the need for a generalizable guide design tool, and given that PspCas13b efficacy strongly correlates with local RNA accessibility, we used RNA accessibility profiles to rationally design PspCas13b crRNAs for all human mRNAs. By sharing these pre-computed targets online, we aim to help researchers utilize PspCas13b for applications that require robust RNA binding or cleavage in human cells.

### Limitations

While our high-resolution knockdown profiling reveals clustered potent sites, two primary limitations need to be addressed to generalize these findings. First, experimentally, our single-nucleotide tiling is currently restricted to a limited number of transcripts. Although we systematically mapped *TTR* (using shRNAone, CasRx, and PspCas13b) and *PCSK9* (using shRNAone), a much larger and more diverse cohort of genes must be profiled to definitively establish these clusters as a universal feature of human mRNAs. Second, computationally, local RNA accessibility only partially explains the formation of these clusters. Even after rigorous parameter optimization, accessibility yields a modest correlation with knockdown efficacy (Spearman’s *r* ≈ 0.3). While this thermodynamic metric successfully identifies broadly permissive regions, it leaves substantial site-to-site variation unexplained. Consequently, other unidentified biological determinants must also govern target-site efficacy, highlighting the need for more comprehensive predictive models to fully decode the rules of clustered potent sites.

## MATERIALS AND METHODS

### Plasmid construction for shRNA

shGAPDH GIPZ plasmid (pGIPZ) was purchased from Horizon Discovery (cat# RHS4371). The GFP coding sequence was replaced by the mCherry coding sequence with NotI and NcoI restriction sites, and the new plasmid was termed pRIPZ. Stuffer was placed between XhoI and EcoRI restriction sites for efficient cloning. shRNAs were subsequently cloned into pRIPZ between XhoI and EcoRI restriction sites using a homemade Gibson Assembly mix. pLKO shRNAs were prepared by primer annealing and were subsequently cloned into pLKO.1 vector within AgeI and EcoRI sites using T4 DNA ligase (TaKaRa, cat# 2011B) according to the manufacturer’s instructions. Oligonucleotide sequences are provided in Supplementary Table 1.

### Cell culture and transfection

HEK293E cells (human embryonic kidney 293 EBNA1; authenticated by ATCC STR profiling), wild-type and AGO2 knockout HEK293 cells (provided by Honglin Chen), MIHA cells (immortalized hepatocytes; provided by Stephanie Ma), and MHCC97-L cells (hepatocellular carcinoma cell line; provided by Stephanie Ma) were cultured in Dulbecco’s Modified Eagle’s Medium (DMEM) (Thermo, cat# 12100061) supplemented with 10% Fetal Bovine Serum (FBS) (Gibco, cat# A5256701). All cultures were incubated at 37°C in 5% CO_2_ supplemented with humidified air. Cells were seeded at 40–50% confluence the day prior to transfection using GeneJuice Transfection Reagent (Sigma-Aldrich, cat# 70967) or Lipofectamine 2000 Transfection Reagent (Thermo, cat# 11668019) according to the manufacturer’s instructions. Cells were lysed for assays 48 h after transfection.

### RNA extraction and AQ-seq

Following the removal of culture medium, cells were lysed with TRIzol (Thermo, cat# 15596018) according to the manufacturer’s instructions. Libraries were prepared as previously described^77^. In brief, 15 µg of total RNA was size-selected between 17–29 nt on a 15% polyacrylamide urea gel. Purified small RNAs were ligated to a 3′ adaptor (/5rApp/NNNNTGGAATTCTCGGGTGCCAAGG/3ddC/) using T4 RNA ligase 2, truncated KQ (NEB, cat# M0373L) in 20% PEG-8000. After size selection on a 15% polyacrylamide urea gel between 40–55 nt, purified RNA-DNA hybrids were ligated to 5′ adaptor (rGrUrUrCrArGrArGrUrUrCrUrArCrArGrUrCrCrGrArCrGrArUrCrNrNrNrN) using T4 RNA ligase (Thermo, cat# EL0021) in 20% PEG-8000. RNA-DNA hybrids were reverse transcribed using SuperScript III (Invitrogen, cat# 18080044). cDNA libraries were amplified by PCR using Phusion DNA polymerase (NEB, cat# M0530S) in 17–20 cycles with indexed primers. Libraries were size-selected on 6% polyacrylamide gel and purified. Libraries were sequenced on the Illumina Novaseq 6000 by Genewiz.

### AQ-seq analysis

Small RNA-seq adaptor sequences and their associated random sequences were trimmed using cutadapt (-a TGGAATTCTCGGGTGCCAAGG --trimmed-only --quality-cutoff 20 -m 18)^78^. Four random nucleotides were then removed from both ends (-u 4 -u -4), inserts of 18-26 nt were retained (-m 18 -M 26), and trimmed reads were written as FASTA. The insert sequences were mapped onto shRNA sequences using bowtie2 (-f -p 4 --score-min C,0,-1 -k 2)^79^. Multiple mapped reads in SAM files were removed using SAM flags. Next, the filtered files were converted into BED files using samtools and bedtools^80,81^. Subsequently, read counts and positions were obtained using awk, sort, and uniq. During this step, reverse-strand mapped reads were removed, resulting in only forward-strand mapped reads as output.

The read depth for the Round 3 NGS run was 1,024,565 reads for 144 pGIPZ variants (median reads per variant: 6430.5) and 4,409,995 reads for 144 shRNAone variants (median reads per variant: 25833.5). The total read depth for 7 shRNA designs for 144 variants was 22,346,295 reads.

### Validation of guide heterogeneity

HEK293E cells were transfected with plasmids using GeneJuice. After three days, total RNA was extracted using the RNA Miniprep kit (Zymo, cat# R1055) with on-column DNase I treatment. Dumbbell-PCR was performed as previously described by Honda and Kirino^82^. In brief, 500 ng of total RNA was mixed with 10 pmoles of a 5′-dumbbell adaptor mix (5 pmole of adaptor for the shRNA variant and 5 pmole of adaptor for miR-92) in a total volume of 9 µL and incubated at 90°C for 3 min. After removing the tube from the PCR machine, 1 µL of 10x dumbbell annealing buffer (50 mM Tris pH 8.0, 5 mM EDTA, and 100 mM MgCl₂) was added, and the mixture was further incubated at 37°C for 20 min. Next, 1.5 µL of 10x T4 RNA ligase 2 buffer (NEB), 0.5 µL of T4 RNA ligase 2 (NEB, cat# M0239L), 1 µL of UltraPure water (Thermo), and 2 µL of PEG8000 (NEB) were added and incubated at 37°C for 1 hour, followed by an overnight incubation at 12°C. For reverse transcription, 5 µL of ligated RNA and 2 pmole of the variant-specific RT primer were mixed and incubated at 70°C for 2 minutes. A separate reverse transcription reaction was prepared with a miR-92 RT primer for normalization. Maxima H Minus Reverse Transcriptase (Thermo) and its buffer were added to the samples and incubated at 53°C for 1 hr, followed by 85°C for 5 min. TaqMan qPCR was carried out using Premix Ex Taq (Probe qPCR) (Takara, cat# RR390A) in the presence of 0.4 µM TaqMan probe. The expression levels of shRNA variants were normalized to the endogenous level of miR-92.

### Validation of passenger retention

HEK293E cells were transfected with plasmids using the calcium phosphate method. After 48 hours, total RNA was extracted using Trizol. Following DNase I treatment (Takara, cat# 2270B), the RNA was further purified using acidic phenol-chloroform extraction. The guide and passenger strands were reverse transcribed separately using specific stem-loop RT primers as described by Chen et al. and He et al.^83,84^. Since the passenger sequences from shRNAmir and shRNAone were different, we included chemically synthesized 22-nt RNAs as standards. qPCR was performed using TB Green (Takara, cat# RR420A), and the absolute amounts of guide and passenger strands were quantified based on a standard curve obtained from 20 fmole, 2 fmole, and 0.2 fmole of synthetic RNA.

### Plasmid construction and dual luciferase reporter assay

In dual luciferase assays, small RNA targeting sequences were inserted into the 3′ UTR of Firefly luciferase, where inserts were prepared by primer annealing and were cloned into pmirGLO (Promega, cat# E1330) with NheI and SalI restriction sites using T4 DNA ligase (TaKaRa). HEK293E cells were co-transfected with pmirGLO dual luciferase reporter plasmid and pRIPZ plasmid carrying the corresponding targeting site and shRNA accordingly. After 48 h, luciferase activities were assayed with Dual Luciferase Reporter Assay Kit (Vazyme, cat# DL101-01) according to the manufacturer’s instructions. Relative Firefly luciferase activity is normalized to *Renilla* luciferase activity, and relative knockdown is normalized to non-shRNA control.

### Western blotting

Cell lysate was prepared by homemade SDS-lysis buffer [100mM Tris, pH 7.0, 6.7% (v/v) SDS, 20% (v/v) glycerol, 400mM beta-mercaptoethanol, 0.2% (w/v) bromophenol blue] and is subsequently heated at 90°C. Equal volumes of cell lysates were loaded and size-separated on a 12.5% polyacrylamide SDS gel and then transferred onto 0.2 µm nitrocellulose membranes (BIO-RAD, cat# 1620112) using Trans-Blot Turbo Transfer System (BIO-RAD, cat# 1704150). Membranes were blocked in 5% skim milk in PBS-T and subsequently blotted with primary and secondary antibodies with PBS-T washing in between. Commercial antibodies used in this study: anti-AGO2 (Cell Signaling, cat# CST-2897T), anti-GAPDH (Santa Cruz, cat# sc32233), anti-α-Tubulin (Thermo, cat# ab52866), sheep anti-mouse IgG (Cytiva, cat# NA931), and donkey anti-rabbit IgG (Cytiva, cat# NA934).

### Human miRNA shRNAone library design and cloning

pV7 vector was provided by Ralf Jauch and was modified for doxycycline-inducible shRNA expression, where shRNAs were inserted in the 3′ UTR of mCherry (termed pV7-RP).

Non-redundant 18-mers (2nd–19th nt) were extracted from all human miRNA annotated on MirGeneDB 2.1 and inserted into uridine- and adenosine-starting shRNAone backbone (540 shRNAs each). For every 18-mer, three more variants were designed by: (1) miR_7merSupp, flipping central (8th–11th nt with reference to 18-mer) and tail (16th–18th nt with reference to 18-mer) region; (2) miR_7mer, flipping central to tail (8th–18th nt with reference to 18-mer); (3) miR_6mer, flipping central to tail region (7th–18th nt with reference to 18-mer, extending one more flipping region towards the seed region). shRNAs were synthesized as single-stranded oligos, including 100 non-targeting shRNAs, and were PCR amplified with Phanta Super-Fidelity DNA Polymerase (Vazyme, cat# P501) and cloned into pV7-RP using a homemade Gibson Assembly mix. PHGDH shRNA guide sequences were designed based on previous literature^85^ and were cloned into pV7-RP.

### Survival screen on HCC

MHCC97-L cells were infected with a lentiviral shRNA library at MOI < 0.2, ensuring >500-fold coverage. Infected cells were selected with 3 µg/mL puromycin for 48 h. Following selection, cells were treated with 1 µg/mL doxycycline for 5 days to induce shRNA expression. A total of 5 × 10^6^ cells were harvested for pretreatment library preparation, while the remaining cells were divided and treated with either DMSO (control) or 3.25 µg/mL sorafenib for 7 days. Subsequently, 5 × 10^6^ cells were collected from each treatment group. Genomic DNA was extracted using the QIAamp DNA Blood Midi Kit (QIAGEN, cat# 51183), and shRNA sequences were amplified using Phanta Super-Fidelity DNA Polymerase. Amplified shRNA libraries were sequenced on the NovaSeq 6000 platform by Genewiz. Log_2_ fold-changes were calculated using DESeq2^86^. Differential effects between 7merSupp- and 7mer-matched shRNAs were assessed using DESeq2 log_2_ fold changes. For each paired shRNA, the difference in log_2_FC (Δlog_2_FC) and its standard error (propagated from lfcSE) were calculated. Z-scores and two-sided *P* values were computed assuming normality; *P* values were FDR-adjusted (Benjamini–Hochberg).

### CCK-8 cell viability assay

Validation cell lines were treated with 1 µg/mL doxycycline for 7 days to induce shRNA expression. Subsequently, 2,000 cells per well were seeded in 96-well plates in DMEM supplemented with either DMSO (control) or 3.25 µg/mL sorafenib, with at least three technical replicates per condition. After 5 days of incubation, cell viability was assessed by adding 10 µL of CCK-8 reagent to each well and measuring absorbance at 450 nm after 2 h of incubation. Corrected absorbance values were calculated by subtracting the blank absorbance from raw absorbance readings. Relative cell viability was determined by dividing the corrected absorbance of sorafenib-treated samples by that of DMSO-treated samples.

### RNA-seq

Total RNA was isolated from cells using TRIZol and was submitted to Genewiz for RNA sequencing library preparation and sequencing. RNA abundance was quantified using Salmon 1.10.2 using NCBI MANE Select^87^ or Ensembl Canonical transcript as references. Differentially expressed gene analysis and adjusted *P* value calculation were performed with DESeq2^86^.

### Prediction of miRNA target sites

A non-redundant human mRNA reference set was derived from the UCSC hg38 knownCanonical transcript collection (GENCODE v47) and the UCSC foldUtr3 track, both obtained from the UCSC Table Browser on 14 January 2025. The initial table contained 412,034 transcript records. Records lacking an annotated protein-coding gene (knownCanonical.protein) or an annotated 3′ UTR (foldUtr3.seq) were discarded, leaving 22,274 transcripts; transcripts on alternative or variant contigs were then excluded, retaining 19,552 transcripts on the primary chromosomes (chr1–22, X and Y). One transcript per gene symbol was retained, selecting the transcript with the longest 3′ UTR for the 24 genes represented by more than one canonical transcript, yielding 19,528 gene-level entries. Exact sequence duplicates (37) were removed after sorting by gene symbol and keeping the first entry in alphabetical order, leaving 19,491 non-redundant mRNAs. mRNA and 3′ UTR sequences were exported as tab-separated files.

The 19,491 mRNAs were scanned for bipartite miRNA-binding sites with a custom Python script. Two 19-nt miR-494-3p variants (miRBase miR-494; Mir-154-P18 in MirGeneDB v2.1 and Mir-154-P9 in MirGeneDB v3.0) were analysed: miR-494-7merSupp (5′-TGAAACATTGTGGGGATTG-3′) and miR-494-7mer (5′-TGAAACATTGTGCCCTTTG-3′). miRNA nucleotides were numbered 1–19 from the 5′ end, and all target motifs were written as reverse complements (rc) in the mRNA 5′→3′ orientation. Seed motifs were defined as 8mer, rc(m2–m8)+A; 7mer-m8, rc(m2–m8); 7mer-A1, rc(m2–m7)+A; and 6mer, rc(m2–m7), where A is the target nucleotide opposite position 1. A 3′-supplementary target site was defined as the reverse complement of miRNA positions 12–15, 13–16 or 14–17; in that order, the resulting motifs were CCCC, TCCC and ATCC for miR-494-7merSupp, and GGGC, AGGG and AAGG for miR-494-7mer.

Transcripts were searched for a supplementary motif followed by a 1–20 nt spacer and a 6mer seed match in the mRNA 5′→3′ orientation, using the regular expression (SUPPL.{1,20}SEED6) where SUPPL is one of the three supplementary motifs, SEED6 is the 6mer motif (TGTTTC for both variants), and .{1,20} permits 1–20 intervening nucleotides; the outer parentheses capture the complete composite match. For example, the 13–16 window of miR-494-7merSupp was searched with (TCCC.{1,20}TGTTTC). Matching was exact, with no mismatches, G:U wobble or degenerate positions; 8mer, 7mer-m8 and 7mer-A1 seed sites were also recorded separately. Searches used the full-length mRNA sequence.

### RNA extraction and qRT-PCR for mRNA quantitation

Total RNA was isolated using the RNA Miniprep kit (Zymo, cat# R1055) according to the manufacturer’s instructions. Total RNA concentrations were measured with NanoDrop 2000c (Thermo, cat# ND-2000C), and 500 ng total RNA was subjected to reverse transcription using RevertAid RT Reverse Transcription Kit (Thermo, cat# K1691) according to the manufacturer’s instructions. cDNAs were diluted 5-fold and subjected to qPCR with TB Green Premix Ex Taq II (Tli RNase H Plus) (TaKaRa, cat# RR82WR) according to the manufacturer’s instructions on CFX Opus 384 Real-Time PCR System (BIO-RAD, cat# 12011452). qPCR primers are listed in Supplementary Table 1. Relative expression differences were calculated with the 2^-ΔΔCt^ method.

### Establishment of MHCC97-L cell lines stably expressing miRNA variants

shRNAone carrying different guide sequences were cloned into pV7-RP and were subjected to lentivirus preparation. Subsequently, MHCC97-L cells were infected with the corresponding lentivirus at MOI ∼0.3 to ensure single infection in most selected cells.

### Design of human genome-targeting shRNAone sequences and analysis on targeted shRNA screen

The original TRC shRNA library file was retrieved from the Broad Institute. The complementary sequence for positions 2 to 17 of the target sequence was inserted into positions 2 to 17 of the guide strand of shRNAone. The 18th position of the shRNAone guide strand was kept as T, following the original TRC pLKO design. See Supplementary Table 6.

Raw counts were subjected to MAGeCK analysis^88^ for gene-level enrichment analysis and *P* value calculation.

### Plasmid construction and lentivirus preparation for parental reporter MIHA cell line

pLVX plasmid was modified by replacing puromycin N-acetyltransferase with Blasticidin S deaminase. Then, [ER signal peptide; MGWSCIILFLVATATGAHS]-[GFP_1-10_/mNG2_1-10_]-[ER retention signal; KDEL] was cloned into BamHI and XbaI sites. pLVX plasmid was co-transfected with psPAX.2 (Addgene, cat# 12260) and pMD2.G (Addgene, cat# 12259) at a ratio of 10:7:3 into HEK293E using polyethylenimine transfection method. Medium was changed 6 h and was harvested 48 h after transection. Lentiviral soup was concentrated by 100-fold using Lenti-X Concentrator (TaKaRa, cat# 631232) following the manufacturer’s instructions. Wild-type MIHA cells were infected with different viral titres and, after blasticidin selection, cells that survived the highest viral dosage were kept for CRISPR knock-in.

### Split GFP_11_ or mNG2_11_ tag knock-in

Homology repair template were designed as: [500 bp homology arm upstream of stop codon]-[triple alanine]-[split GFP/mNG2_11_ tag(s)]-[ER retention signal; KDEL]-[500 bp homology arm downstream of stop codon] (Supplementary Fig. 15b) and were cloned into pUC19 vector between EcoRI and XbaI sites. sgRNA spacers were designed near the stop codons and were validated prior to knock-in.

Both SpCas9/sgRNA and homology repair template plasmids were co-transfected into corresponding parental MIHA cell lines. Transfected cells were allowed to expand to a 100-mm dish scale. Cells showing the top 5% GFP fluorescent signal were enriched with FACS three times. Subsequently, single-cell clones were isolated to construct the final reporter cell line for siRNA target screening experiments.

After single-cell cloning of reporter cell lines, Sanger sequencing showed that 9 copies of GFP_11_ for *TTR* and 10 copies of mNG2_11_ for *PCSK9* were knocked in (Supplementary Fig. 16).

### Reporter cell line validation for shRNA screening

MIHA reporter cell lines for *TTR* and *PCSK9* were transfected patisiran and inclisiran siRNA mimic (Supplementary Table 1a). siLuc siRNAs were transfected as non-targeting controls. Cells were subjected to FACS for fluorescence quantification on day two post-transfection.

### siRNA target site screening library design and cloning

To design shRNAs tiling the *TTR* (NM_000371.3) transcript, antisense 18-mers were generated at 5-nt intervals from CDS and 3′ UTR and inserted into the uridine-starting shRNAone backbone (110 shRNAs). As a technical replicate, the same 110 antisense 18-mers were inserted into the adenosine-starting shRNAone backbone. Including 100 non-targeting shRNAs, a total of 320 shRNAs were tested in a tiling screen on the *TTR* transcript. For single-nucleotide shRNA tiling screen on *TTR* (NM_000371.3), antisense 18-mers were inserted into uridine-starting shRNAone backbone from the start of annotated transcript to the end and along with 100-non-targeting controls.

For the *PCSK9* (NM_174936.3) transcript, antisense 18-mers were generated at 3-nt intervals from 5′ UTR to 3′ UTR and inserted into uridine-starting shRNAone backbone. Including 100 non-targeting shRNAs, a total of 1,330 shRNAs were tested in a tiling screen on the *PCSK9* transcript.

Pooled shRNA libraries were synthesized as single-stranded oligonucleotides (Twist Biosciences), followed by PCR amplification using Phanta Super-Fidelity DNA Polymerase (Vazyme, cat# P501) and cloning into pRIPZ using a homemade Gibson Assembly mix.

### siRNA target site screening experiments (5-nt and 3-nt shRNA tiling screen on TTR and PCSK9, respectively)

Before starting the screening, reporter cell lines were sorted to obtain a homogeneous population in terms of the fluorescence signals. Five days after sorting, shRNAs were transduced at MOI < 0.2 and selected by puromycin treatment. After 7 days, cells were sorted into 4 bins (bin1: top 10%, bin2: 11–20%, bin3: 21–30%, and bin4: 31–100%) according to the fluorescence intensity.

### crRNA library cloning

CasRx and PspCas13b crRNA libraries were cloned using the pSK027.pXPR-U1-Rfx_DR-stuffer and pSK028.pXPR-U1-Psp_DR-stuffer vectors, respectively. Plasmid backbones were digested with BsmBI (Thermo Fisher Scientific, cat. no. IVGN0136) in the presence of 10 mM DTT at 37°C for 4–5 hours, followed by heat inactivation at 80°C for 20 minutes. Pooled oligonucleotide libraries were synthesized with sublibrary-specific amplification arms. Each sublibrary was first amplified by PCR using its corresponding primer set, followed by purification with PCRClean DX (Aline Biosciences). A second round of PCR was then performed to append sequences required for cloning, followed by another PCRClean DX purification step. Purified inserts were assembled into the digested vectors using a homemade Gibson assembly mix. The resulting plasmid libraries were transformed and expanded to achieve a coverage of at least 100 per guide.

### Constitutive expression plasmid construction

Plasmids for constitutive expression of PspCas13b and CasRx (pSK149.lenti-Cas13b_Psp-NES-HA-P2A-Hygro and pSK148.lenti-CasRx-HA-P2A-Hygro) were generated using the pKW891.lenti-ADAR-dPsp.dHEPN2-msfGFP backbone. The vector was digested with XbaI (Thermo Fisher Scientific, cat. no. IVGN0128) and EcoRI (Thermo Fisher Scientific, cat. no. IVGN0116) at 37°C for 2 hours, followed by heat inactivation at 80°C for 20 minutes. The coding sequences of PspCas13b and CasRx, together with the hygromycin resistance cassette, were PCR-amplified from pJK712.pV7-PspCas13b-HA-HygroR and pJK711.pV7-RfxCas13d-HA-HygroR, respectively. Amplified fragments were assembled into the digested backbone using a homemade Gibson assembly mix to generate the final constructs.

### Stable cell-line preparation for PspCas13b and CasRx

Cell lines expressing PspCas13b or CasRx were generated by lentiviral transduction. Lentiviral particles were produced from a single batch and concentrated 100-fold using Lenti-X Concentrator (TaKaRa, cat. no. 631232) according to the manufacturer’s instructions. MIHA-WT and MIHA-reporter cells were infected in parallel with varying volumes of the same viral preparation to achieve a range of viral titers (MOI >1).

Following transduction, cells were subjected to 150 µg/mL hygromycin (Invivogen, cat. no. ant-hg-1) selection for approximately four passages to enrich for transduced populations. Multiple infection conditions were evaluated, and cells that survived higher viral input conditions were retained. These cells were subsequently cultured for an additional 7–10 days to allow recovery, followed by a second round of hygromycin selection to further stabilize the population.

### Pooled screening setup for 1-nt tiling libraries

Reporter cell lines were transduced with the corresponding shRNA, PspCas13b crRNA or CasRx crRNA libraries under pooled screening conditions at an MOI of <0.5. Following transduction, cells were cultured for 1 day and selected with 1 μg/mL puromycin (GoldBio, cat. no. P-600-100) for 3 days until non-infected cells were eliminated. For the shRNA and PspCas13b screens, cells were subsequently recovered for 5 days before GFP-based knockdown assessment. For the CasRx screen, cells were recovered for 1 day, after which half of the population was analyzed by FACS and the remainder was cultured until day 5 before assessment.

### FACS-based assessment of knockdown

After recovery, knockdown efficiency was assessed by GFP fluorescence using flow cytometry. Cells were treated with DNase I (Takara) and 5 mM MgCl_2_ (Thermo) for 15 min and then resuspended in PBS containing 2% FBS, 2mM EDTA, and 2 mM NaN_3_ for flow cytometry analysis, and sorted on BD FACSAriaTM SORP and BD FACSAriaTM Fusion cell sorters. For each screen, two GFP-positive populations were collected: bin 1, comprising the lowest 15% of GFP-positive cells, and bin 4, comprising the top 60%. Genomic DNA was extracted from each bin, integrated guide cassettes were amplified, and the products were sequenced on an Illumina platform to quantify the abundance of each library member in the low- and high-GFP fractions.

### Random positional shuffling for cluster definition

To objectively define transcript regions enriched with high-efficiency target sites, we applied a rolling-mean random positional shuffling framework to the shRNAone log_2_FC profiles of *TTR* and *PCSK9*. Rolling means were computed with a symmetric window of 21 nt (converted to array steps by dividing by the median spacing between adjacent guides: 1 nt for *TTR*, 3 nt for *PCSK9*). The shRNAone log_2_FC values were randomly positionally shuffled 1,000 times, the rolling mean was recomputed for each permutation, and the 99.5th and 0.5th percentiles were taken as empirical null bounds. Target sites exceeding the upper bound were classified as optimal, those below the lower bound as poor, and the remainder as background. Accessibility values (log_10_-transformed) under both parameter sets were compared across the three categories using Wilcoxon rank-sum tests, and the ability of accessibility to discriminate optimal from non-optimal sites was assessed by ROC analysis with the Youden-index threshold reported.

### Gaussian smoothing of knockdown profiles

To enable qualitative comparison of positional knockdown patterns across different RNA-targeting systems, raw log₂ fold-change (log₂FC) values were smoothed along transcript coordinates using Gaussian kernel. Guide positions were defined by their nucleotide start sites along the *TTR* transcript (NM_000371.4). Smoothing was performed with a Gaussian filter, in which each data point is replaced by a weighted average of neighboring values.

### Offset alignment of knockdown profiles

To compare positional knockdown patterns across different RNA-targeting systems, we performed an offset-alignment analysis using shRNAone and PspCas13b (mean of biological replicates) and cytolytic CasRx (day 5 dataset). For each pairwise comparison, we evaluated the correlation between raw data points, not the Gaussian smoothed trend lines, by applying a series of integer-nucleotide offsets to one profile while keeping the other fixed. At each offset, we identified overlapping positions between the two profiles and quantified their similarity using the Pearson correlation coefficient of the aligned log₂FC values. The optimal offset was defined as the shift that maximized the Pearson correlation. Offsets are given in nucleotides and are defined along the target transcript using the guide coordinate system: positive offsets correspond to shifts towards the 3′ end of the target, and negative offsets to shifts towards the 5′ end.

### Accessibility parameter screen and optimal parameter identification

To identify the RNAplfold parameter set most predictive of knockdown efficacy, we performed a combinatorial grid search over the three thermodynamic parameters that govern local unpaired probability: the sliding window size (*W*, 20–200 nt), the maximum base-pairing distance (*L*, from 0.25*W* to 0.75*W*), and the unpaired region length (*U*, 2–30 nt depending on guide length). For each of approximately 8,000 (*W*, *L*) combinations, an RNAplfold_lunp file was generated on the *TTR* transcript (NM_000371.4, wild-type sequence) with a fixed per-nucleotide constraint *U* = 30. At each (*W*, *L*, *U*) combination, the unpaired probability at every experimental target site was extracted under each of four direction definitions: 3prime (pos + *Lg* − 1), 5prime (pos + *U* − 1), upstream (pos − 1), and downstream (pos + *Lg* − 1 + *U*), where pos is the 1-based start position and *Lg* the guide length. Spearman correlations with experimental log_2_FC were then computed for the Whole transcript, CDS, and 3′ UTR regions.

To determine which direction most reliably captured the relationship between accessibility and knockdown, all configurations with positive Spearman *r* were retained and corrected for multiple testing (Benjamini–Hochberg, FDR < 0.05). Two analyses were performed on the FDR-significant subset, focusing on the Whole transcript region for final presentation. First, the proportion of top-5% configurations belonging to each direction was computed, quantifying the extent to which the best-performing parameter space was dominated by one direction. Second, a linear mixed-effects model (LMM) was fitted with direction × region as fixed effects and random intercepts for *W*, *L*, and *U*; pairwise contrasts between 3prime and each alternative direction were extracted with estimated marginal means, and the Whole-region contrasts are reported. Based on convergent evidence from both analyses, the 3prime direction was selected for all subsequent work. Separately, the relative non-linear importance of *W*, *L*, and *U* was assessed by random forest permutation importance (ranger, 500 trees), quantified as the relative increase in mean squared error upon permuting each parameter.

With direction anchored to 3prime and *W* and *L* provisionally fixed to the RNAxs reference (*W* = 80, *L* = 40), the unpaired-region length *U* was varied across its valid range for the Whole transcript region. For each effector, the *U* plateau was defined as the interval in which Spearman *r* exceeded 75% of its maximum value, yielding tool-specific constraints: shRNAone *U* = 6–14, PspCas13b *U* = 9–23, and CasRx *U* = 8–14. Within these plateaus, all (*W*, *L*) combinations (*W* = 20–200, *L* = 0.25*W*–0.75*W*, constrained by *W* ≥ *L*) were evaluated by computing the mean Spearman *r* across the available *U* values. For each combination, a weighted score was constructed (0.50 × PspCas13b + 0.30 × shRNAone + 0.20 × CasRx). A 5 × 5 sliding-window block analysis was applied to the resulting (*W*, *L*) landscape, and spatial non-maximum suppression (minimum separation ≥ 5 units in *W* or *L*) was used to select the top 20 spatially independent candidate blocks. The final ranking of these candidates was determined by a composite score integrating three performance metrics: the weighted block score, the first quartile of Spearman *r* across *U* = 6–16 on a published siRNA efficacy dataset (2,327 siRNAs, of which 122 were classified as functional and 121 as non-functional after excluding intermediate-efficacy constructs)^61^, and the first quartile of binary AUC on the same dataset. The composite score was computed as a robust Z-score average, with the mean and standard deviation of each metric estimated exclusively from the 20 candidates (the reference *W*80_*L*40 point was excluded from the scaling to prevent it from distorting the normalization). (*W* = 35, *L* = 20) was retained as the optimal parameter set. At *W*35_*L*20, the *U* values yielding maximal correlation in the Whole region were shRNAone *U* = 7, PspCas13b *U* = 11, and CasRx *U* = 11. The corresponding values under the RNAxs reference *W*80_*L*40 were shRNAone *U* = 7, PspCas13b *U* = 17, and CasRx *U* = 9. Both parameter sets were carried forward for validation.

### Validation of the optimal parameters

Tafer et al.^60^ derived the original W80_L40_U8 parameters for siRNA from the Huesken et al.^61^ siRNA dataset (n = 2,433; our parameter optimization used the 2,327-siRNA subset of this dataset) and generated an independent set of 360 siRNAs to evaluate their own model. We evaluated our W35_L20_U7 model for shRNAone on this independent set (n = 360) by computing Spearman’s r between predicted accessibility and measured knockdown.

Hu et al.^62^ generated PspCas13b tiling data on mCherry and BCR-ABL1 (n = 117). Because efficient and precise U6-driven expression requires a purine at the transcription start site^63^, and the starting nucleotide is not fixed across the guides in this dataset, we additionally analyzed the subset of purine-starting guides (n = 54) separately.

### Circular shift for correlation testing

To test whether the observed accessibility–log_2_FC correlations could arise spuriously from the intrinsic spatial autocorrelation of both signals, we performed circular shift. At each comparison (external model at best offset; self-derived accessibility–wet data for *TTR* and *PCSK9* under both parameter sets), the experimental log_2_FC vector was circularly shifted by a random offset (sampling uniformly from 1 to n − 1 positions), preserving its autocorrelation structure while breaking point-to-point correspondence. The Spearman correlation was recomputed for each of 1,000 such shifts, and the empirical two-sided *P* value was calculated as the fraction of shifted absolute *r* values exceeding the observed absolute *r*.

### Accessibility-based knockdown prediction and cross-system validation

For each effector and each of the two final parameter sets, a dedicated RNAplfold_lunp file was generated for the *TTR* transcript (*U* = 30), and the 3prime accessibility at each experimental target site was extracted and merged with the cleaned log_2_FC values. Spearman correlations were computed within the Whole, CDS, and 3′ UTR regions, and accessibility was compared between the top and bottom 20% of target sites ranked by log_2_FC using Wilcoxon rank-sum tests. The same procedure was applied to the *PCSK9* transcript (NM_174936.3, shRNAone data at 3-nt resolution), with correlations computed across Whole, 5′ UTR, CDS, and 3′ UTR regions.

We compared the two self-derived parameter sets with seven published prediction models, evaluated on the *TTR* transcript at each model’s optimal alignment offset (determined by scanning −50 to +50 nt shifts relative to experimental data and selecting the shift yielding the highest Spearman correlation). For CasRx, active models included Sanjana2020 (23 nt), TIGER2023 (23 nt), DeepCas13 (23 nt), and Hsu2023 (30 nt). For PspCas13b, models included Sanjana2020 (23 and 30 nt), DeepCas13 (30 nt), Hsu2023 (30 nt), and Fareh2024 (30 nt). Spearman correlations with experimental log_2_FC were computed at the best offset for the Whole, CDS, and 3′ UTR regions. Self-derived parameters were not evaluated for shRNAone, as the accessibility model was developed using the shRNAone grid-search data and would therefore not represent an independent comparison.

### Genome-wide PspCas13b guide design

For genome-wide PspCas13b guide design, protein-coding transcripts were extracted from MANE RefSeq (MANE.GRCh38.v1.5) using Biopython. RNAplfold_lunp files were computed for each transcript under both parameter sets (*U* = 30), parallelized across 50 CPU cores. A 30-nt sliding window matching the PspCas13b spacer length was applied to each transcript, and 3prime accessibility was extracted under the corresponding *U* value for each parameter set (*U* = 11 for *W*35_*L*20, *U* = 17 for *W*80_*L*40). Within each transcript, target windows were ranked by descending accessibility, yielding a queryable genome-wide resource of pre-scored PspCas13b guides.

### Statistics and reproducibility

The statistical details are provided in the figure legends.

## ACKNOWLEDGEMENTS

We thank Narry Kim and S. Chan Baek for the initial support of the project idea and Stephanie Ma for kindly providing MIHA and MHCC97-L. We also thank Johnson Ng, Joe Lam, Ezra Cheng, and Jason Li for helpful discussions. We appreciate the staff of the Centre for PanorOmic Sciences (CPOS) in the LKS Faculty of Medicine at the University of Hong Kong for their technical assistance. This work was supported by the Research Grants Council (ECS/GRF #27112522, #17118324, and #17118724 to S.C.K.; GRF #17106622, #17117925, and #1710112012 to R.J.) and the University of Hong Kong (Start-up fund and Platform Technology Funding to S.C.K.).

## CONTRIBUTIONS

J.K. and S.C.K. conceived and developed the idea. J.K., S.S.K.L., and C.F.L. performed experiments. G.W., J.K., and S.C.K. carried out bioinformatics. B.W.Y.M. and H.C. generated AGO2 knockout cells. M.W. and R.J. made a dox-inducible lentiviral vector for efficient miRNA expression. J.K., G.W., S.S.K.L., and S.C.K. analyzed the data and wrote the manuscript.

## DATA AVAILABILITY

The datasets have been uploaded to NCBI GEO (Gene Expression Omnibus).

To review GEO accession GSE310822:

Go to https://www.ncbi.nlm.nih.gov/geo/query/acc.cgi?acc=GSE310822 Enter token izgteweofpsjpel into the box

To review GEO accession GSE310823:

Go to https://www.ncbi.nlm.nih.gov/geo/query/acc.cgi?acc=GSE310823 Enter token uhypwuyujxmrvit into the box

To review GEO accession GSE310827:

Go to https://www.ncbi.nlm.nih.gov/geo/query/acc.cgi?acc=GSE310827 Enter token ivyrcskqpjobrwh into the box

The codes and plasmid maps are available on GitHub (https://github.com/ChulLab/shRNAone). During revision, we will deposit the code to Zenodo.

The website for PspCas13b guide RNA designs for the human transcriptome is available (https://cas13b.chullab.com).

## SUPPLEMENTARY TABLE

Supplementary Table 1. Oligonucleotide sequences.

Supplementary Table 2. shRNA sequences and raw counts from Rounds 1 and 2.

Supplementary Table 3. shRNA sequences and raw counts from Round 3.

Supplementary Table 4. SynthoMir sequences, raw counts, pair-wise comparison from sorafenib resistance screens, and DEG analysis from RNA-seq.

Supplementary Table 5. miRNA nomenclature tables from MirGeneDB 2.1, MirGeneDB 3.0, and miRBase.

Supplementary Table 6. TRC-like shRNAone sequences targeting all human, mouse mRNAs, and controls.

Supplementary Table 7. shRNA sequences, raw counts, and MAGeCK analysis from targeted shRNA screens.

Supplementary Table 8. shRNA sequences, raw counts, and log2FC from shRNAone 5-nt and 1-nt tiling TTR knockdown screens.

Supplementary Table 9. shRNA sequences, raw counts, and log2FC from shRNAone 3-nt tiling PCSK9 knockdown screens.

Supplementary Table 10. crRNA sequences, raw counts, and log2FC from CasRx 1-nt tiling TTR knockdown screens.

Supplementary Table 11. crRNA sequences, raw counts, and log2FC from PspCas13b 1-nt tiling TTR knockdown screens.

## Supplementary Information

**Supplementary Fig. 1:**
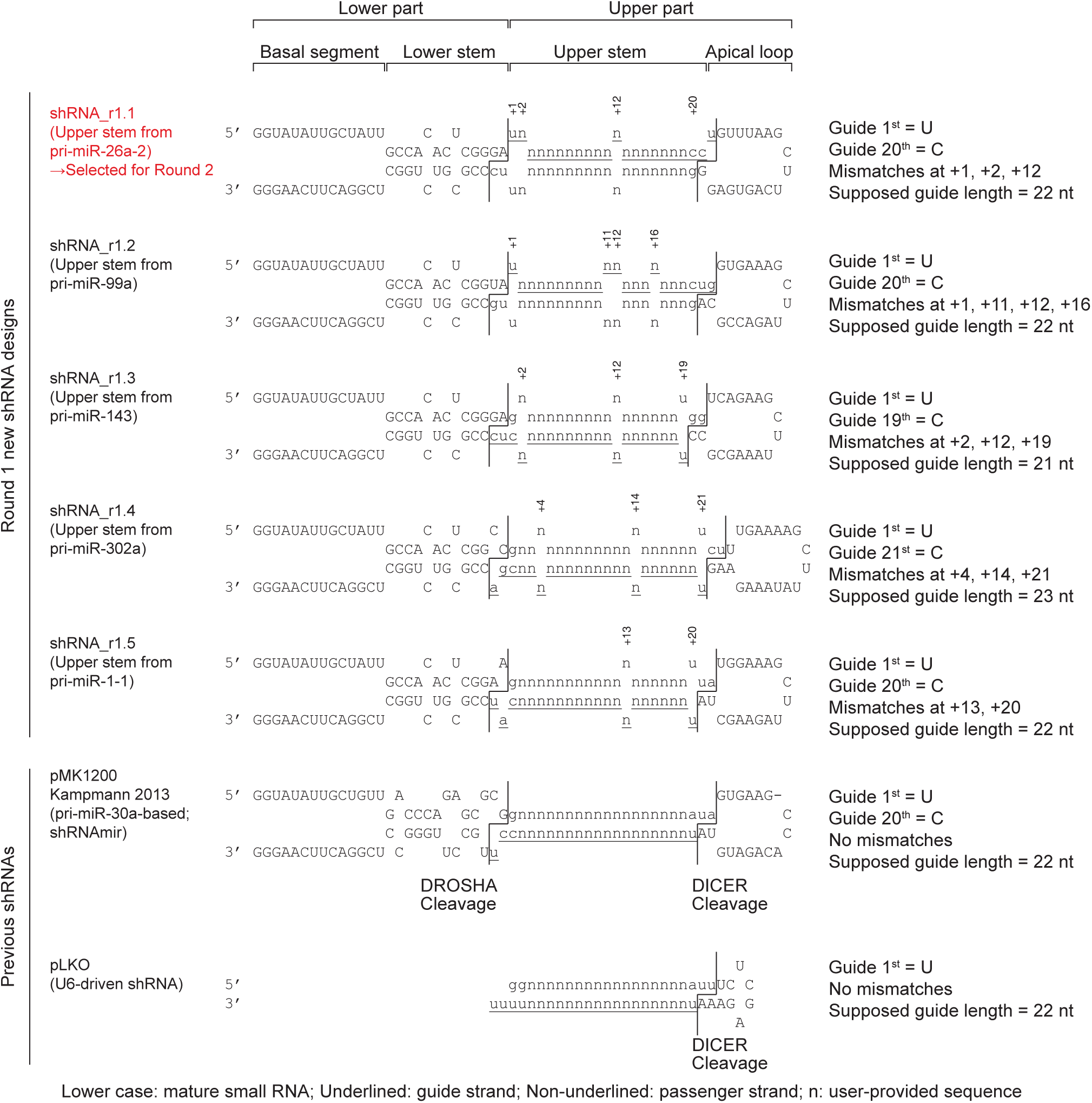
shRNA backbones tested in first round of shRNA optimization. Structures of the five candidate backbones (shRNA_r1.1–shRNA_r1.5) and two reference backbones, pMK1200 (a pri-miR-30a shRNAmir)^32^ and pLKO (U6-driven simple hairpin). Candidate backbones are derived from pri-miR-26a-2 (r1.1), pri-miR-99a (r1.2), pri-miR-143 (r1.3), pri-miR-302a (r1.4) or pri-miR-1-1 (r1.5). Predicted DROSHA and DICER cleavage positions are marked. Red, shRNA_r1.1 for second round optimization; lower case, mature small RNA; underlined, guide strand; not underlined, passenger strand; n, user-defined target-complementary nucleotide.

**Supplementary Fig. 2:**
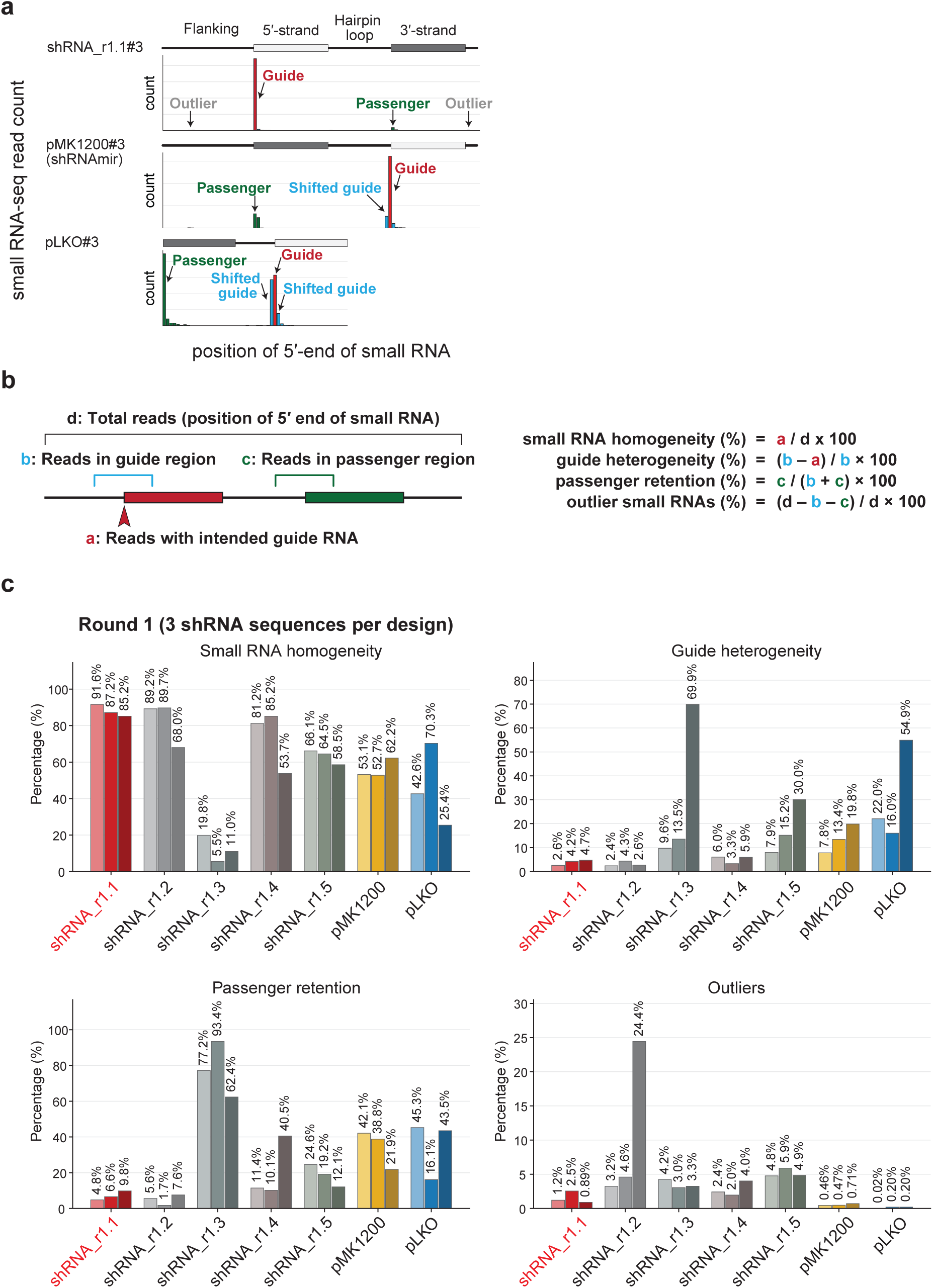
Small RNA-seq metrics and round 1 of shRNA optimization. **a** Small RNA-seq read count mapped with 5′ end position for shRNA_r1.1#3, pMK1200#3 and pLKO#3 expressed in HEK293E cells. Reads are colored as intended guide (red), passenger (green), shifted guide (blue) or outlier small RNA (grey). **b** Definition of the four metrics. a (red), reads with the intended guide 5′ end; b (blue), all reads in the guide region; c (green), all reads in the passenger region; d, all reads mapping to the construct. **c** Guide heterogeneity, passenger retention and outlier small RNAs for shRNA_r1.1–r1.5, pMK1200 and pLKO. Each cross is one independent insert sequence (n = 1).

**Supplementary Fig. 3:**
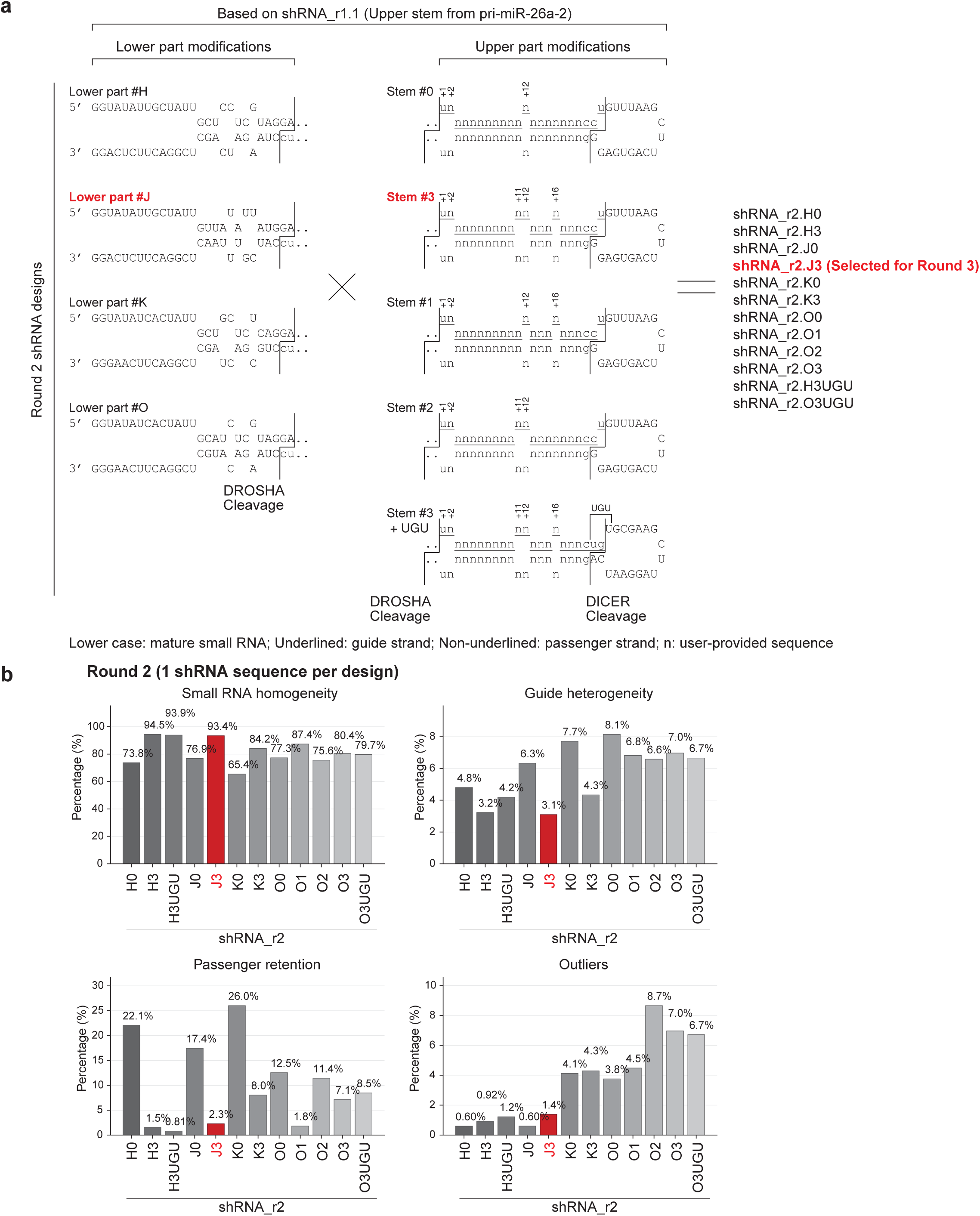
shRNA backbones tested and metrics in round 2 of shRNA optimization. **a** Structures of the upper and lower stems screened, generated by combining 4 lower stem variants (#H, #J, #K, #O) and 5 upper stem variants: shRNA_r2.H0, r2.H3, r2.J0, r2.J3, r2.K0, r2.K3, r2.O0, r2.O1, r2.O2, r2.O3, r2.H3UGU and r2.O3UGU. Selected combination (lower part #J, Stem #3) and the resulting backbone shRNA_r2.J3 are highlighted in red. Annotation as in Supplementary Fig. 1. **b** Small RNAseq metrics for the 12 backbones. Each point is a single insert sequence (n = 1).

**Supplementary Fig. 4:**
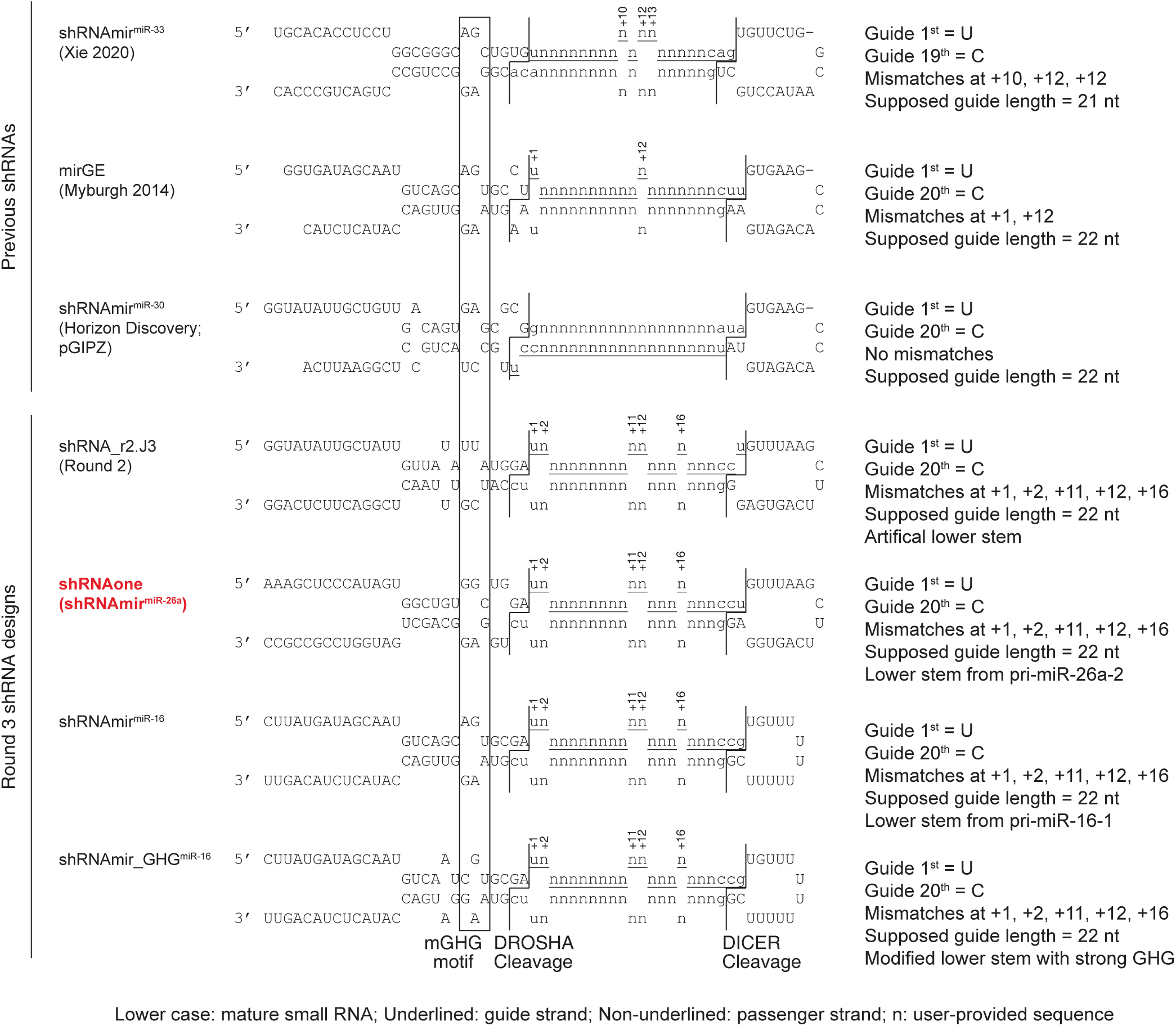
shRNA backbones tested in round 3 of shRNA optimization. Structures of 4 candidate backbones alongside three previously published shRNAmir scaffolds^15,16^ used for comparison: shRNAmir^miR-33^, mirGE, and shRNAmir^miR-30^ (pGIPZ; Horizon Discovery). Round 3 designs comprise shRNA_r2.J3, shRNAmir^miR-26a^ (shRNAone; red), shRNAmir^miR-16^ and shRNAmir_GHG^miR-16^. Annotation as in Supplementary Fig. 1.

**Supplementary Fig. 5:**
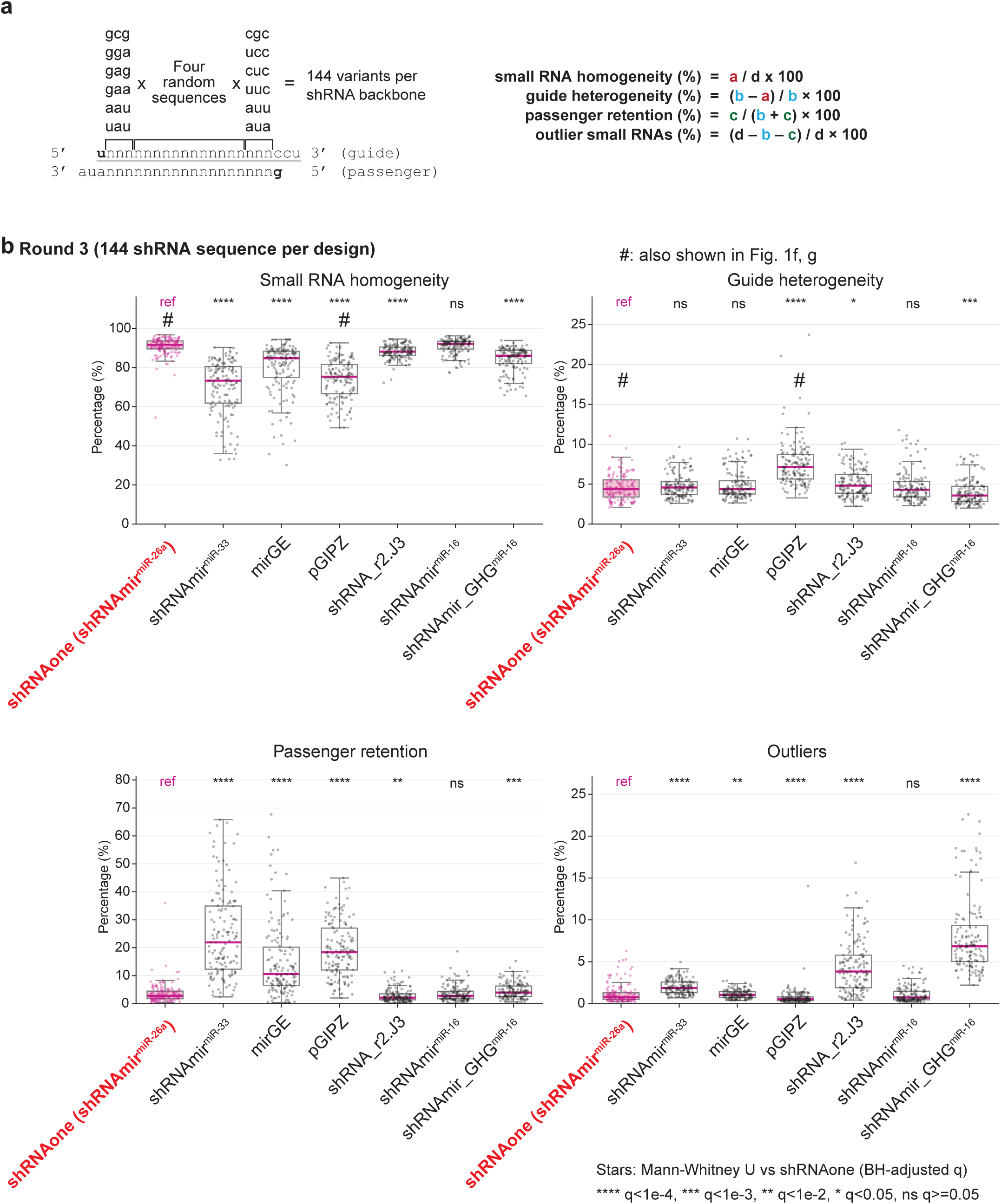
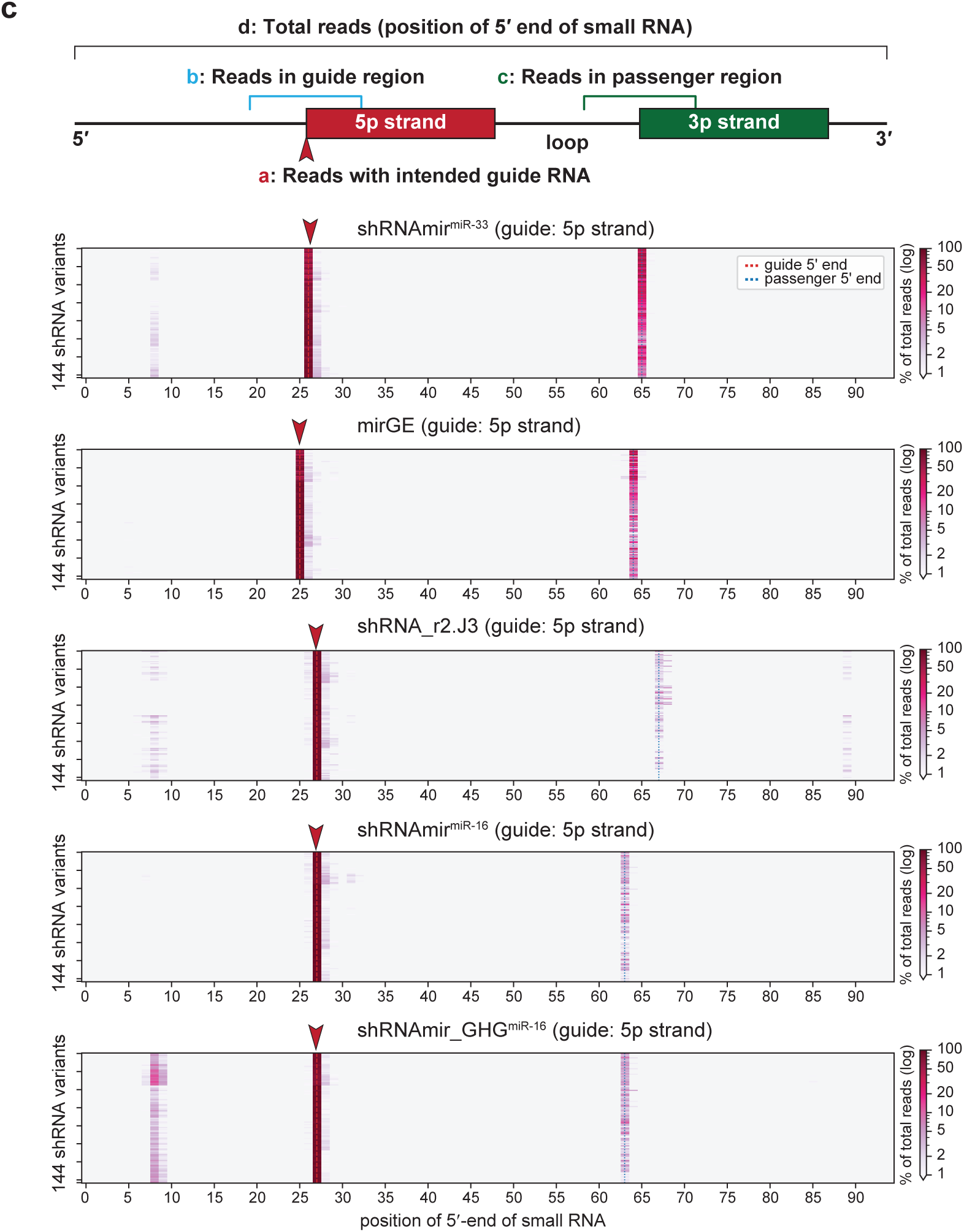
shRNA library design and metrics in round 3 of shRNA optimization. **a** Benchmarking shRNA library design. A matrix of 144 inserts (4 random middle sequences × 6 guide 5′ triplets [gcg, gga, gag, gaa, aau, uau] × 6 guide 3′ triplets [cgc, ucc, cuc, uuc, auu, aua]) was cloned into each backbones, transfected into HEK293E cells, and profiled by small RNA-seq. **b** Small RNA homogeneity, guide heterogeneity, passenger retention, and outlier small RNAs for the seven backbones. Each dot is one sequence variant; horizontal line and whiskers show mean ± sd. shRNAone (shRNAmir^miR-26a^) is highlighted in red. Samples marked with # are also shown in Fig. 1f, g. **c** Heatmap of small-RNA 5′ end positions for the 144 shRNA variants expressed from other designs. Each row is one variant and each column is a 5′ end position of the mapped reads. Color indicates the percentage of a variant’s total reads starting at that position (each row sums to 100%) on a log scale from 1% to 100%. Dashed red and dotted blue vertical lines mark the intended guide and passenger 5′ ends, respectively.

**Supplementary Fig. 6:**
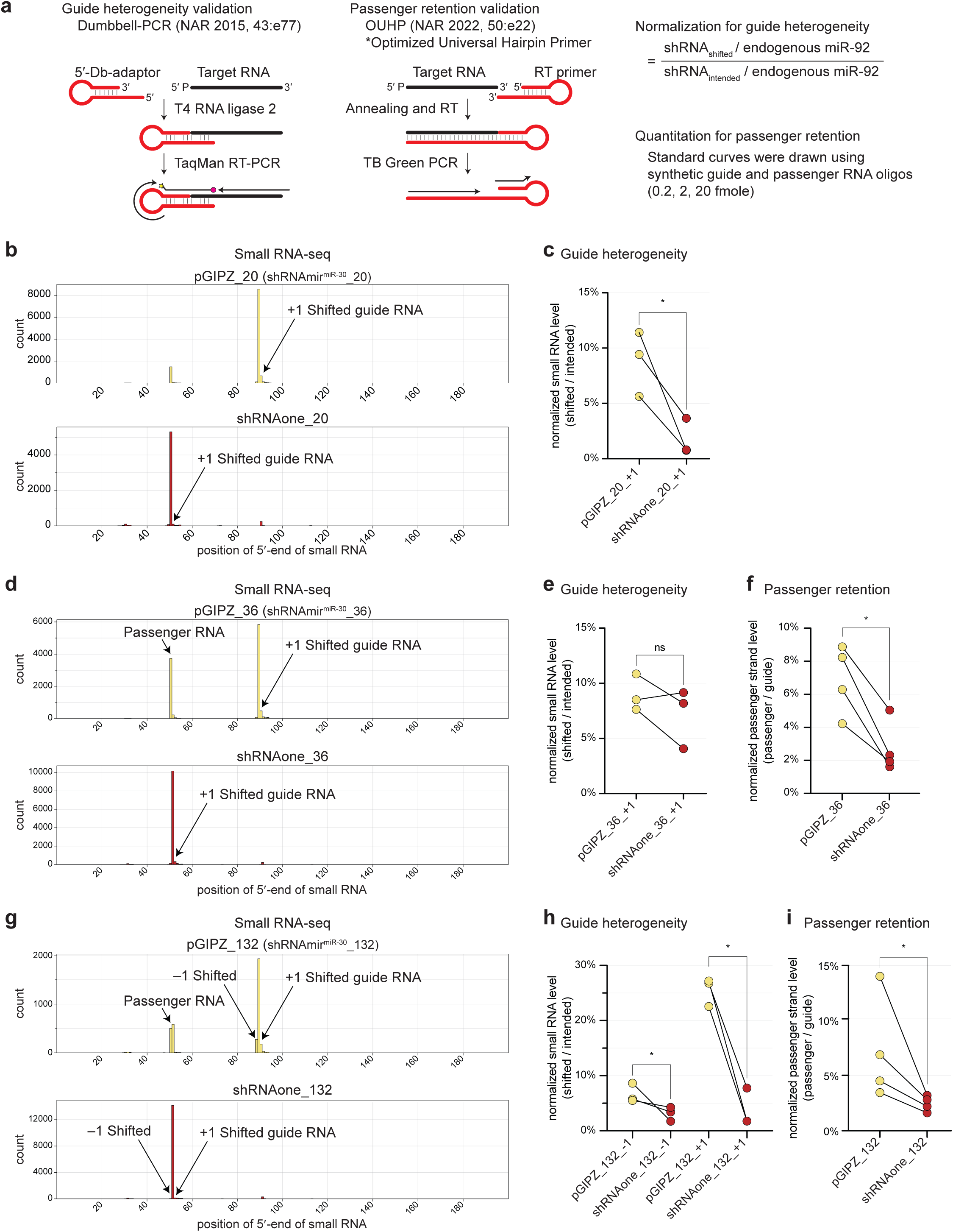
qRT-PCR validation of reduced guide heterogeneity and passenger retention for shRNAone. **a** Validation schematics and normalization steps. Metrics were quantified against standard curves generated from synthetic guide and passenger RNA oligonucleotides (0.2, 2, and 20 fmol). **b**, **d**, **g** Small RNA-seq read count mapped with 5′ end position for reads #20 (b), #36 (d) and #132 (g) expressed from pGIPZ (shRNAmir^miR-30^; top) or shRNAone (bottom). Peaks corresponding to +1- and −1-shifted guide RNAs and to the passenger RNA are indicated. **c**, **e**, **h** qRT-PCR quantification of guide heterogeneity, expressed as normalized small RNA level (shRNA_shifted/shRNA_intended), for the +1-shifted species of #20 (c) and #36 (e) and for the −1- and +1-shifted species of #132 (h) (n = 3). **f**, **i** qRT-PCR quantification of passenger retention, expressed as normalized passenger strand level (passenger/guide), for #36 (f) and #132 (i) (n = 4). In c, e, f, h, and i, connected pairs of points are matched replicates (pGIPZ, yellow; shRNAone, red). One-tailed Student’s t-test; *P < 0.05; ns, not significant.

**Supplementary Fig. 7:**
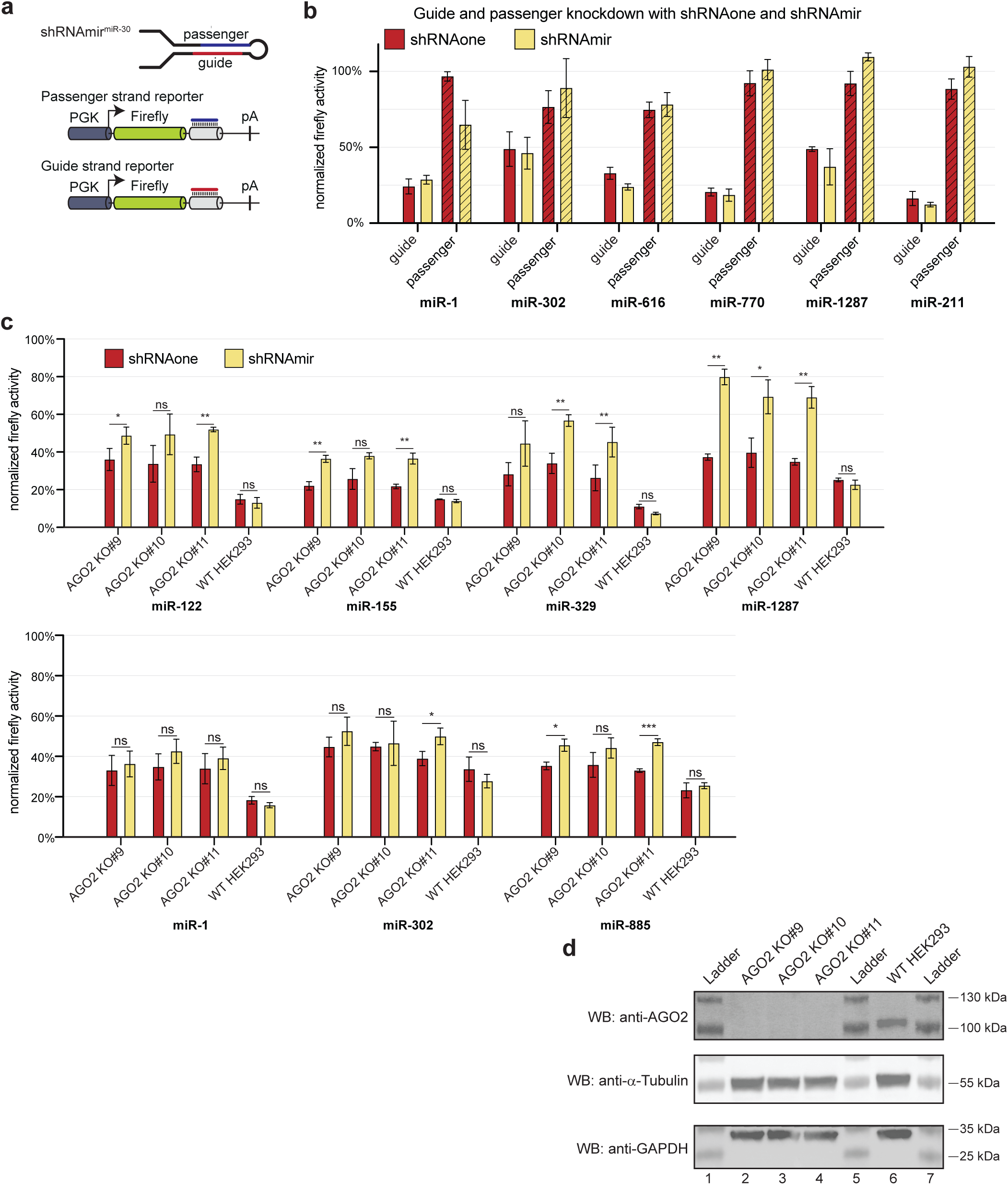
Luciferase reporter validation of reduced guide heterogeneity and passenger retention for shRNAone. **a** Schematic of the shRNAmir^miR-30^ hairpin (guide, red; passenger, blue) and of the PGK-driven firefly luciferase reporters carrying a fully complementary passenger-strand or guide-strand target site. **b** Normalized firefly activity in wild-type HEK293 cells for six guide sequences (miR-1, miR-302, miR-616, miR-770, miR-1287, miR-211) expressed from shRNAone (red) or shRNAmir (yellow). Solid bars, guide-strand reporter; hatched bars, passenger-strand reporter. **c** Normalized firefly activity of the guide-strand reporter in three independent AGO2-knockout HEK293 clones and in wild-type HEK293 cells, for seven guide sequences (top: miR-122, miR-155, miR-329, miR-1287; bottom: miR-1, miR-302, miR-885), annotation as in b. **d** Western blot confirming loss of AGO2 protein in the knockout clones. Membranes were probed for AGO2, α-tubulin and GAPDH; molecular-mass markers (kDa) are indicated. Data in b and c are mean ± sd (n = 3). Student’s t test; *P < 0.05; **P < 0.01; ***P < 0.001; ns, not significant.

**Supplementary Fig. 8:**
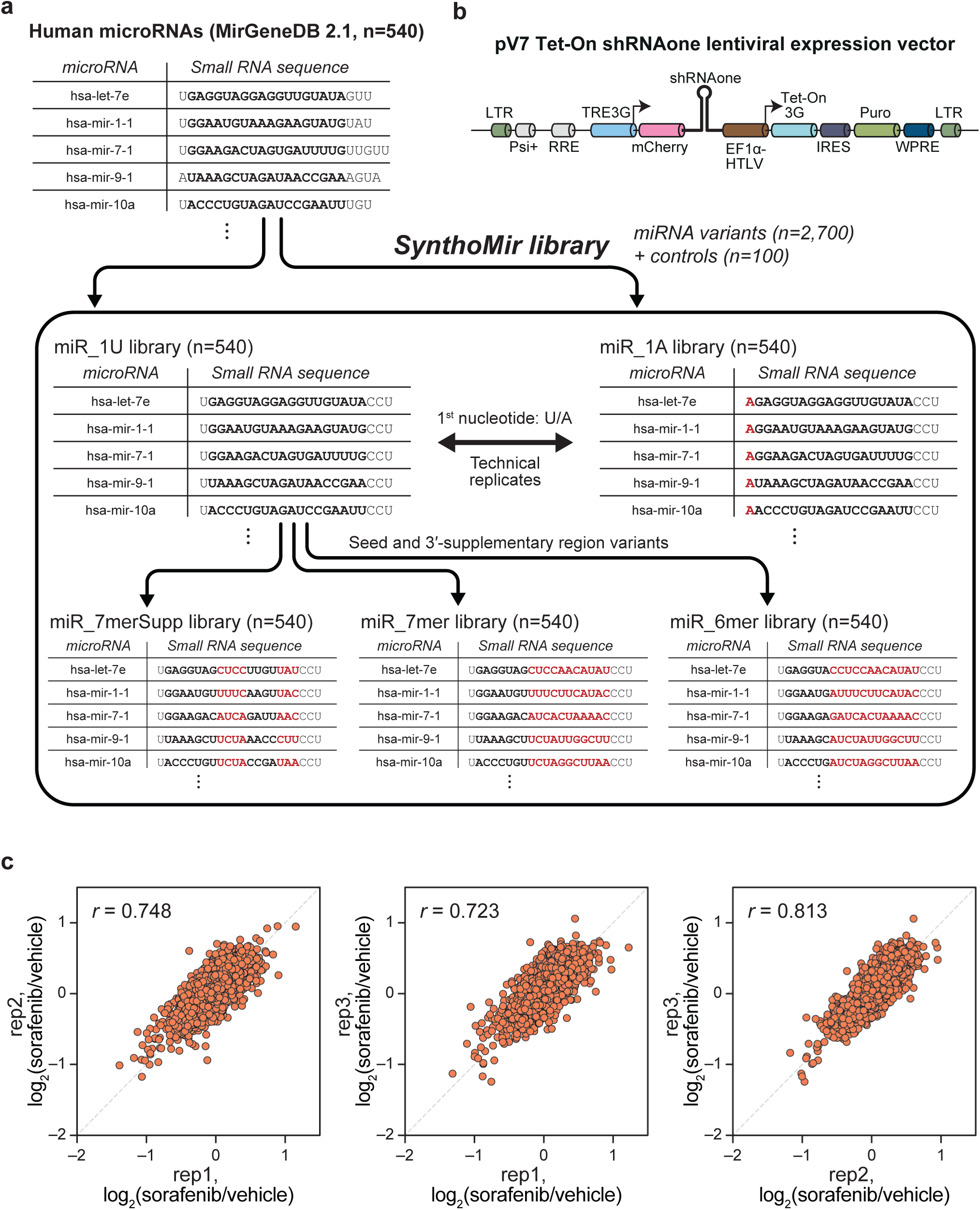
Design and reproducibility of the SynthoMir library. **a** Schematics for SynthoMir library design. All annotated human miRNAs with unique sequences in the 2nd–19th nt in MirGeneDB 2.1 (n = 540) were used to construct the miR_1U library. The miR_1A library (n = 540) differs only at the 5′-terminal nucleotide (U to A, red) from miR_1U. Three variant libraries, miR_7merSupp, miR_7mer, and miR_6mer, contain flipped nucleotides in specific positions (n = 540 each; flipped nucleotides in red). With 100 non-targeting control constructs, SynthoMir comprises 2,800 shRNAs in total. **b** Map of the pV7 Tet-On shRNAone lentiviral expression vector. LTR, long terminal repeat; Psi+, packaging signal; RRE, Rev response element; WPRE, woodchuck hepatitis virus post-transcriptional regulatory element. **c** Pairwise comparison of shRNA-level log2(sorafenib/vehicle) enrichment between the three biologically independent replicates of the SynthoMir screen in MHCC97-L cells. Pearson’s r = 0.748 (rep1 vs rep2), 0.723 (rep1 vs rep3) and 0.813 (rep2 vs rep3).

**Supplementary Fig. 9:**
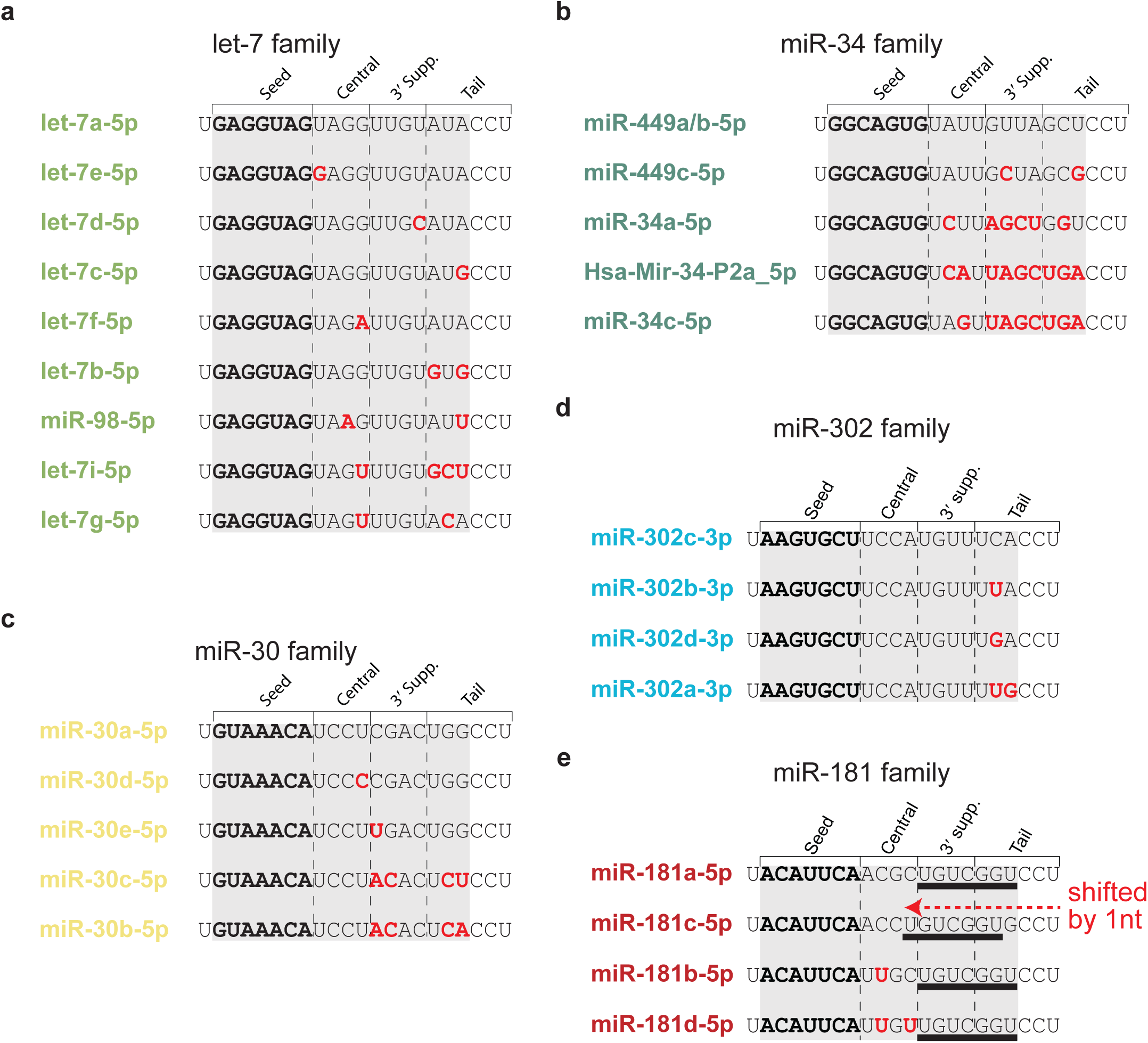
Individual miR_1U shRNA guides in let-7, miR-34, and miR-30 families. **a–e** Aligned mature sequences of the let-7 family (a), the miR-34 family (b), the miR-30 family (c), the miR-302 family (d), and the miR-181 family (e). Vertical dividers segregate the seed, central, supplementary and tail regions. Bold, seed sequences; red, nucleotides differing from the reference family member; shaded, customizable region in shRNAone; horizontal bars, the 3′-supplementary and tail block in miR-181 family, displaced by one nucleotide towards the 5′ end in miR-181c (red arrow).

**Supplementary Fig. 10:**
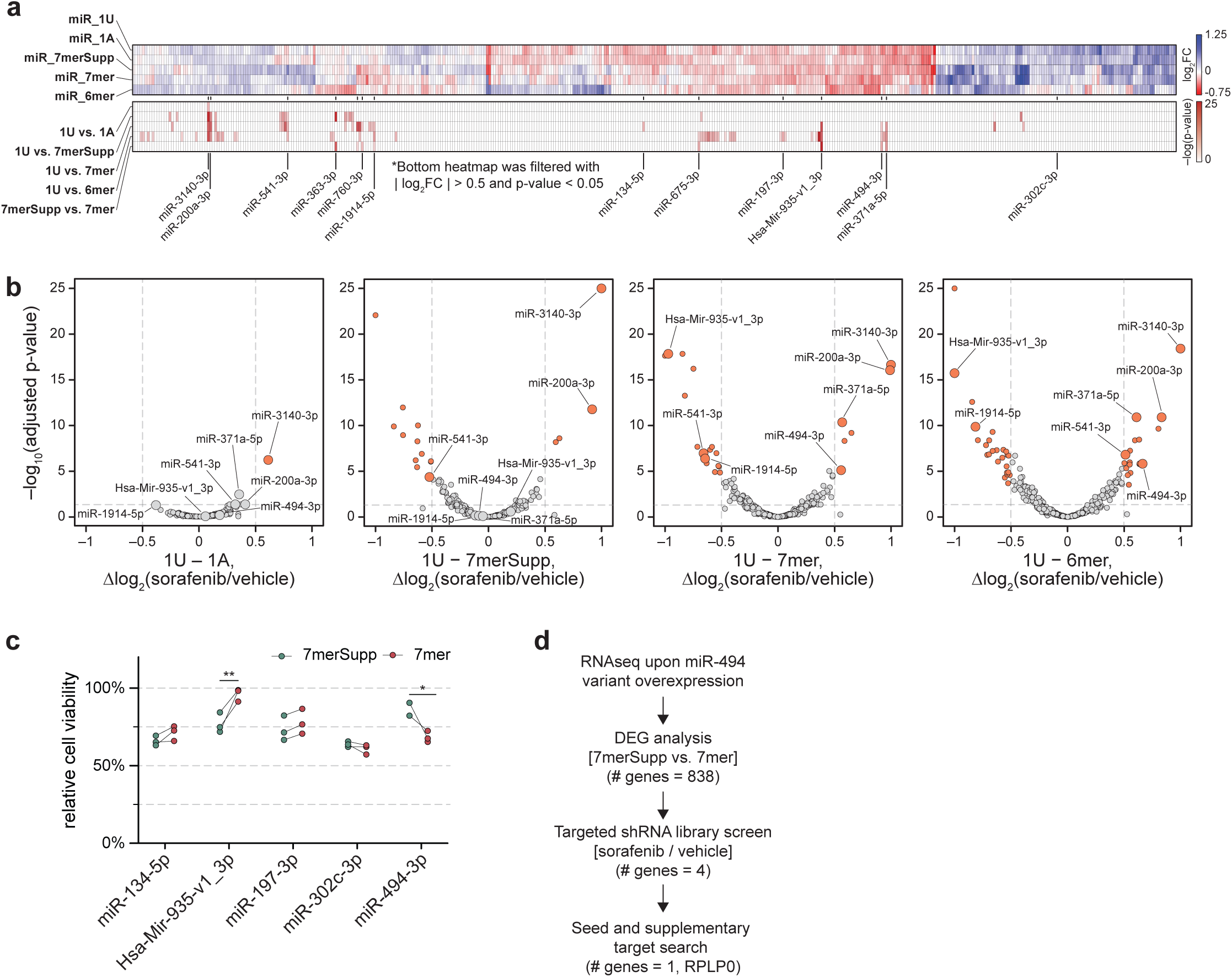
3′-supplementary pairing by miR-494-3p represses RPLP0 and confers sorafenib resistance in hepatocellular carcinoma cells. **a** Pooled screen with SynthoMir in MHCC97-L cells. Top, log2(sorafenib/vehicle) for each library miRNA (columns, identically ordered in both heatmaps) across the five pairing architectures (rows): miR_1U, miR_1A, miR_7merSupp, miR_7mer and miR_6mer. Blue, enrichment under sorafenib; red, depletion. Bottom, significance (−log P) of the five indicated pairwise comparisons between architectures; cells are shown only for miRNAs passing |log2(sorafenib/vehicle)| > 0.5 and P < 0.05, and are blank otherwise. miRNAs discussed in the text are indicated below the heatmaps. **b** Volcano plots comparing the sorafenib response of the 1U reference design with each alternative design. x, difference in log2(sorafenib/vehicle) between the two architectures (Δlog2(sorafenib/vehicle)); y, −log10(adjusted P value), Benjamini–Hochberg. Dashed lines, |Δlog2(sorafenib/vehicle)| = 0.5 and P = 0.05; orange, miRNAs passing both cut-offs; grey, all others. Enlarged points denote the labeled miRNAs and carry no quantitative meaning. **c** Validation of the five miRNAs highlighted in Fig. 2h. MHCC97-L cells carrying a doxycycline-inducible 7merSupp (green) or 7mer (red) variant of each miRNA were induced for 5 days and treated with 3.25 µg/mL sorafenib for a further 5 days; viability (CCK-8) is expressed relative to the matched vehicle control. Connected point pairs are individual biological replicates (n = 3, except the sorafenib-treated 7merSupp condition for miR-494-3p, n = 2). *P < 0.05, **P < 0.01, Student’s t-test; unmarked comparisons are not significant. **d** Strategy used to identify the effector target of miR-494-3p 3′-supplementary pairing. RNA-seq of cells overexpressing each miR-494-3p variant, differential expression between 7merSupp and 7mer (838 genes), a targeted shRNA-library screen of those genes under sorafenib versus vehicle (4 genes), and a search for genes carrying both seed and 3′-supplementary complementarity (1 gene, RPLP0).

**Supplementary Fig. 11:**
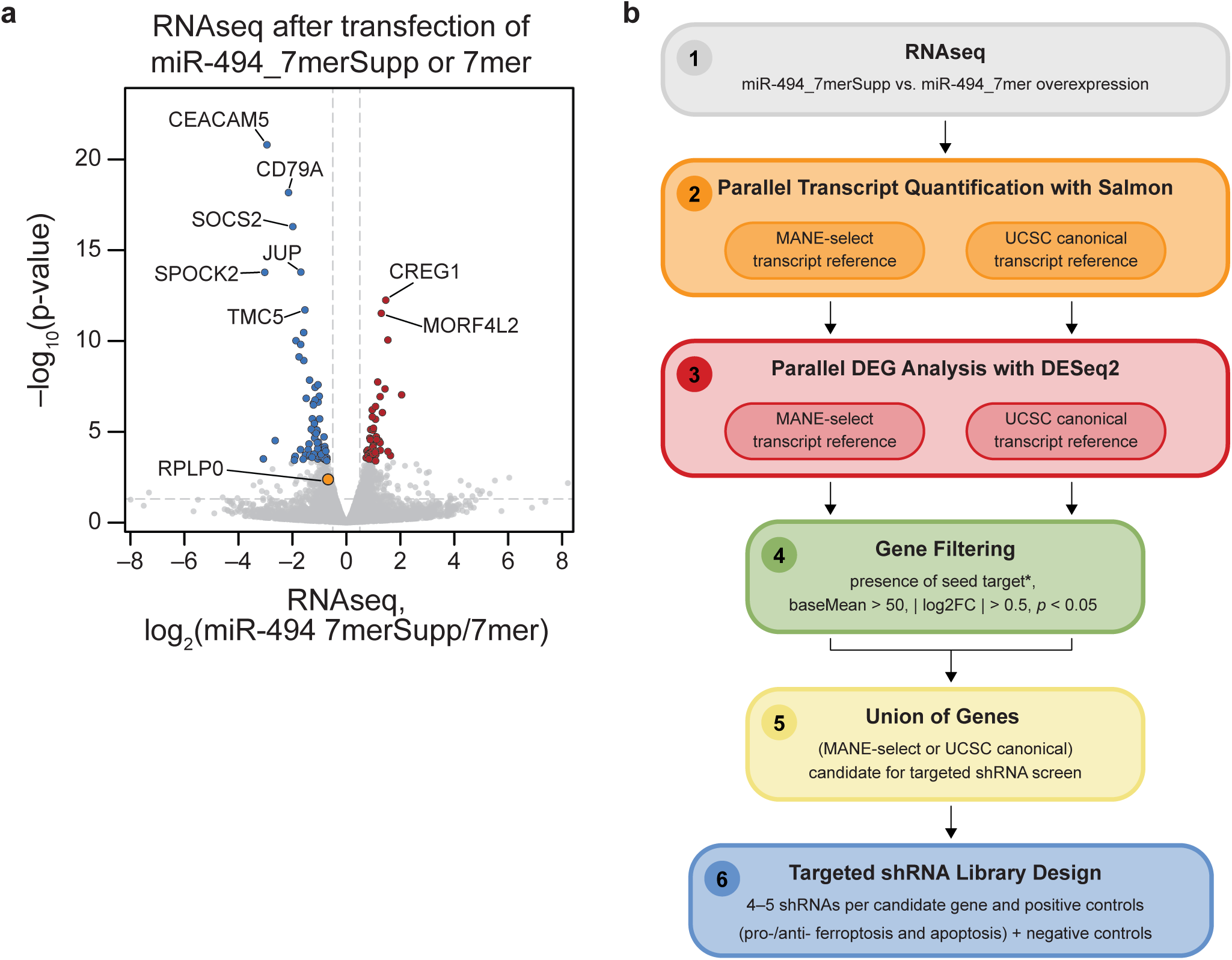
DEG analysis upon miR-494 variants overexpression and pipeline for targeted shRNA screen library design. **a** Volcano plot of differential expression between MHCC97-L cells overexpressing miR-494_7merSupp and miR-494_7mer. x axis, RNA-seq log2(7merSupp/7mer); y axis, −log10(P value). Blue, significantly upregulated genes; red, significantly downregulated genes; grey, non-significant genes. RPLP0 is highlighted in orange. Dashed lines indicate the fold-change and significance cut-offs. **b** Analysis and library-design workflow: (1) RNA-seq of miR-494_7merSupp-versus miR-494_7mer-expressing cells; (2) transcript quantification with Salmon against the MANE Select and UCSC canonical references in parallel; (3) differential expression with DESeq2 for each reference; (4) gene filtering (presence of a canonical seed match, baseMean > 50, |log2(7merSupp/7mer)| > 0.5, P value < 0.05); (5) union of genes passing in either reference; (6) design of a targeted shRNAone sublibrary comprising 4–5 shRNAs from TRC-like shRNAone library (see Supplementary Table 6) per candidate gene, shRNAs against pro- and anti-ferroptotic and apoptotic control genes, and non-targeting controls.

**Supplementary Fig. 12:**
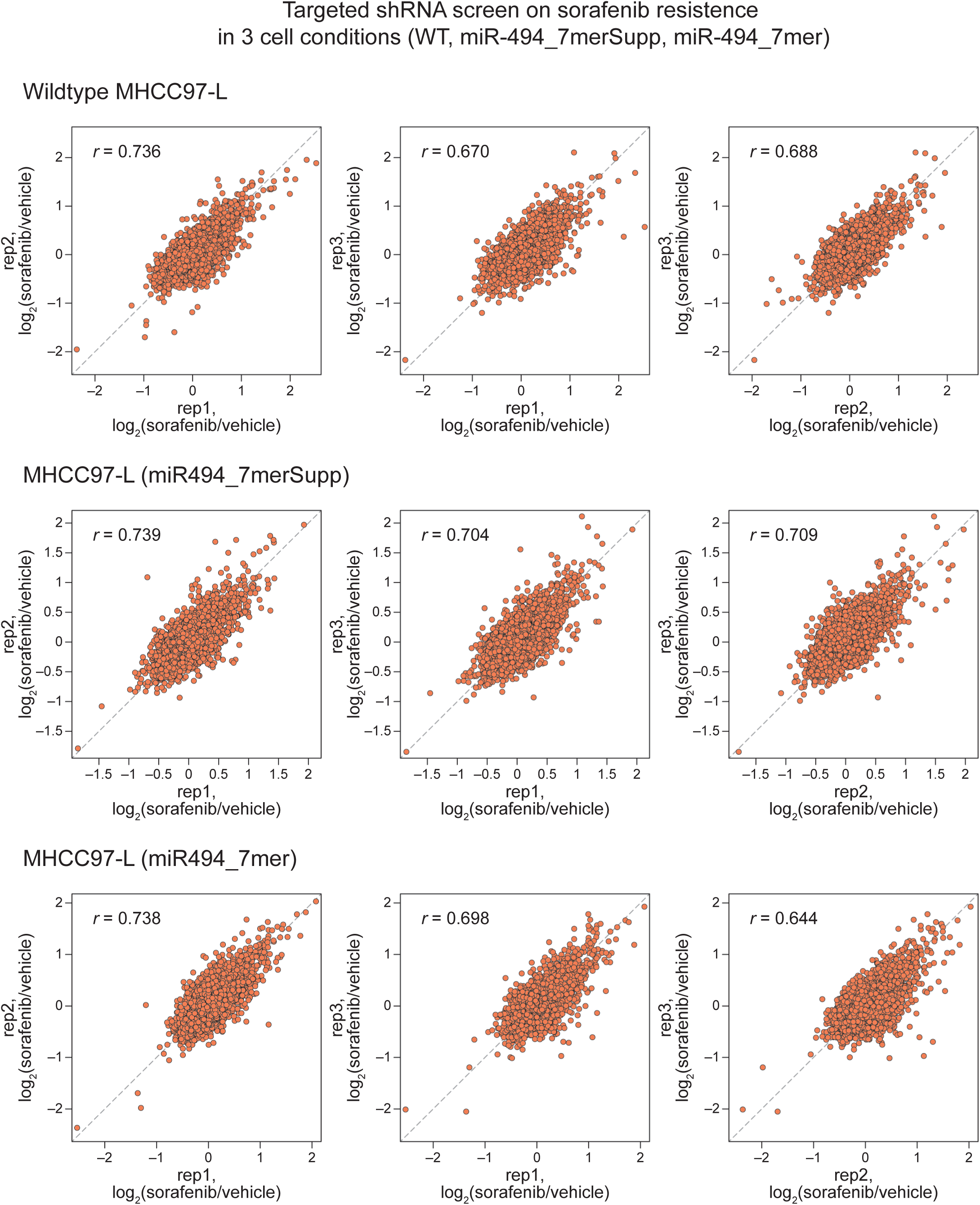
Reproducibility of the targeted shRNA screens. Pairwise comparison of shRNA-level log2(sorafenib/vehicle) enrichment between the three biologically independent replicates of each screen: wild-type MHCC97-L cells (upper; Pearson’s r = 0.736, 0.670 and 0.688), MHCC97-L miR-494_7merSupp cells (middle; r = 0.739, 0.704, and 0.709) and MHCC97-L miR-494_7mer cells (bottom; r = 0.738, 0.698, and 0.644).

**Supplementary Fig. 13:**
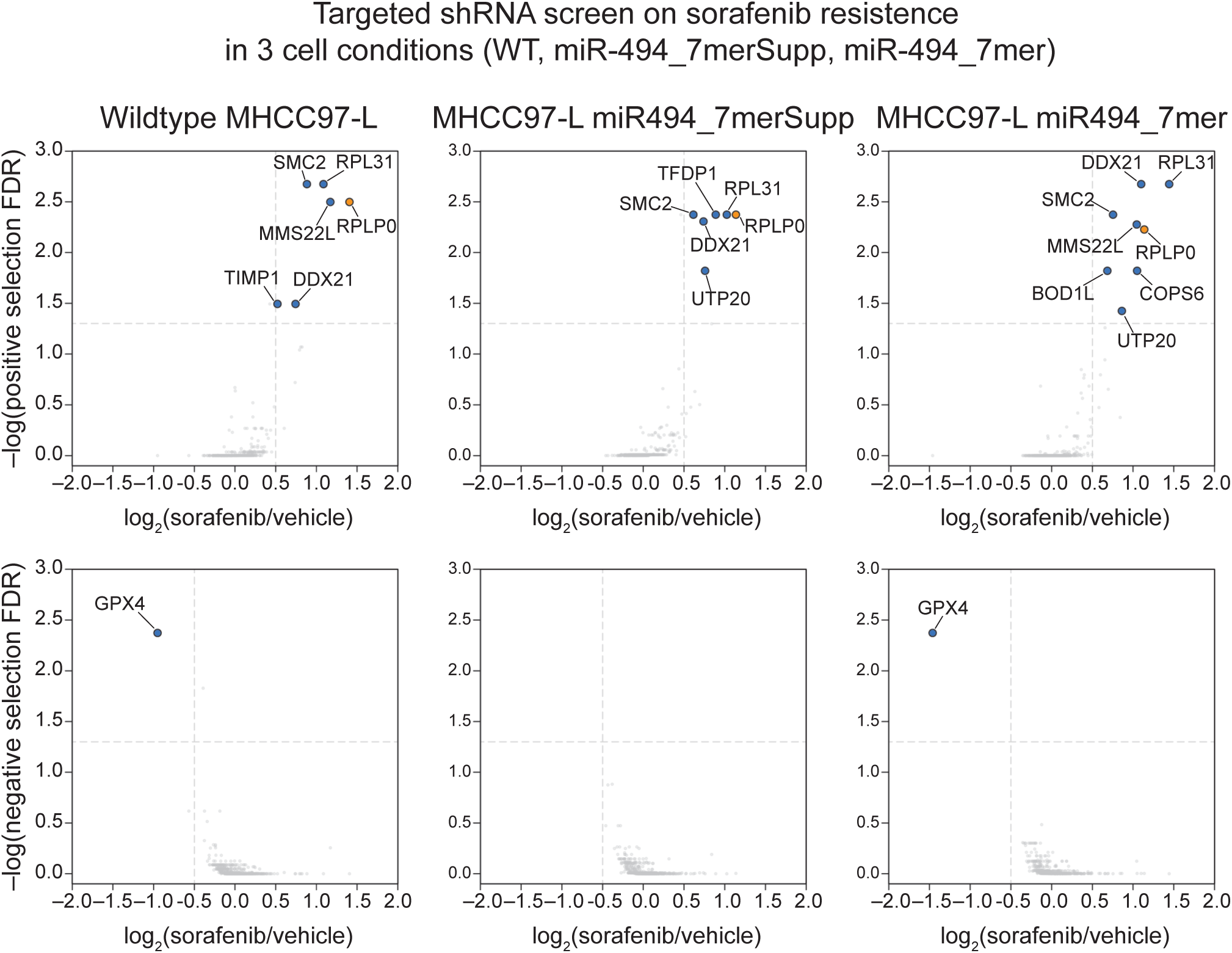
Positive- and negative-selection statistics from MAGeCK analysis for the targeted shRNA sorafenib screens. Gene-level results of the targeted shRNA screen in wild-type MHCC97-L cells (left), MHCC97-L miR-494_7merSupp cells (middle) and MHCC97-L miR-494_7mer cells (right). GPX4 is a positive control gene for the ferroptosis pathway. x axis, log2(sorafenib/vehicle); y axis, −log10(MAGeCK positive-FDR) (upper) or −log10(MAGeCK positive-FDR) (lower). Dashed lines mark −log10(FDR) = 0.05 (horizontal) andlog2FC = +0.5 (upper) or −0.5 (lower) (vertical). Blue, genes pass both thresholds; orange, RPLP0; grey, non-significant genes.

**Supplementary Fig. 14:**
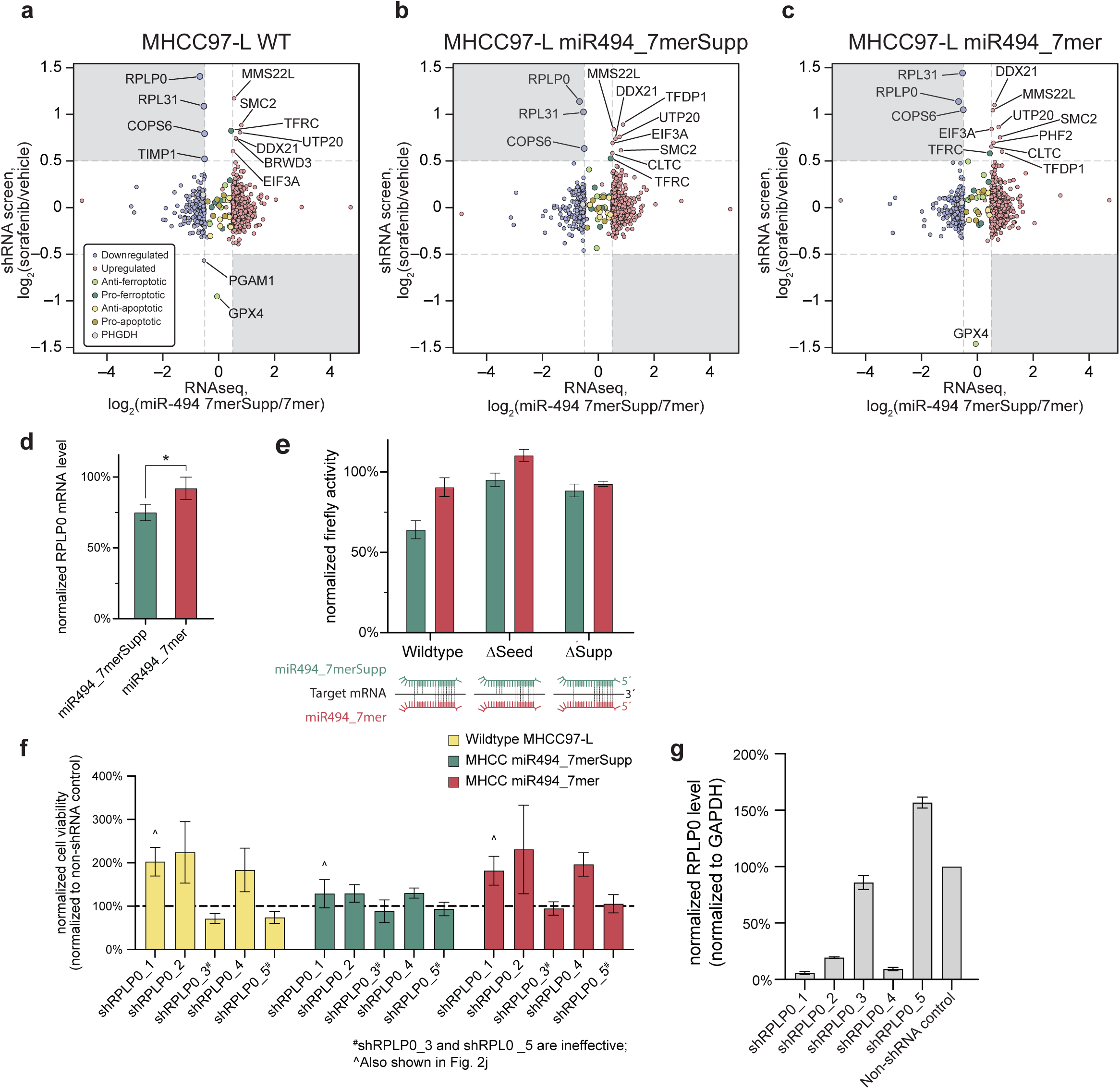
Integration of transcriptome and targeted shRNA screen data identifies RPLP0 as a supplementary-dependent target. **a–c** Integration of the RNAseq and targeted shRNA screen in MHCC97-L WT cells (a), miR494_7merSupp-expressing cells (b), and miR-494_7mer-expressing cells (c). Targeted shRNA-screen log2(sorafenib/vehicle) (y) versus RNA-seq log2(miR-494_7merSupp/7mer) (x) for every gene in the targeted library. Dashed lines, ±0.5 on both axes; shaded quadrants mark potential target genes. Point color indicates RNA-seq direction of change or curated ferroptosis/apoptosis controls, as shown; enlarged points denote labeled genes. **d** qRT-PCR of RPLP0 mRNA in MHCC97-L cells after 5 days of doxycycline-induced miR494_7merSupp (green) or miR494_7mer (red) expression, normalized to GAPDH and to the uninduced control. Bars, mean ± s.d.; n = 3 biological replicates. *P < 0.05, Student’s t-test. **e** Luciferase reporter assay in MHCC97-L cells. Firefly activity, normalized to Renilla and to the non-shRNA control, for reporters carrying the wild-type RPLP0 target region or the same region with the seed match (ΔSeed) or the supplementary match (ΔSupp) mutated, co-expressed with miR494_7merSupp (green) or miR494_7mer (red). Schematics show the pairing retained by each combination. Bars, mean ± s.d.; n = 3 biological replicates. **f** Normalized cell viability upon RPLP0 knockdown and non-shRNA control (yellow) or miR-494_7merSupp (green) or miR-494_7mer (red) expression. Cell viability is normalized to the corresponding non-shRNA control (dashed line, 100%). #shRPLP0_3 and shRPLP0_5 do not deplete RPLP0. Bars, mean ± s.d.; n = 3 biological replicates. Samples marked with ^ are also shown in Fig. 2j. **g** RPLP0 mRNA measured by qRT-PCR in wild-type MHCC97-L cells expressing each of five RPLP0-targeting shRNAs, normalized to GAPDH and to the non-shRNA control (n = 3). Data are mean ± s.d.

**Supplementary Fig. 15:**
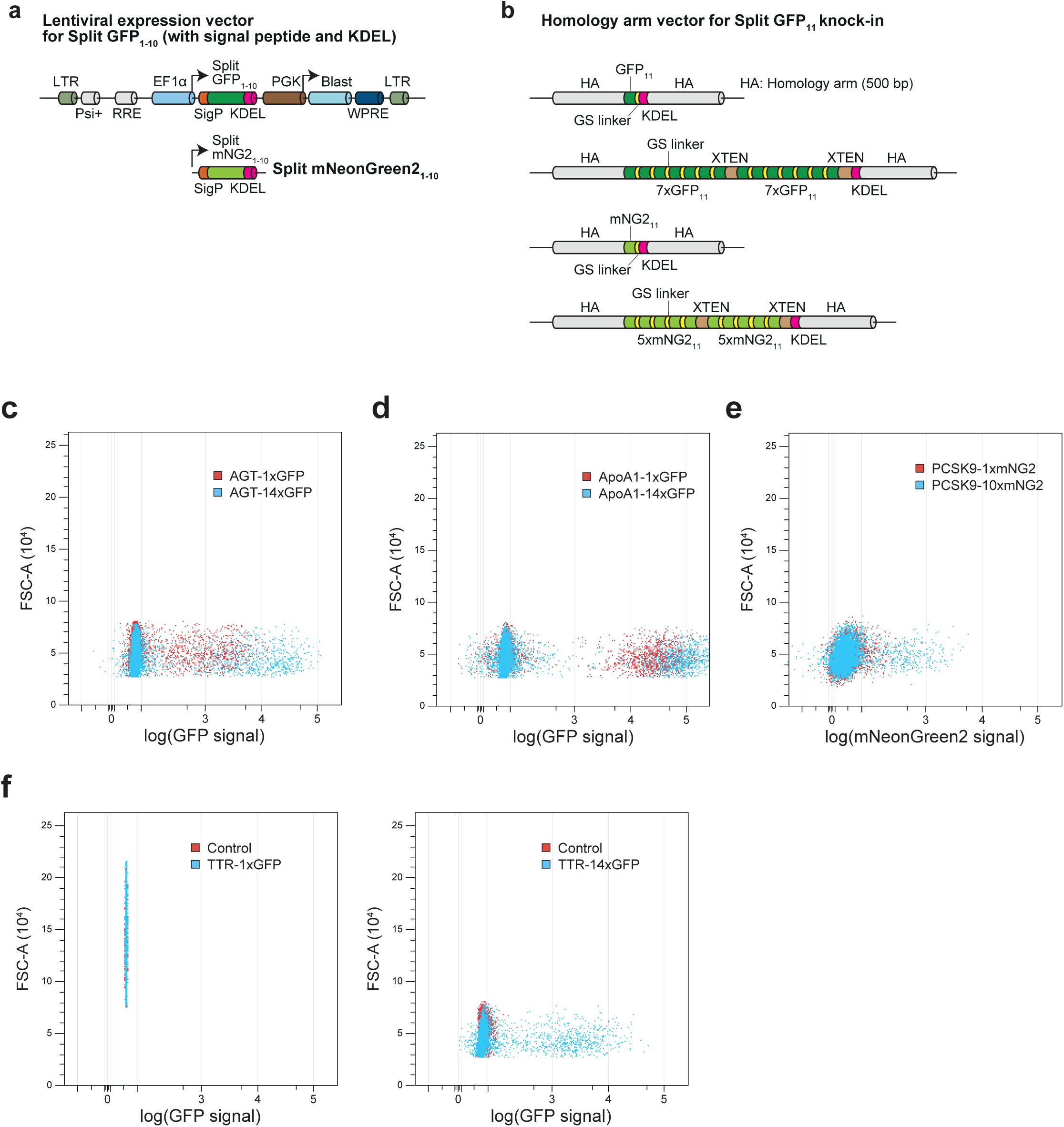
Split-fluorescent protein knock-in reporter cells for endogenous secretory proteins. **a** Lentiviral vectors expressing ER-targeted split GFP1–10 or split mNeonGreen21–10 (mNG21–10). **b** Donor vectors for knock-in of the split GFP11 or split mNG211 tag at the 3′ end of the endogenous coding sequence. **c**–**f** Flow cytometry of polyclonal knock-in populations in MIHA cells before FACS enrichment, plotting forward scatter (FSC-A) against log fluorescence intensity: AGT-1×GFP11 (red) versus AGT-14×GFP11 (blue) (c); ApoA1-1×GFP11 versus ApoA1-14×GFP11 (d); PCSK9-1×mNG211 versus PCSK9-10×mNG211 (e); and untransfected control versus TTR-1×GFP11 (left) and versus TTR-14×GFP11 (right) (f).

**Supplementary Fig. 16:**
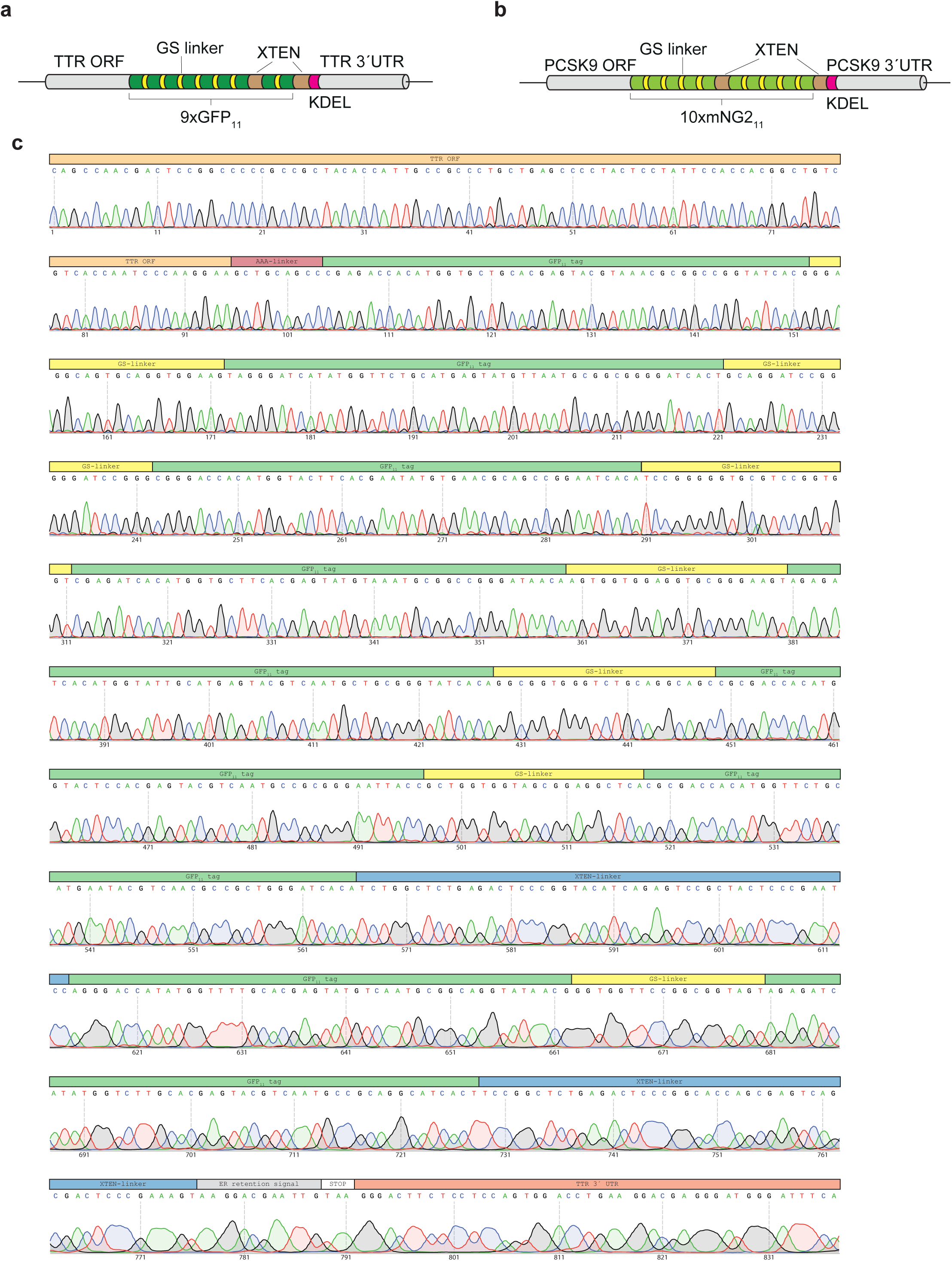
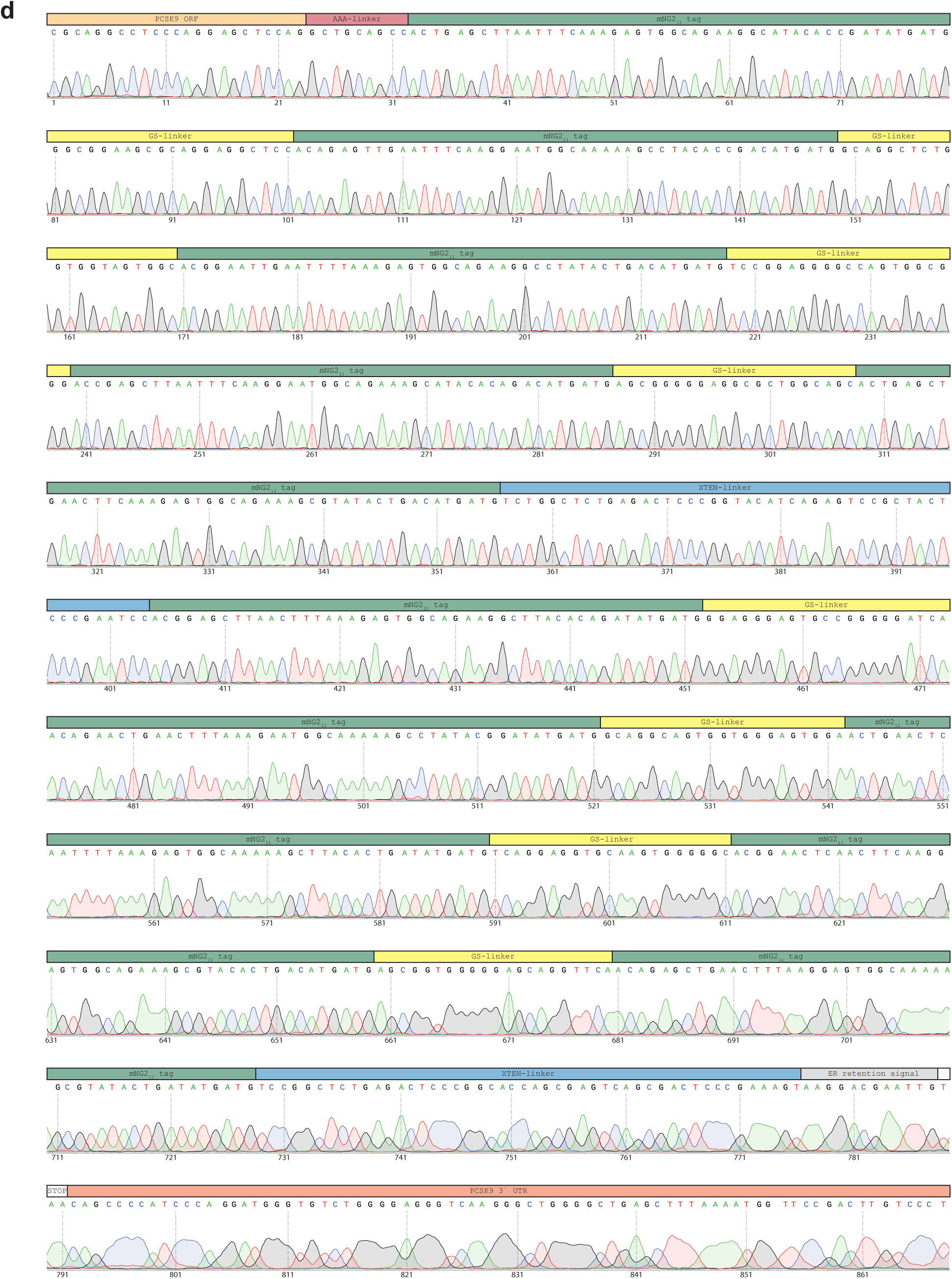
Structure and sequence validation of the TTR and PCSK9 reporter single-cell clone. **a**, **b** Schematics of the validated knock-in at TTR (a) and PCSK9 loci (b). **c**, **d** Sanger sequencing chromatograms spanning the knock-in cassette of the single-cell-derived TTR-9×GFP11 reporter line (c) and the PCSK9-10×mNG211 reporter line (d).

**Supplementary Fig. 17:**
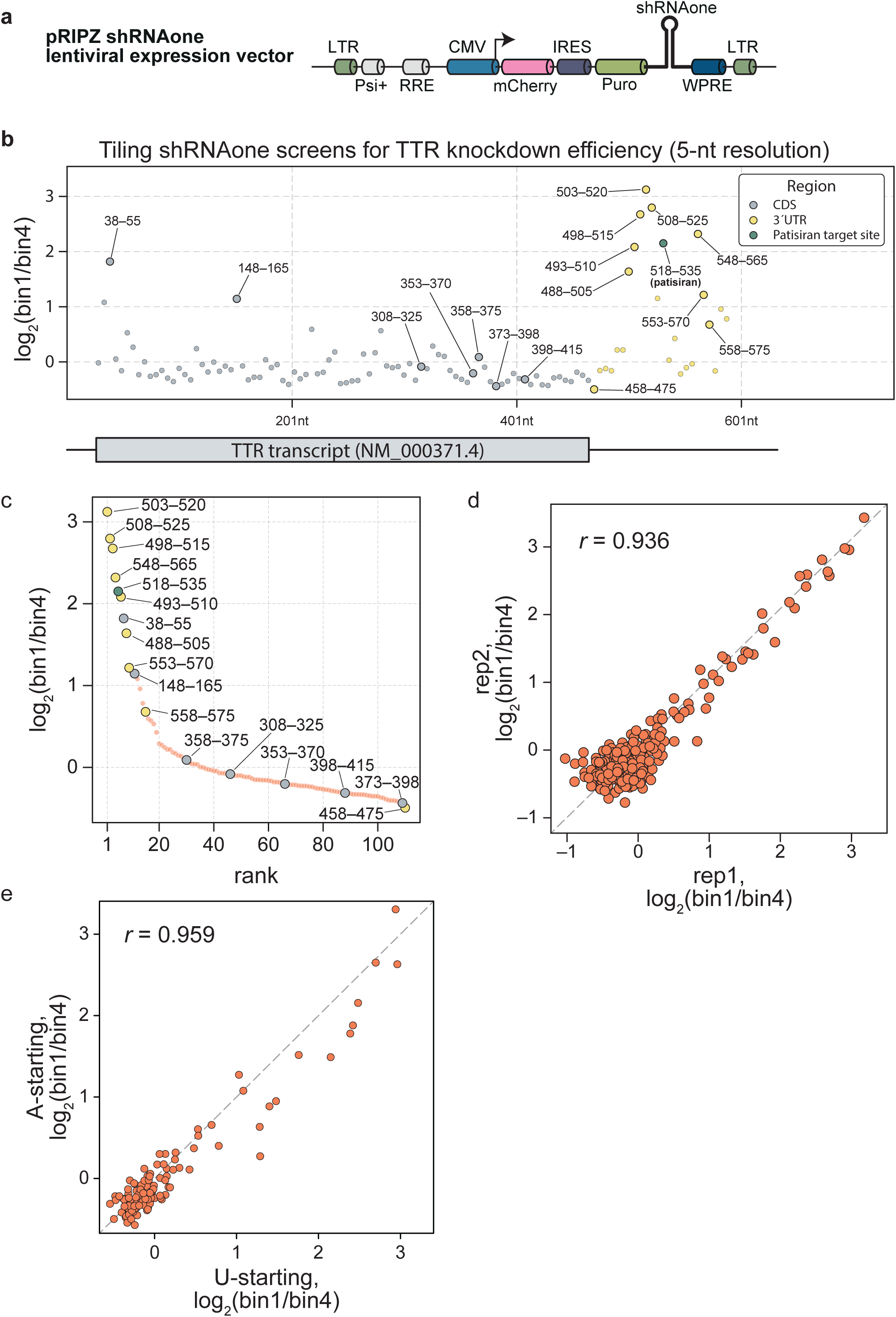
Pilot shRNAone 5-nt tiling screen of the TTR transcript. **a** Map of the pRIPZ shRNAone lentiviral expression vector. LTR, long terminal repeat; Psi+, packaging signal; RRE, Rev response element; WPRE, woodchuck hepatitis virus post-transcriptional regulatory element. **b** Knockdown score, expressed as log2(bin1/bin4), for each shRNA plotted against the position of its target site on TTR (NM_000371.4). Grey, CDS target sites; yellow, 3′ UTR target sites; teal, patisiran target site. **c** The same data ranked from most to least enriched; labeled points are colored as in b. **d** Correlation of log2(bin1/bin4) between two biological replicates (Pearson’s r = 0.936). **e** Correlation of log2(bin1/bin4) between the U- and A-starting sublibraries (Pearson’s r = 0.959). Dashed lines in d and e indicate y = x.

**Supplementary Fig. 18:**
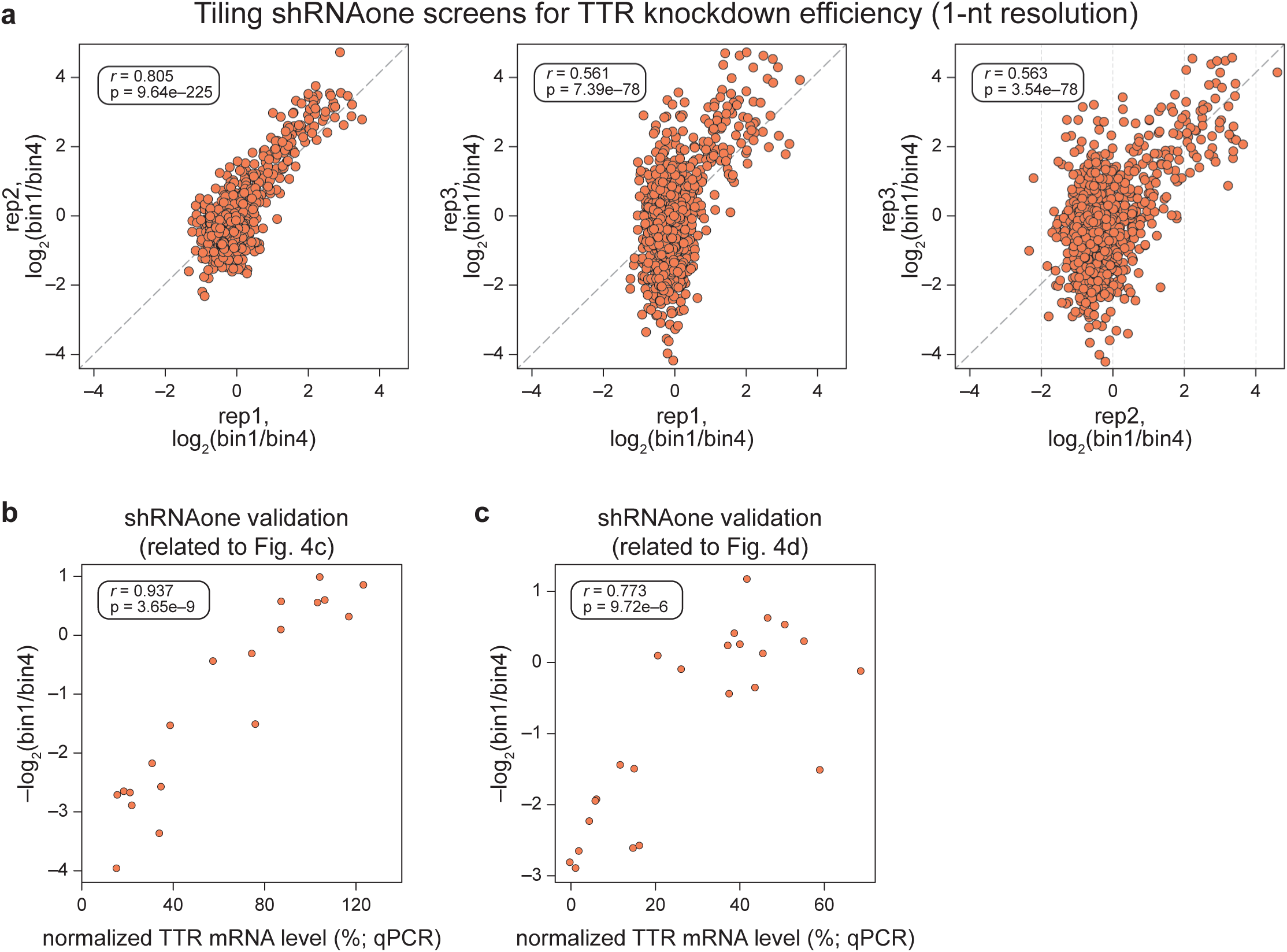
Single-nucleotide-resolution shRNAone tiling screen of TTR. **a** Pairwise comparison of log2(bin1/bin4) between the three biological replicates of the 1-nt TTR tiling screen (n = 1,090). Pearson’s r = 0.805, P value = 9.64 × 10^−225^ (rep1 vs rep2); r = 0.561, P = 7.39 × 10^−78^ (rep1 vs rep3); r = 0.563, P = 3.54 × 10^−78^ (rep2 vs rep3). Dashed lines indicate y = x. **b**, **c** Correlation of individual validation to log2(bin1/bin4) (related to Fig. 4c and Fig. 4d, respectively). Normalized endogenous TTR mRNA measured by qRT-PCR is plotted against the pooled-screen score, −log2(bin1/bin4). Pearson’s r = 0.937, P value = 3.65 × 10^−9^ (b) and r = 0.773, P value = 9.72 × 10^−6^ (c).

**Supplementary Fig. 19:**
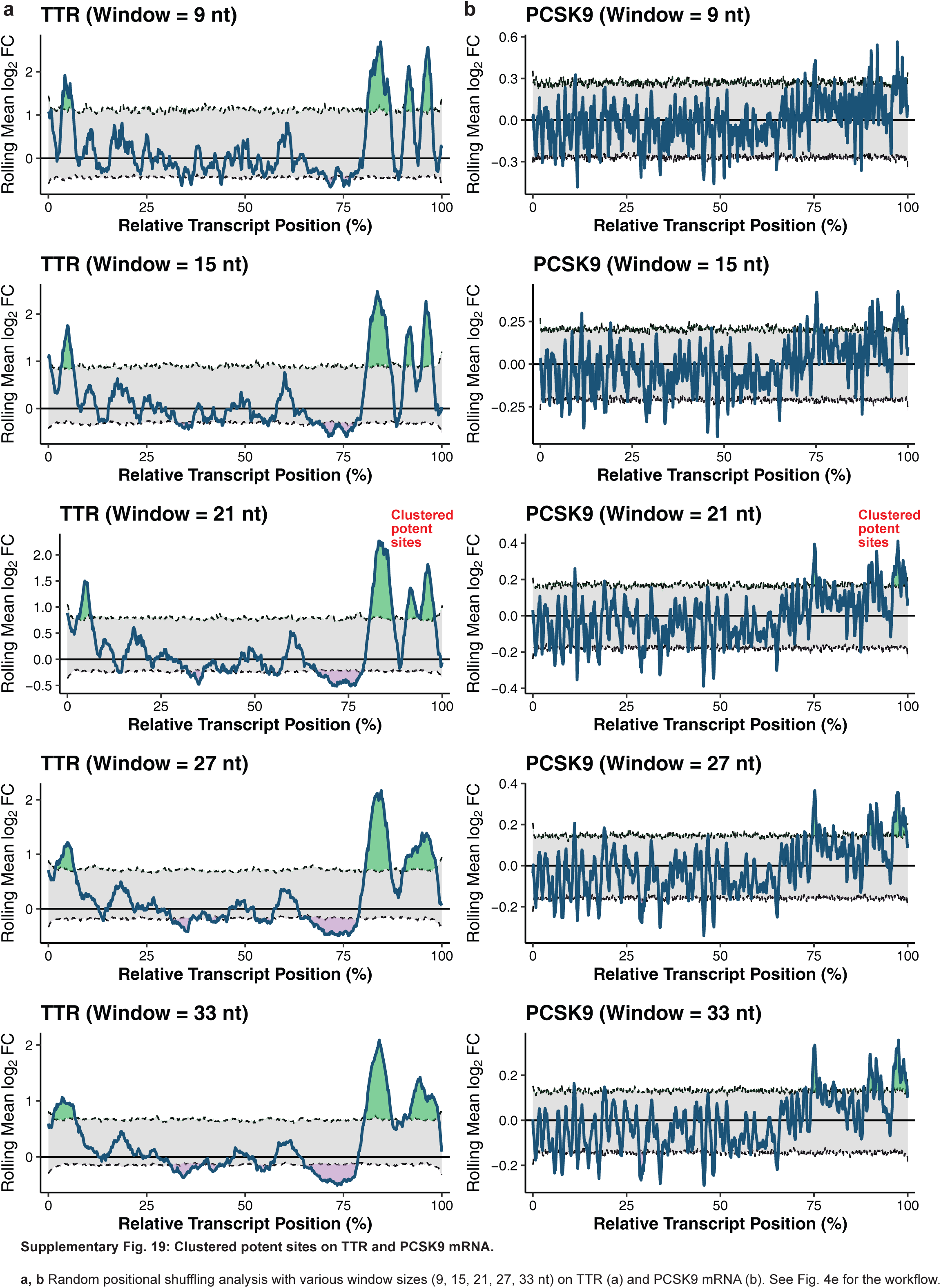
Clustered potent sites on TTR and PCSK9 mRNA. **a, b** Random positional shuffling analysis with various window sizes (9, 15, 21, 27, 33 nt) on TTR (a) and PCSK9 mRNA (b). See Fig. 4e for the workflow.

**Supplementary Fig. 20:**
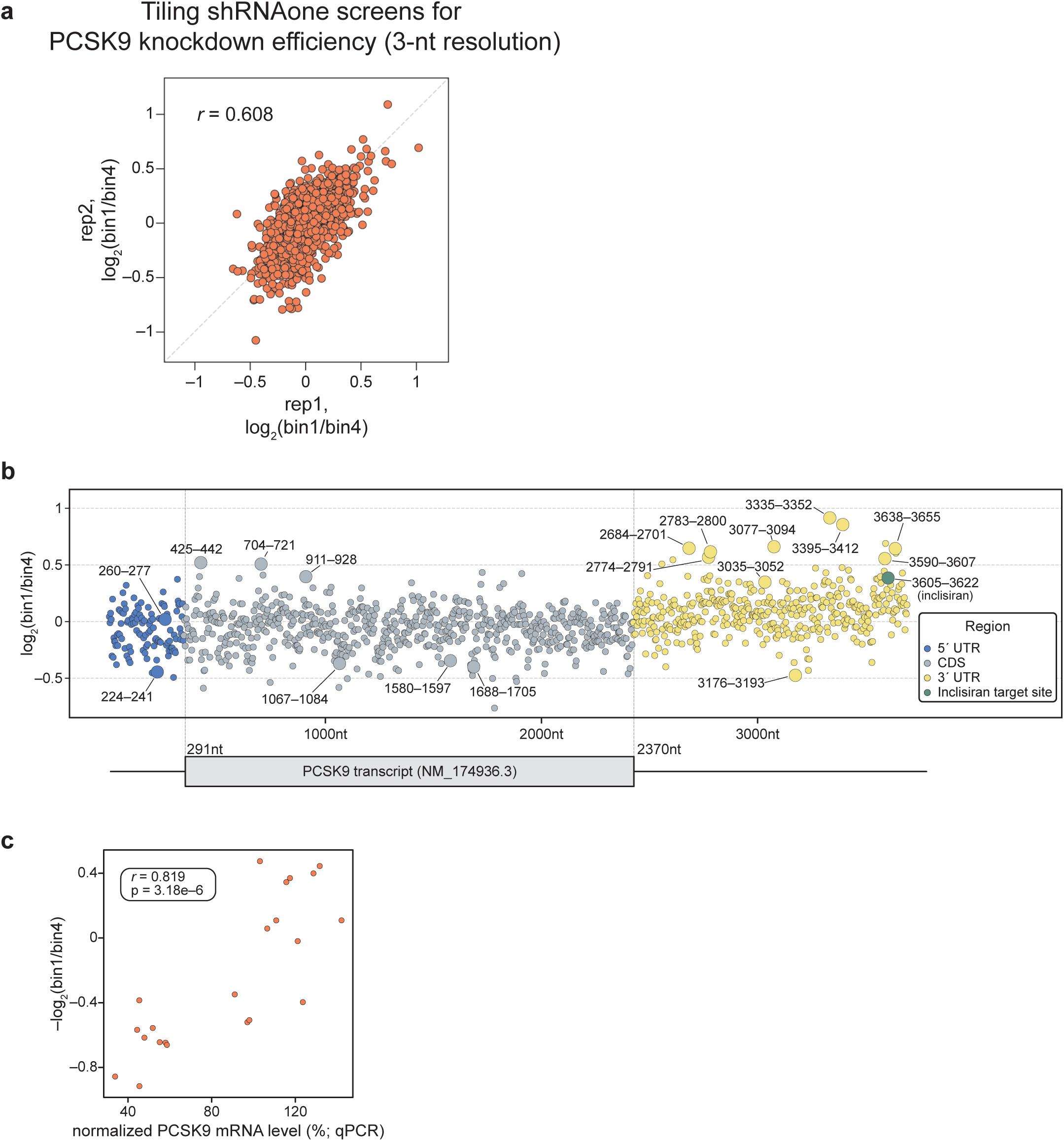
shRNAone tiling screen of the PCSK9 transcript at 3-nt resolution. **a** Correlation of log2(bin1/bin4) between two biological replicates of the PCSK9 3-nt tiling screen (n = 1,230). Pearson’s r = 0.608; dashed line indicates y = x. **b** shRNA log2(bin1/bin4), from a tiling shRNAone screen in clonal PCSK9–10×mNG211 MIHA reporter cells (screen design as in Fig. 3a), plotted against target-site position and colored by transcript region; the inclisiran site (3605–3622 nt) is highlighted. Enlarged points, shRNAs validated in Fig. 5c. Transcript model below (box, coding sequence 291–2370 nt; lines, UTRs). **c** –log2(bin1/bin4) versus arrayed qPCR knockdown for the 22 shRNAs in Fig. 5c. Two-sided Pearson correlation.

**Supplementary Fig. 21:**
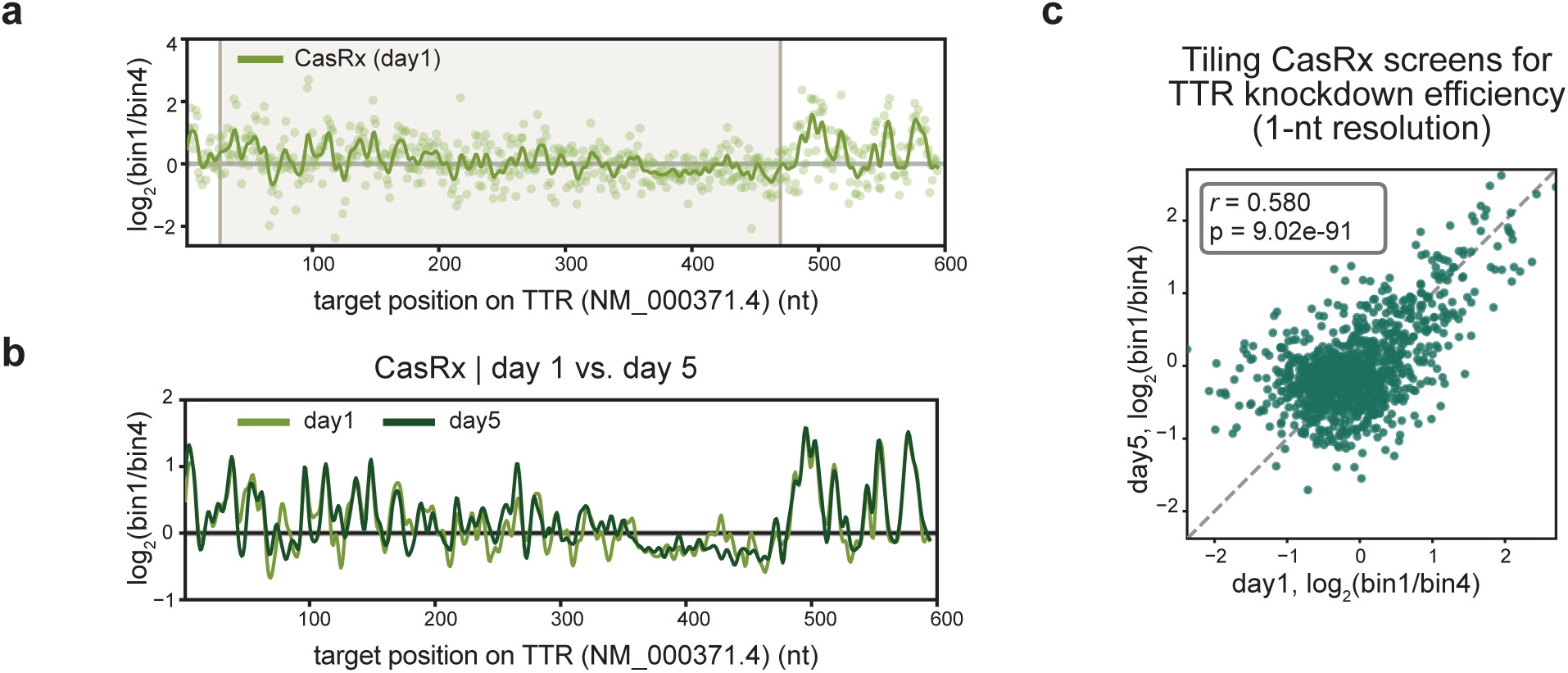
Day-1-sorted CasRx crRNA 1-nt tiling screen library and correlation to day-5-sorted library. **a** Gaussian smoothed line for log2(bin1/bin4) across TTR transcript (NM_000371.4) from day-1-sorted MIHA reporter cells infected by CasRx crRNA lentivirus library. Individual crRNAs are shown as points; the shaded band marks the coding sequence (27–470 nt). **b** Overlapping knockdown Gaussian smoothed line for CasRx screen sorted on day 1 (light green) and 5 (dark green). **c** Correlation of crRNA enrichments log2(bin1/bin4) between day 1 and 5. Pearson’s r = 0.580, P value = 9.02 × 10^−91^.

**Supplementary Fig. 22:**
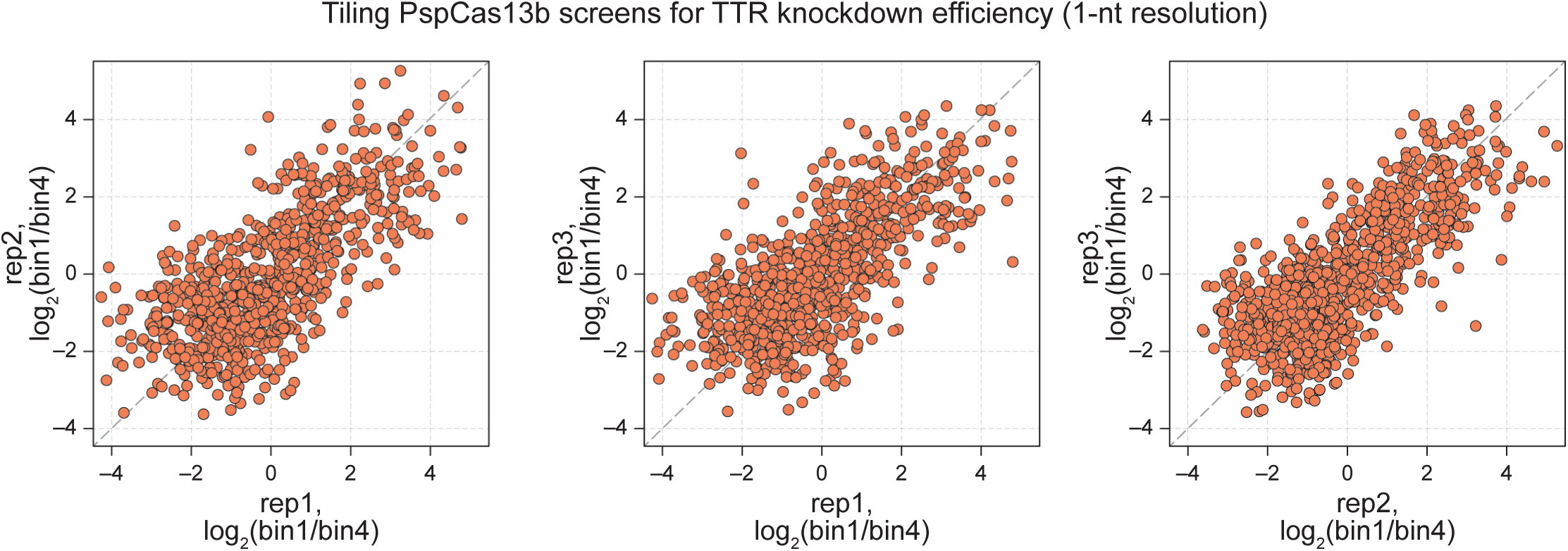
Reproducibility of the single-nucleotide-resolution PspCas13b tiling screen of the TTR transcript. Pairwise comparison of crRNA-level log2(bin1/bin4) between the three biological replicates of the PspCas13b tiling screen: rep1 versus rep2 (left), rep1 versus rep3 (middle) and rep2 versus rep3 (right). Pearson’s r = 0.707 (left), 0.700 (middle), 0.763 (right), P value = 2.50 × 10^−117^, 2.39 × 10^−114^, 2.47 × 10^−173^. Dashed lines indicate y = x.

**Supplementary Fig. 23:**
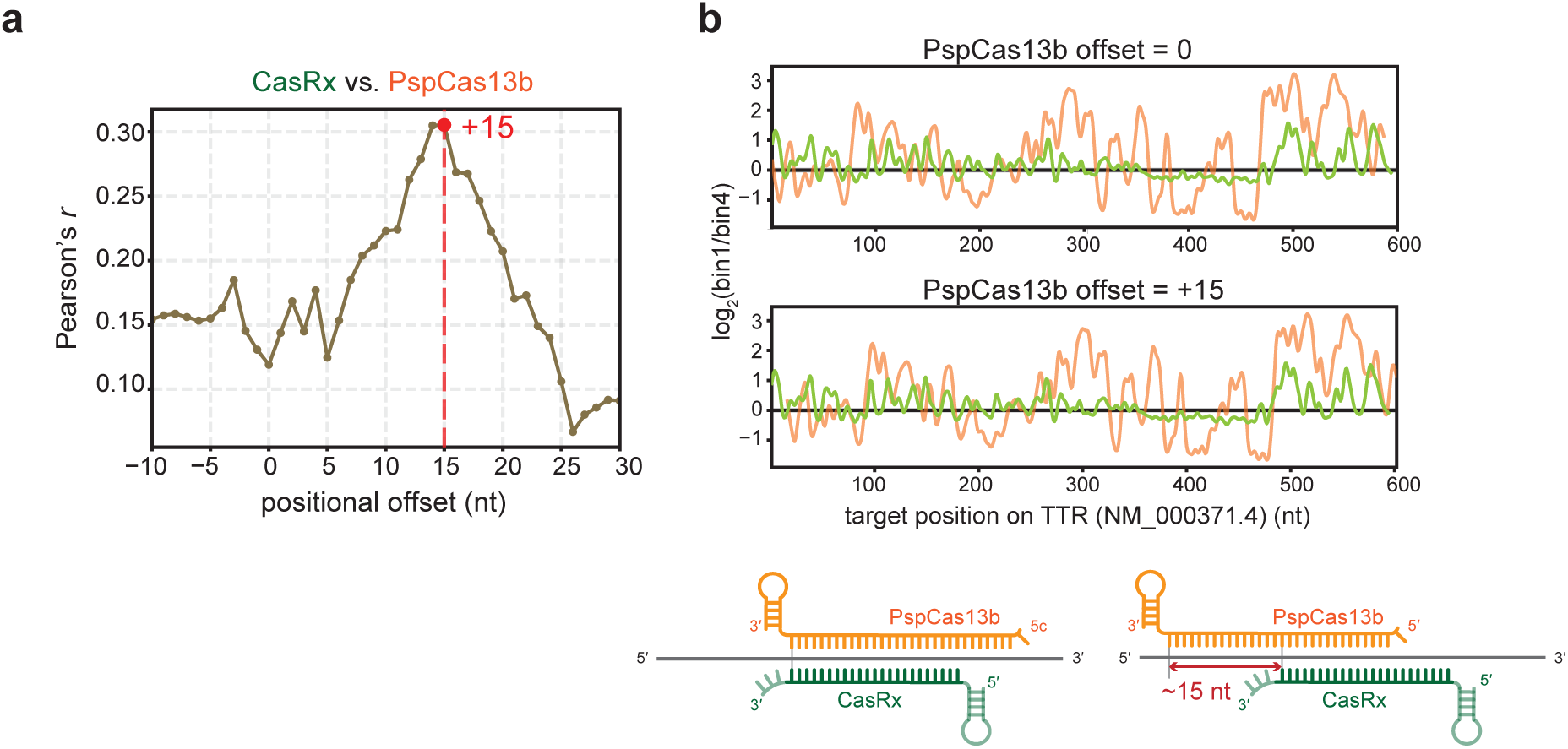
Positional offset alignment on TTR knockdown landscape between CasRx and PspCas13b crRNA 1-nt tiling screens. **a** Optimal positional offset analysis comparing cytolytic CasRx to PspCas13b. The PspCas13b profile was shifted by 15 nt towards the 3′ end to align log2FC data points with CasRx; correlations were calculated using Pearson’s r. **b** Shifted knockdown profiles with schematics for CasRx (green) and PspCas13b (orange).

**Supplementary Fig. 24:**
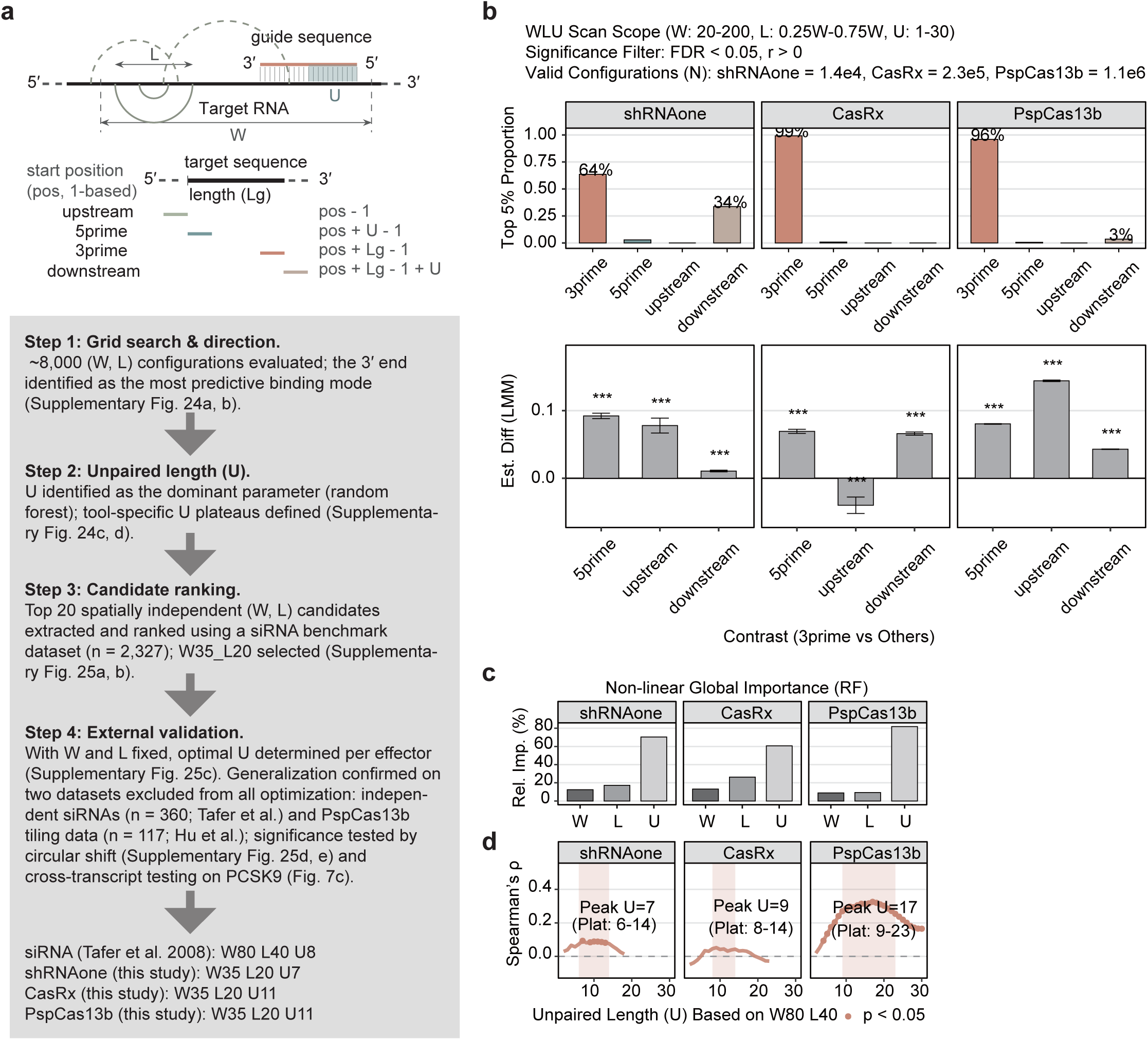
Systematic identification of optimal accessibility parameters (Steps 1 and 2). **a** Schematic illustration of the RNAplfold accessibility computation. Upper panel: a target RNA of fixed length is characterized by two parameters — the sliding window size (W), within which the maximum base-pairing distance (L) constrains the fold, and the unpaired region length (U). Guide sequence (red) hybridizes to the target via base pairing; the U segment at the 5′ end of the guide denotes the unpaired region whose accessibility is quantified. Allowed (solid arc) and disallowed (dashed arc) base pairs illustrate how W and L jointly define the folding landscape. Lower panel: formal definitions of the four direction schemes used to extract accessibility from the RNAplfold output matrix, expressed as functions of the target start position (pos), the guide length (Lg), and the unpaired constraint (U). **b** Direction assessment restricted to the Whole transcript region. Upper panel: proportion of top-5% (by Spearman r) parameter configurations attributed to each of the four direction definitions (3prime, 5prime, upstream, downstream), shown as individual bars faceted by tool (shRNAone, CasRx, PspCas13b). All four directions are displayed regardless of representation; bars with zero proportion are retained to enable direct comparison. Lower panel: linear mixed-effects model (LMM) estimates of the fixed-effect difference between 3prime and each alternative direction within the Whole region, with error bars denoting 95% confidence intervals. Positive estimates indicate superior performance of the 3prime direction. Asterisks denote significance of the contrast (***P < 0.001). The global title reports the WLU scan scope and the number of valid FDR-significant (FDR < 0.05, r > 0) configurations retained for each tool across all regions. **c** Non-linear parameter importance assessed by random forest permutation importance. Bar height represents the relative increase in mean squared error (%IncMSE) when each thermodynamic parameter (W, L, U) is randomly permuted. Results are shown separately for shRNAone, CasRx, and PspCas13b, computed from all grid-search configurations with positive Spearman r under the 3prime direction. **d** Reference U plateau trajectories under the classical RNAxs parameters (W = 80, L = 40). Spearman correlation between predicted 3prime accessibility and experimental log2FC is plotted as a function of the unpaired region length (U) for the Whole transcript region. The shaded area marks the plateau interval, defined as the U range where Spearman r exceeds 75% of its peak value. Solid circles denote U values achieving raw P < 0.05; open circles are non-significant. Peak U and plateau boundaries are annotated for each tool.

**Supplementary Fig. 25:**
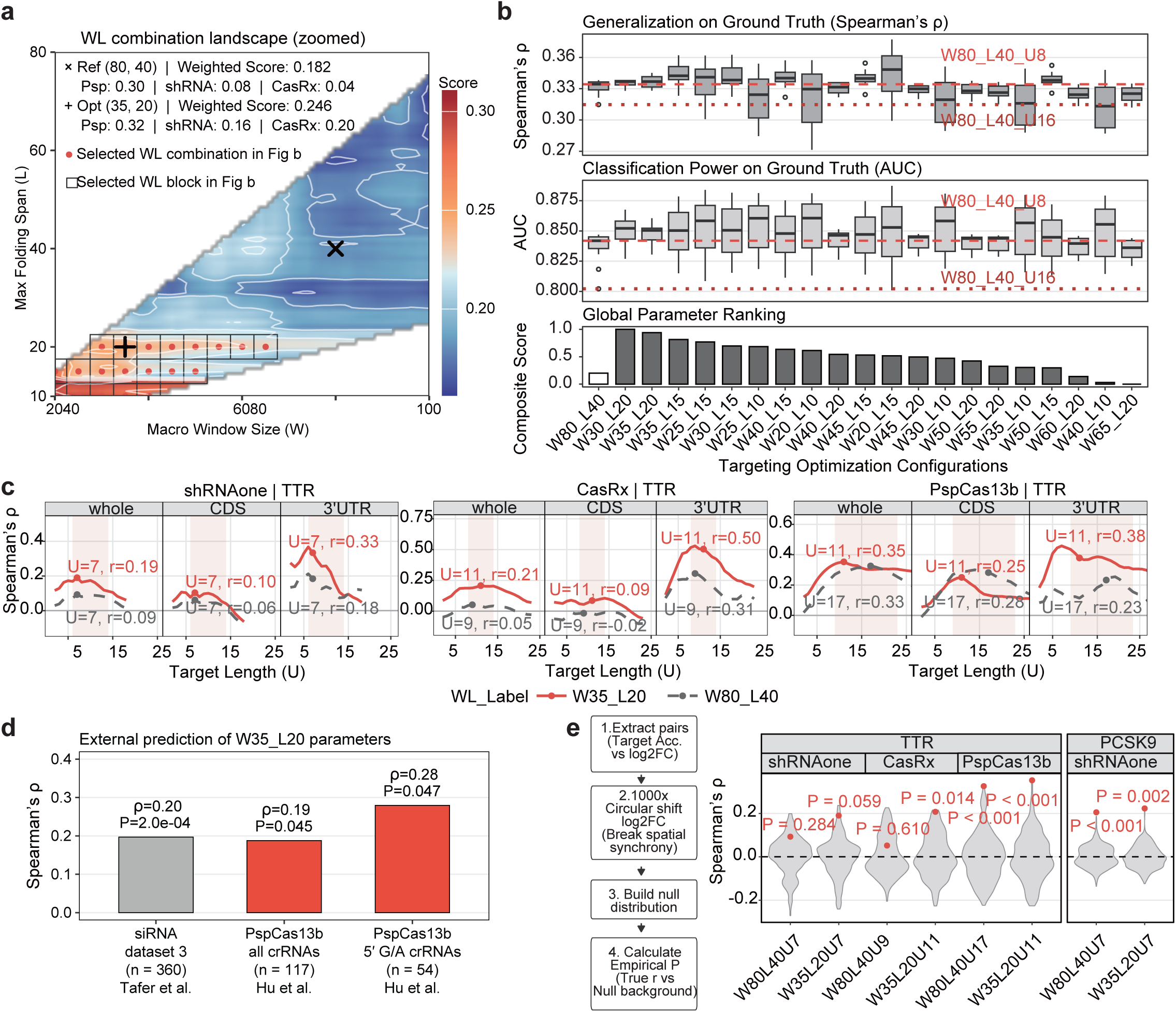
Systematic identification of optimal accessibility parameters (Steps 3 and 4). **a** Weighted mean score landscape of (W, L) parameter combinations within the U plateau defined in panel d. The color gradient represents the weighted mean Spearman r across the three effectors (PspCas13b, 0.50; shRNAone, 0.30; CasRx, 0.20), computed from the whole region and constrained to U values within each tool’s plateau. White contour lines delineate iso-performance regions. The reference point (W = 80, L = 40; ×) and the spatial-NMS-selected optimum (W = 35, L = 20; +) are marked. Red points and thin grey rectangles indicate the top 20 spatially independent 5 × 5 block candidates selected by non-maximum suppression. The insets report the per-tool performance of the reference and optimal configurations. **b** Performance evaluation of candidate (W, L) combinations on the siRNA ground-truth dataset. top, Spearman correlation box plots across U values 6–16 for each candidate, ranked by their composite score (derived from a robust Z-score average of the weighted block score, the first quartile of siRNA Spearman r, and the first quartile of siRNA AUC). Red dashed and dotted lines mark the performance of the reference W80_L40 at U = 8 and U = 16, respectively. middle, Binary AUC box plots for the same candidates and reference benchmarks. bottom, Composite score of each candidate; the W80_L40 reference is indicated by an unfilled bar. **c** Comparison of U-scan trajectories between the TTR-derived optimal parameters (W = 35, L = 20; solid red) and the RNAxs reference parameters (W = 80, L = 40; dashed grey). Spearman correlation between 3prime accessibility and experimental log2FC is plotted as a function of U across three regions (Whole, CDS, 3′ UTR) and three effectors (CasRx, PspCas13b, shRNAone). The shaded rectangles mark the tool-specific U plateau intervals derived from the reference W80_L40 configuration. Peak U values and their corresponding r are annotated for each curve. **d** External validation of the optimized accessibility models. Spearman’s r between RNAplfold-predicted target-site accessibility and measured knockdown, computed on three datasets excluded from all stages of parameter optimization. Left: Tafer et al. siRNAs evaluated with the shRNAone model (W35_L20_U7; n = 360). Middle: all PspCas13b guides from Hu et al. evaluated with the PspCas13b model (W35_L20_U11; n = 117). Right: purine-starting subset of the same PspCas13b guides (n = 54). **e** Circular shift analysis assessing whether the observed correlations between accessibility and log2FC for TTR (left) and PCSK9 (right) exceed those expected under randomised spatial structure. Grey violin plots show the null distribution of Spearman r obtained from 1,000 circular shifts in which the log2FC vector was randomly shifted while preserving its autocorrelation structure. Red points mark the true observed r value; empirical two-sided P values are annotated. The left panel provides a schematic of the permutation workflow: (1) extract true accessibility–log2FC pairs; (2) circularly shift the log2FC vector to break point-to-point correspondence while preserving spatial autocorrelation; (3) build a null distribution from 1,000 such shifts; (4) compute the empirical P value as the fraction of permuted |r| exceeding the observed |r|.

**Supplementary Fig. 26:**
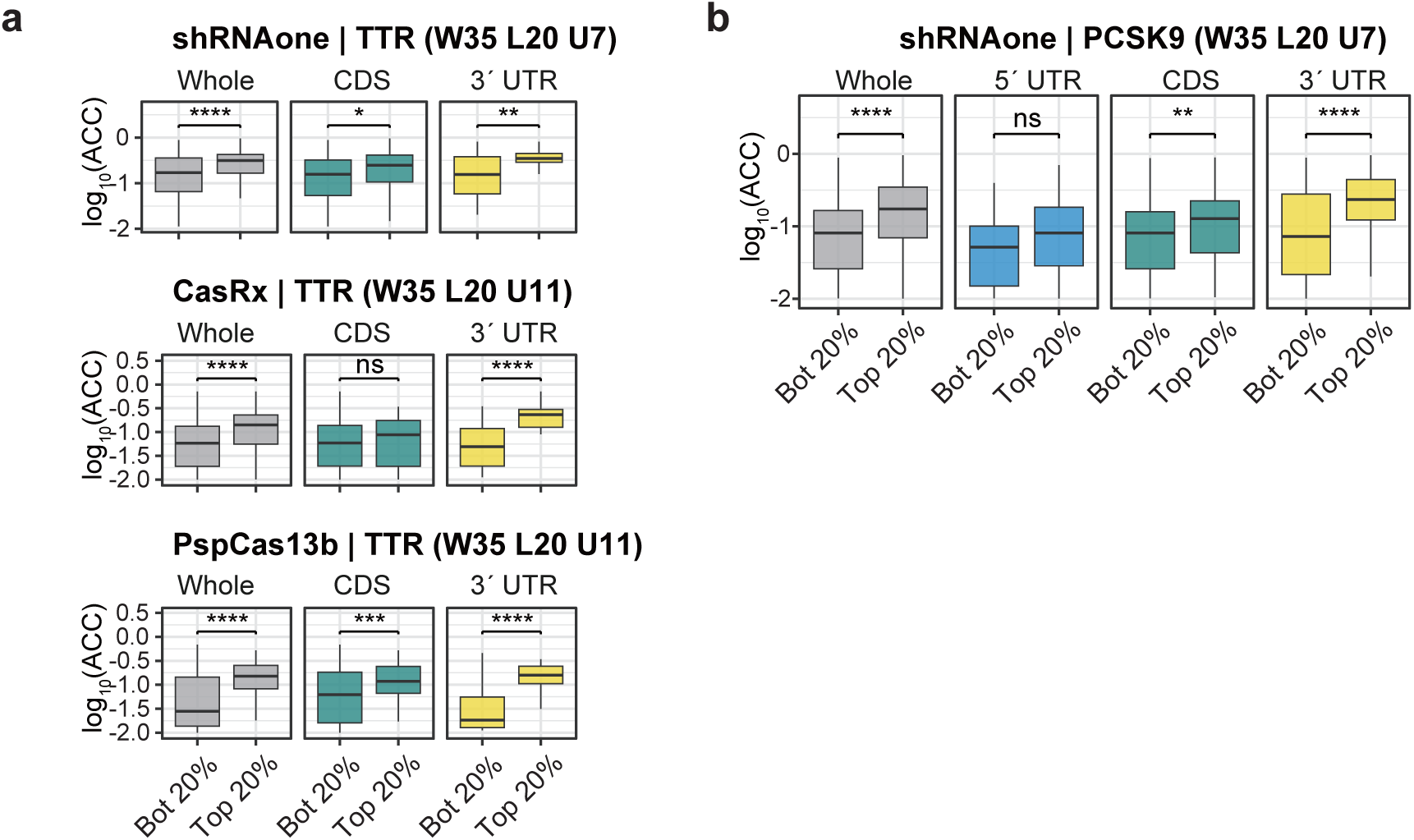
Target RNA accessibility is associated with guide efficacy across diverse targeting systems. **a, b** Box plots compare accessibility (W35 L20) between the top and bottom 20% of target sites ranked by log2FC. TTR (a), PCSK9 (b).

**Supplementary Fig. 27:**
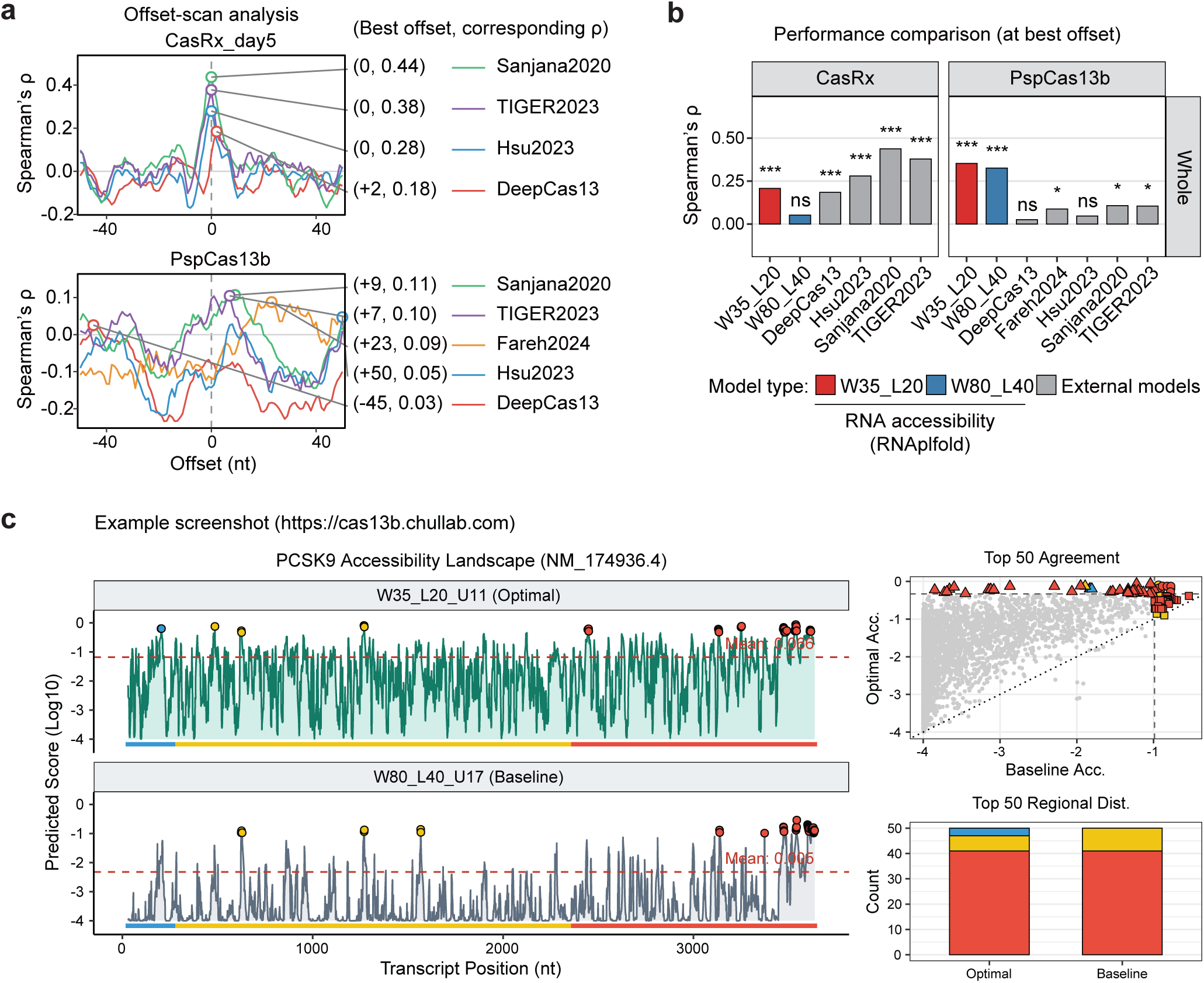
A transcriptome-wide resource for PspCas13b putative guide design. **a** Offset-scan analysis assessing the concordance between seven external prediction models and experimental CasRx (top) and PspCas13b (bottom) log2FC. Spearman correlation is plotted as a function of the positional offset (in nucleotides) applied to the model predictions relative to the experimental data. Colored lines distinguish individual models; filled circles mark the best offset for each model, with the optimal offset and corresponding r annotated on the right via leader lines. **b** Performance comparison between our parameter sets (W35_L20 and W80_L40, colored) and seven external prediction models (grey) across the ‘whole’ transcript region for CasRx (left) and PspCas13b (right). Bar height represents Spearman correlation between model predictions and experimental log2FC. Significance labels reflect raw Spearman P values (*P < 0.05, **P < 0.01, ***P < 0.001). External models include: Sanjana2020 (23 nt and 30 nt), TIGER2023 (23 nt), DeepCas13 (23 nt and 30 nt), Hsu2023 (30 nt), and Fareh2024 (30 nt). **c** Example screenshot showing the PspCas13b guide accessibility landscape for PCSK9. Left: track plot showing the log10-transformed accessibility score for all 30-nt PspCas13b target windows under both parameter sets (W35_L20_U11, green; W80_L40_U17, grey). The colored rug plot along the bottom marks transcript regions (5′ UTR, blue; CDS, yellow; 3′ UTR, red). Points indicate the top 50 ranked guides. Dashed red lines mark the transcript-wide mean accessibility for each parameter set. Upper-right: scatter plot comparing log10 accessibility scores between the two parameter sets, with points colored by region and shaped by shared or parameter-specific Top-50 membership. The diagonal identity line and Top-50 cut-off thresholds (dashed grey) are indicated. Lower-right: regional distribution of the top 50 ranked guides under each parameter set.

